# Enhancing enzymatic bioconjugation efficiency via installation of a substrate recruitment domain

**DOI:** 10.1101/2025.10.27.684804

**Authors:** Christopher Shelby, Kaylee P. Kuzelka, Jonathan M Ellis, Zhiyuan Yao, Amelia C. McCue, Rodney Park, Satish K. Nair, Albert Bowers, Brian Kuhlman

**Author notes:** **Corresponding Author Brian Kuhlman** - Department of Biochemistry and Biophysics, University of North Carolina, Chapel Hill, North Carolina, 27599, United States; Lineberger Comprehensive Cancer Center, University of North Carolina, Chapel Hill, North Carolina, 27599, United States.

## Abstract

Enzyme mediated bioconjugation provides a method for easy and rapid formation of protein-protein and protein-small molecule conjugates under mild conditions. Promiscuous enzymes are of particular interest because they can catalyze conjugation reactions on a broad set of substrates. However, this promiscuity carries the risk of undesirable off-target modifications. To mitigate this effect, we used computational design to install a substrate recruitment domain (SRD) onto the promiscuous enzyme, tyrosinase. The redesigned tyrosinase, called D42, preferentially modifies tyrosine residues within the peptide core (core) linked to a 6-amino acid recognition motif/sequence (RS) specific for the SRD. Incorporation of the recognition sequence along with a neighboring tyrosine in peptides or proteins allows for rapid D42-mediated conversion of the tyrosine to an orthoquinone, which can be selectively modified with a variety of nucleophiles. We demonstrate the utility of our design system by rapidly installing cytotoxic molecules on a monoclonal antibody.

**For Table of Contents Only:** 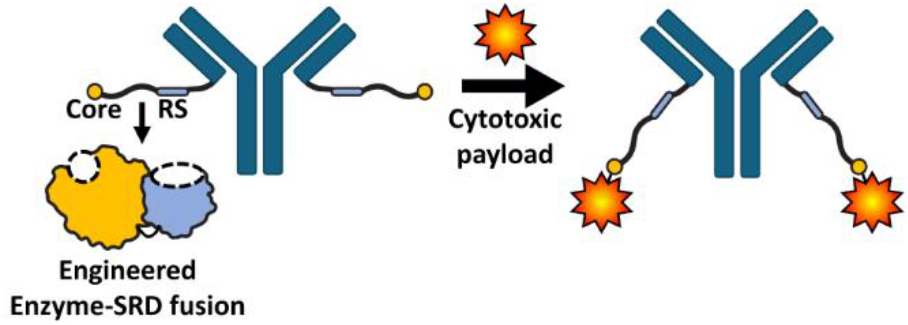

## INTRODUCTION

Many protein and peptide post translational modification (PTM) strategies have been developed to study biological systems and create new therapeutics, with notable examples including biotin/streptavidin, fluorophore labeling, and antibody-drug conjugates (ADCs) for anticancer therapy. While many of these PTMs can be generated without the use of enzymes, these strategies often require harsh conditions that can damage the target biomolecule. Additionally, some chemistry based PTM methods require lengthy and laborious reaction cascades, consisting of multiple protection, de-protection, wash, extraction, and purification steps that can reduce overall sample yields. Enzymes overcome these shortcomings by allowing for direct, rapid, and selective PTM generation under mild conditions^1^.

Tyrosinase is a copper dependent tyrosine oxidase that is found in all domains of life. Tyrosinase is the primary driver in the melanin synthesis pathway, catalyzing the first few critical steps in tyrosine oxidation: first hydroxylating tyrosine at the 3’ position to form the catechol, L-DOPA; followed by oxidizing L-DOPA to the dione, resulting in a highly reactive orthoquinone electrophile (Figure 1A)^2,3,4,5^. In recent years, the bacterial tyrosinase from *Bacillus megaterium* (bmTyr) and the mushroom tyrosinase from *Agaricus bisporus* (abTyr) have been re-purposed to install orthoquinone functional handles that are accessible to nucleophilic attack by thiols^6,7,8,9,10,11,12,13^, primary amines^11,14^, prolines^15,16^, histidines^11^, and strained alkynes^9,17^ to facilitate protein-protein, protein-peptide and protein-small molecule conjugation.

**Figure 1.**
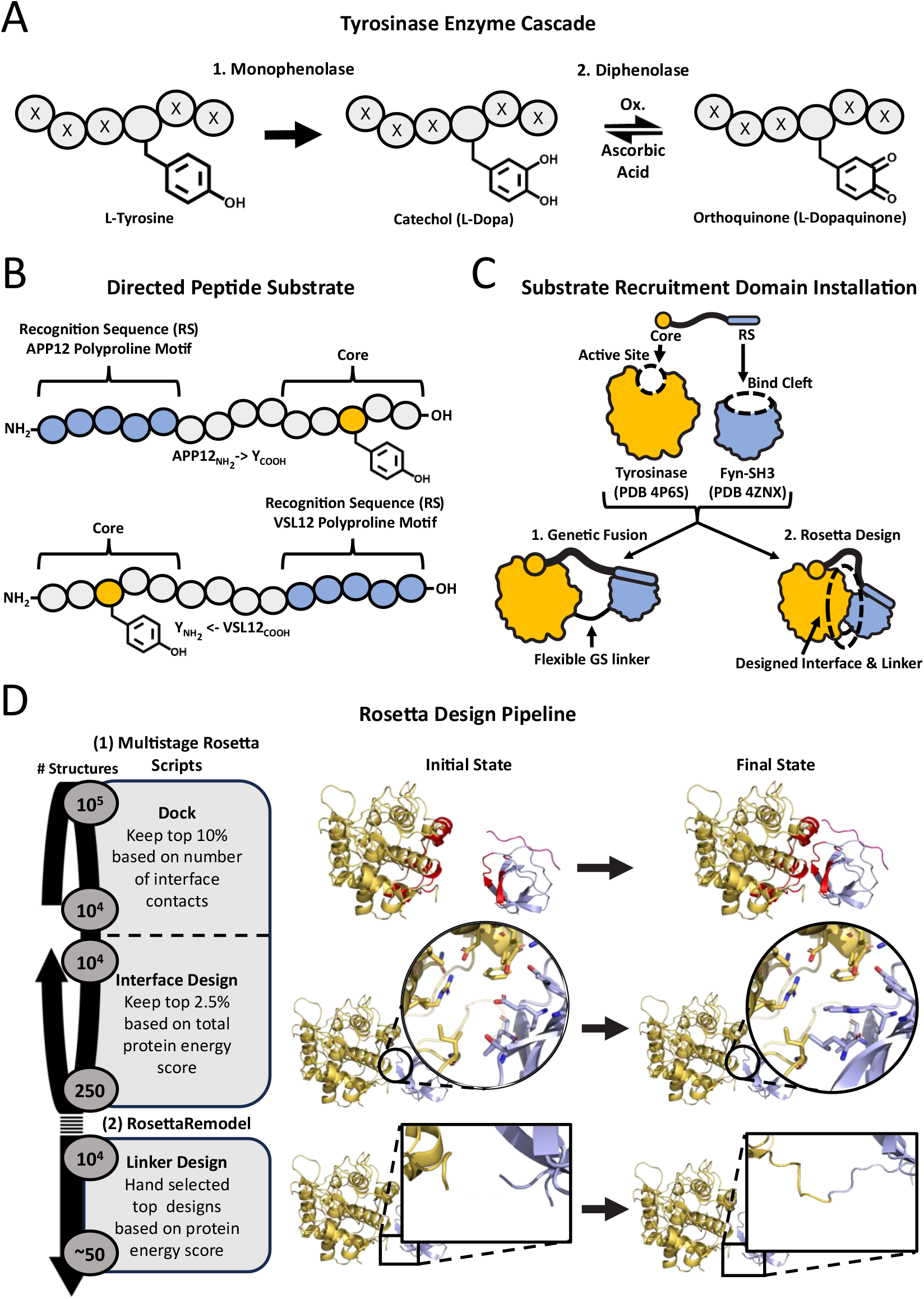
General strategy for designing substrates and SRD fusion proteins. **A**. Diagram of tyrosinase’s sequence promiscuous activity on L-tyrosine to form L-dopa and L-dopaquinone and non-enzymatic reduction of L-dopaquinone back to L-dopa via ascorbic acid. **B**. Substrate construct models depicting two possible peptide construct options. One construct consists of an N-terminal APP12 recognition sequence (RS) for directing a C-terminal core motif (core) to the enzymes active site. The second construct consists of a C-terminal VSL12 recognition sequence (RS) for directing a n-terminal core motif (core) to the enzymes active site. The RS and core motifs are separated by an appropriate length glycine-serine linker. **C**. Diagram depicting mode of targeted substate engagement, proteins selected, and design approaches used for installing a substrate recruitment domain onto tyrosinase. **D**. Overview of Multistage Rosetta scripts pipeline for engineering the fusion proteins.

Both bmTyr and abTyr were shown to install these reactive orthoquinone functional handles in a largely sequence-agnostic manner. However, these enzymes display varying degrees of sensitivity to steric and electrostatic effects of neighboring residues. Specifically, abTyr was shown to be significantly less reactive towards internal peptidyl-tyrosines, yielding only 62% modification against a KETYSK peptide as opposed to 97% for the KNFLDY peptide^1^. abTyr was also shown to have little to no activity against phenol-ssDNA or peptidyl-tyrosine substrates with net negative charges, likely due to repulsion of the largely negatively-charged surface of abTyr^18^. Conversely, bmTyr was shown to have reduced activity on tyrosines with positively charged neighboring amino acids because of the electropositive surface charge of bmTyr. This substrate charge bias was even greater for the bmTYR D55R mutant, which showed less activity towards tyrosines near positively charged amino acids. The charge biases of the D55R variant in abTyr or bmTyr was leveraged to direct these enzymes to basic and acidic terminal tyrosine tags respectively, allowing for tandem, charged directed multiprotein complex formation^18^. Using this approach, investigators were able to tune tyrosinase’s substrate affinity at the expense of the enzyme’s substrate scope^18^.

Here, we explore an alternative approach for tuning tyrosinase substrate preferences while simultaneously enhancing activity towards sterically and electrostatically encumbered substrates. Using computational tools for protein interface design, we installed a substrate recruitment domain (SRD) on tyrosinase, so that substrates containing amino acid sequences recognized by the SRD are directed towards the enzyme active site. In nature, there are numerous examples of enzymes leveraging SRDs to improve their enzymatic efficiencies and substrate selectivity. For example, ring/U-box E3 mediated ubiquitination relies largely on E3 both stabilizing the closed/reactive state of ubiquinated-E2 while also colocalizing ubiquitinated-E2 and the target substrate for efficient ubiquitin transfer^19,20^. Another notable example is the SRD activity of PPP1R15A within the holophosphatase trimer (PPP1R15A, G-actin, and protein phosphatase 1 (PPIc))^21^. The holophosphatase trimer was shown to improve PPIc’s enzyme efficiency 50-fold over PPIc alone^22,^. Reactions involving ribosomal synthesized and post translationally modified peptides (RiPPs) are acted on by enzymes that leverage SRDs to increase the local concentration of their substrates^23^. The SRD recognizes and binds the RiPP’s N-terminal leader sequence and extends the peptides C-terminal core motif to the adjoining catalytic domain^24,25^. In many cases the N-terminal leader sequence is cleaved off to reveal a peptide consisting of only the modified core substrate^25^. This mechanism allows the enzyme to focus its activity on a particular sub-sequence within a complex macromolecular environment while preserving enzyme promiscuous activity towards diverse core sequences.

Previously, we developed and validated a Rosetta design pipeline to engineer enzymes containing covalently linked SRDs, and demonstrated that we can enhance both the catalytic efficiency and selectivity of the enzyme COMT^26^. Here, we apply this approach to a mutated variant of tyrosinase (R209H) from *Bacillus megaterium* (referred to as bmTyr* in this paper). The mutation R209H is located near the tyrosinase active site and improves reactivity with positively and negatively charged peptide substrates^18^. For the SRD, we employ a hyper stable variant of the Fyn-Src Homology 3 (Fyn-SH3) domain, which binds the polyproline motif PLPPxR^27,28,29,30,31^. Computational methods for protein interface design from the molecular modeling program Rosetta were used to position the SH3 domain adjacent to the tyrosinase active site in a manner that directs peptide substrates towards the enzyme active site (Figure 1B,1C). We show that the designed enzymes rapidly modify tyrosine residues positioned adjacent to polyproline motifs in a peptide substrate, allowing for rapid bioconjugation of proteins or peptides that are engineered to include a tyrosine and neighboring polyproline motif.

## RESULTS AND DISCUSSION

### Rosetta Design of Tyrosinase with an SRD

Crystal structures of bmTyr (PDB 4P6S) and Fyn-SH3 (PDB 4ZNX) served as the starting points for computational design^32^,^33^. Fyn-SH3 was docked against the surface of bmTyr and residues on the surface of bmTyr and Fyn-SH3 were mutated to stabilize the interaction. Docking proceeded in two stages: manual docking via Pymol to place Fyn-SH3 adjacent to the bmTyr active site followed by energy-based refinement with Rosetta. Fyn-SH3 was positioned to ensure that the interface was composed largely of residues on the surface of select secondary structure elements. Specifically, we docked Fyn-SH3’s beta barrel against helical segments on tyrosinase, then adjusted the relative orientation of the proteins so that the C-terminus of peptides bound to FYN-SH3 would be directed towards the active site of bmTYR (Figure 1D). In addition, we included a distance constraint between the C-terminus of bmTyr and the N-terminus of Fyn-SH3 to ensure that they would be in proximity to one another. This reduced the length of linkers needed to fuse the two proteins. Rosetta uses a stochastic Monte Carlo algorithm to dock proteins, so to increase the diversity and quality of the docked complexes, we performed ∼10,000 rigid body docking simulations (Figure 1D). We then performed two-sided interface design, wherein select residues at the protein-protein interface underwent iterative rounds of sequence optimization and side chain rotamer sampling to build new and favorable hydrophobic and hydrophilic interactions between bmTyr and Fyn-SH3. Finally, RosettaRemodel was used to design a linker between the proteins. During this design process, designs were sorted and filtered using calculated energies and desired structural features to enrich for designs with numerous and favorable interface contacts (see methods section).

We sorted the final designs by their energies calculated with Rosetta, and hand selected 54 from the top scoring designs for experimental validation. Structural features that were considered during this process included proximity of bound peptides to the bmTyr active site, linker length required for fusing bmTyr and FYN-SH3, number of buried hydrophobic residues at the protein-protein interface, and presence of both hydrophobic and hydrophilic interface residue contacts.

### Initial Experimental Screening

Of the 54 designs, 50 constructs expressed and were soluble as fusions with a N-terminal His tag and the small protein SUMO. Of these, 49 showed activity against free tyrosine as determined by incubating enzyme from crude cell lysates with free tyrosine and observing the visible color change from clear to red during dopachrome formation. This color change was not observed for untransformed crude cell lysate. Each of the 49 designed fusions were then purified using nickel affinity column chromatography and gel filtration. At the same time, we constructed and expressed a naïve fusion of bmTyr and Fyn-SH3 by linking the two domains with a short glycine/serine linker (referred to herein as NF). It is expected that NF will bind and recruit proline containing peptides, but the peptide termini will not be strictly constrained to point at the enzyme active site.

Each enzyme was individually assayed with the directed peptide, APPLPPRNRPRLGSGSGSGSKNFLDY (peptide 1), as substrate. Peptide 1 is composed of an N-terminal APP12 polyproline motif (APPLPPR)^27^,^31^, a C-terminal core motif containing a terminal tyrosine, and a 4x glycine-serine linker to allow the tyrosine of the bound peptide to reach the enzymes active site (Table S1). These reactions were run in the presence of ascorbic acid to trap the L-DOPA product for easy analysis via MS. Enzymatic activity was measured as the fraction of peptide converted to the L-DOPA product as determined by MALDI-TOF. We identified 6 designs that converted substrate to product more quickly than the parent enzyme bmTyr* at substrate concentrations of 1 μM and 10 μM (Figure S1, Table S2 and S3). Design 42 (D42) exhibited the highest activity at both substrate concentrations.

Interestingly, design D8 performed similarly to D42 at a substrate concentration of 10 μM but was catalytically slower than D42 at a substrate concentration of 1 μM. To better understand this difference, we used fluorescence polarization measurements to determine the affinity of selected designs for a peptide containing the APP12 polyproline motif and a N-terminal TAMRA label. As all the designed proteins contain the same FYN-SH3 domain we expected them to have similar affinity for the peptide. NF, D42, and D40 all bound to the peptide with similar affinities (K_d_ < 2 μM), while D8 bound weaker to the peptide (K_d_ = 18.5 μM) (Figure 2A). This result is consistent with the activity assay and suggests that the peptide-binding site in D8 is obstructed or distorted. As expected, bmTyr* did not show affinity for the APP12 recognition sequence at the concentrations tested. Based on the initial activity and binding measurements, D42 was chosen for further biophysical characterization and applications.

**Figure 2.**
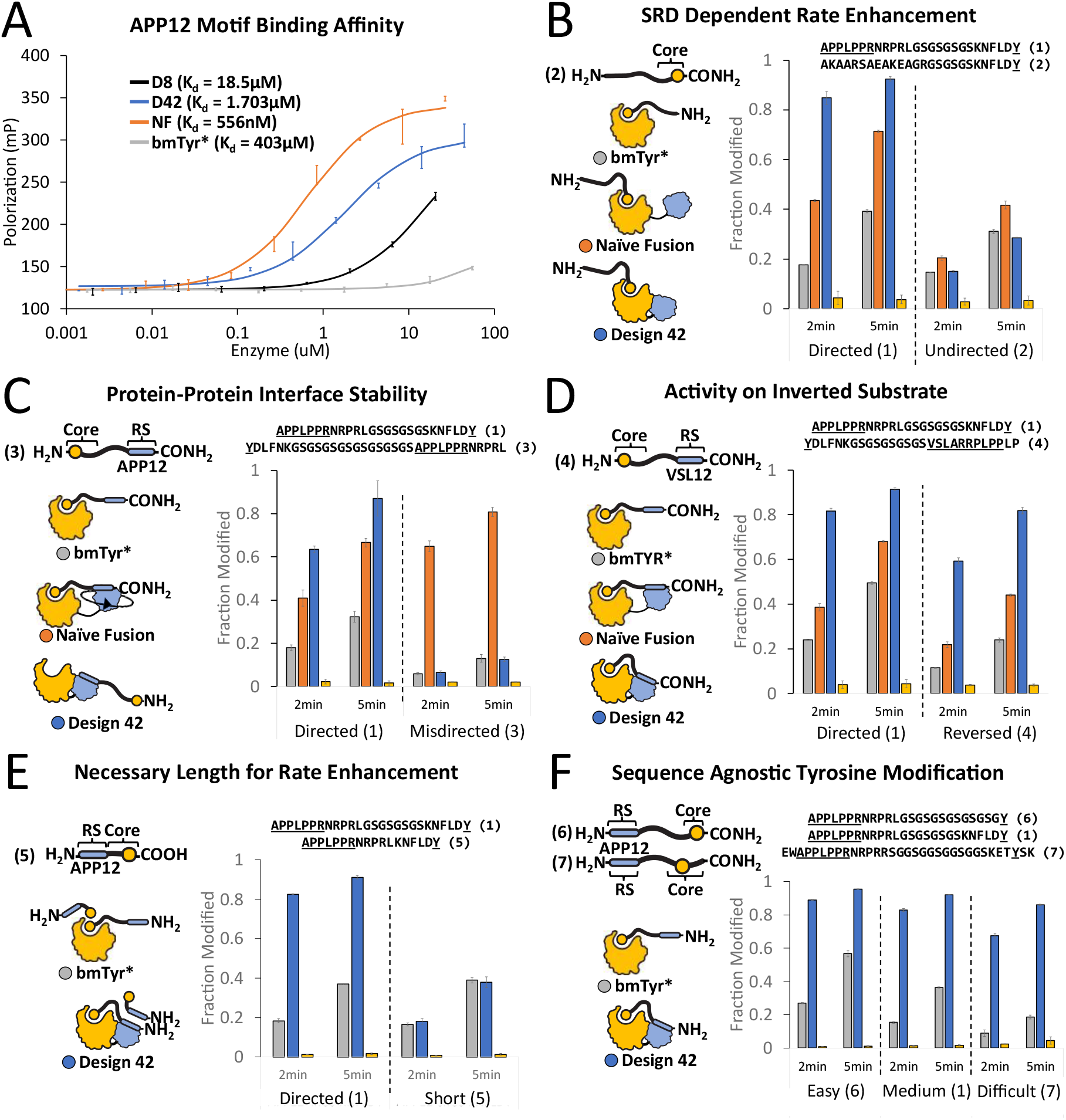
Biophysical and Biochemical Characterization of Rosetta Design 42. **A**. Fluorescence polarization measurements with a N-terminal TAMRA labeled APP12 peptide fitted to a first order binding curve showing K_d_s for bmTyr*, NF and D42. **B.-F**. LC-ESI-TOF EIC ratios of product [hydroxylated] over sum of product and substrate peptide EICs. All reactions were performed at 10μM peptide and 50nM enzyme in the presence of ascorbic acid and quenched with 4mM tropolone. The error bars represent standard deviation across three replicates centered about the mean.

### Biophysical and Biochemical Characterization of D42

To verify that the enhanced reaction rates observed with D42 are due to substrate recruitment, we performed activity assays with a peptide lacking the polyproline motif, AKAARSAEAKEAGRGSGSGSKNFLDY (peptide 2). The reactions were again performed in the presence of 12.5 μM ascorbic acid to trap the L-DOPA product. Reactions were quenched at 2 minutes and 5 minutes using 4mM tropolone and analyzed using LCMS. Both peptides 1 and 2 were tested with three enzymes, bmTyr*, NF and D42. With the undirected peptide, all three enzymes converted similar amounts of substrate to product (<20% in 2 minutes) (Figure 2B and S2). In contrast, with the directed peptide, D42 converted 84% of the substrate to product within 2 minutes while NF converted 44%, and bmTYR* converted less then 20%. These results suggest that the enhanced reaction rates observed for D42 is SRD dependent, and that the designs basal enzymatic activity has not been attenuated by installing a SRD.

In the design model of D42, the C-terminus of the APP12 peptide is directed towards the active site of tyrosinase. If D42 folds as intended, substrates with a tyrosine located C-terminal to the polyproline motif should be poised for modification while substrates with a tyrosine located N-terminal to the polyproline motif should not benefit from substrate recruitment. To test this hypothesis, activities of bmTyr*, NF, and D42 were tested with either a C-terminal tyrosine (peptide 1) or N-terminal tyrosine (misdirected peptide, peptide 3). Consistent with the design model, D42 exhibited rate enhancement with peptide 1, but no rate enhancement was observed with the misdirected peptide (Figure 2C and S3). NF enhanced reaction rates with both substrates, as would be expected if the SH3 domain is able to move freely relative to the catalytic domain. This result highlights the utility of designing a rigid interface between the substrate recognition domain and catalytic domain, as it affords more control over the site of modification in the substrate.

One interesting feature of FYN-SH3 is that polyproline sequences have been discovered that bind to this domain in opposite orientations. For instance, the VSL12 polyproline motif binds in the opposite direction as the APP12 motif^27,32,34,35^. In the context of D42, this means that if the VSL12 motif is incorporated into a peptide substrate, then the N-terminus of the peptide should be directed towards the active site (whereas with the APP12 motif the C-terminus is pointed towards the active site). To explore this capability, we generated an inverted substrate containing an N-terminal substrate motif, C-terminal high affinity VSL12 recognition sequence, and interior GS linker: YDLFNKGSGSGSGSGSVSLARRPLPPLP (peptide 4) (Figure 2D and S4). We tested peptide 4 with D42, NF, and bmTYR*. As with the APP12 peptide (peptide 1), the inverted substrate was most efficiently modified by D42, followed by NF, then bmTyr*. These results show that the VSL12 recognition sequence can enable SRD dependent rate enhancement on N-terminal peptidyl-tyrosine targets.

Next, we sought to establish that the enhanced activity of D42 requires simultaneous engagement of both the peptidyl-tyrosine and the polyproline motif to the enzyme. We synthesized a directed peptide with fewer resides between the APP12 polyproline motif and the C-terminal tyrosine, APPLPPRNRPRLKNFLDY (peptide 5). Based on the D42 design model, this peptide should not be able to simultaneously reach the active site and bind to the SH3 domain. When peptide 5 was used as a substrate with D42 or bmTYR*, the same amount of product was formed with each enzyme (Figure 2E and S5), consistent with a loss of effective tandem substrate recruitment and catalytic engagement.

Previous studies have shown that the reactivity of tyrosinase with tyrosine containing peptides varies depending on the amino acids adjacent to the tyrosine^1,18^. SRDs improve enzyme activity by effectively increasing the local concentration of substrate, potentially allowing modification of less favorable substrates. To test D42s activity against substrates with varied intrinsic favorability, we synthesized two additional peptides. Peptide 6 has small and flexible amino acids adjacent to a C-terminal tyrosine (APPLPPRNRPRLGSGSGSGSGSGSG**Y)**, while peptide 7 places the tyrosine two residues from the C-terminus (EWAPPLPPRNRPRRSGGSGGSGGSGGSKET**Y**SK). We incubated the easy (peptide 6), moderate (peptide 1) or difficult (peptide 7) peptides with bmTYR* and observed that the rates of modification decreased with increasingly challenging tyrosine accessibility (Figure 2F and S6). With peptide 7 and bmTyr*, less than 5% of the substrate was converted to product after 2 minutes. In contrast, D42 was able to quickly modify all three substrates, showing that the increased local concentration of the peptidyl-tyrosine helps to alleviate some of the challenges of working with difficult to access tyrosine positions.

In addition to sterics, the electrostatic environment surrounding a tyrosine was shown to influence tyrosinase’s reactivity^18^. The largely charge agnostic enzyme, bmTyr* was shown to have reduced activity against a peptidyl tyrosine containing four neighboring glutamate residues (EEEEY)^18^. To test if SRD can bolster enzyme activity against tyrosines within unfavorable electrostatic environments, we synthesized a peptide containing a APP12 motif, GS linker and a core motif consisting of the four glutamate residues and a C-terminal tyrosine, APPLPPRNRPRLGSGSGSGSG<u>EEEEY</u> (peptide 8) Incubating bmTyr* with peptide 8 resulted in little tyrosine hydroxylation, reaching only 12% after 5 minutes. Conversely, D42 showed enhanced reactivity, reaching 58% conversion over the same time interval (Figure S7). These results demonstrate that SRD can also improve enzyme activity against tyrosines in unfavorable electrostatic environments.

To test the preference of D42, bmTyr* and NF for a target substrate in the presence of a competing substrate, reactions were performed with a mixture of the directed (peptide 1) and undirected peptide (peptide 2). As expected, bmTyr* did not discriminate between the directed and undirected peptides at both substrate concentrations that were tested (1 μM and 10 μM). NF and D42 showed a strong preference for the directed substrate at both 1μM and 10μM (Figure 3 and S8 and S9).

**Figure 3.**
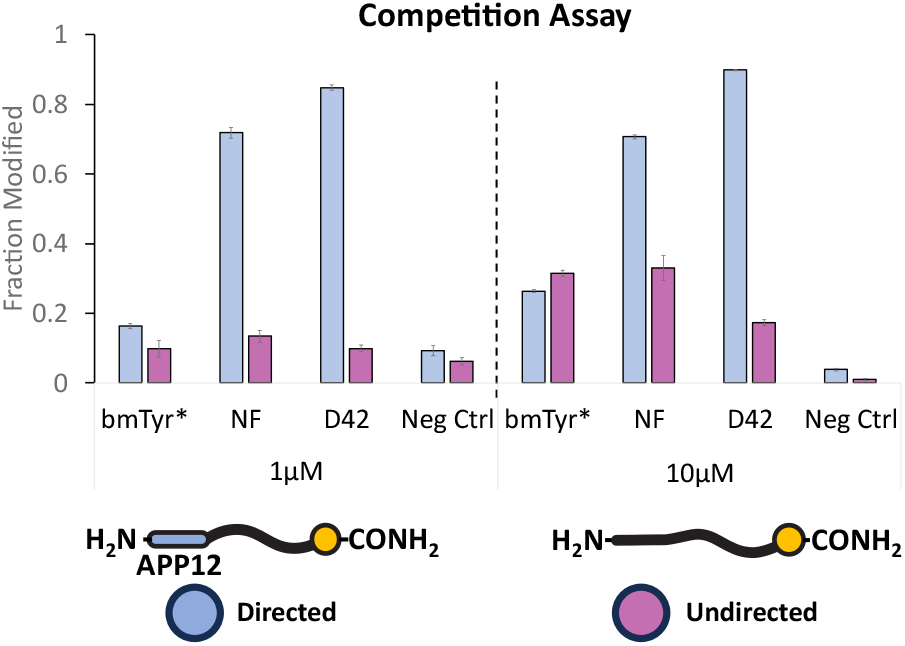
Competition Assay. LC-ESI-TOF EIC ratios of product formed during competition reactions of directed vs undirected peptide at 1μM and 10μM (2μM and 20μM total peptide) when incubated with 10nM and 50nM enzyme, respectively. Each reaction ran for 10min in the presence of 12.5mM ascorbic acid and was quenched using 4mM tropolone. Error bars represent standard deviation across three replicates centered about the mean.

### Crystal structure of D42

To probe the structural accuracy of the computational design, we solved the 2.6 Å resolution crystal structure of D42 in the absence of substrate (Figure 4A, Table S4). In the crystal structure, the SH3 domain interacts with the surface of bmTyr with its peptide binding groove on the desired side of the protein (Figure 4A). The APP12 polyproline motif was superimposed onto the structure and, as intended, its C-terminus points towards the active site (Figure 4B). Modeling peptide 1 onto the crystal structure shows that the C-terminal tyrosine can reach the active site, while modeling with peptide 5 (the short peptide) shows that it should not be able to reach the active site, consistent with the observed loss of activity. Like the design model, the interface between the two proteins is formed between two helices on tyrosinase (residues Thr160-Gln172 and Arg232-His241, numbering derived from PDB 9PGZ) and β-strands on the surface of the SH3 domain (residues Thr334-Tyr340). Despite docking with the desired orientation, the precise contacts formed at the interface are different in the design model and the crystal structure. In the crystal structure, the SH3 domain is shifted several angstroms relative to the SH3 domain in the design model (Figure 4A), and the interface is smaller in the crystal structure than the design model. One explanation for these perturbations could be crystal packing (Figure S10A). In the crystal, residues on the surface of an adjacent molecule of D42 pack against the designed SH3/tyrosinase interface, forming hydrophobic (Figure S10B) and electrostatic interactions (Figure S10C) enabled by the shifted position of the SH3 domain. Whether it is due to crystal packing or other energetic factors, the small size of the SH3/tyrosinase interface (∼600 Å^2^) and the shift in SH3 positioning suggests that designed interface is not highly stable. This highlights that weak interactions are probably sufficient in many cases to orient domain-domain interactions, as the substrate preferences exhibited by D42 are consistent with the design model and crystal structure.

**Figure 4.**
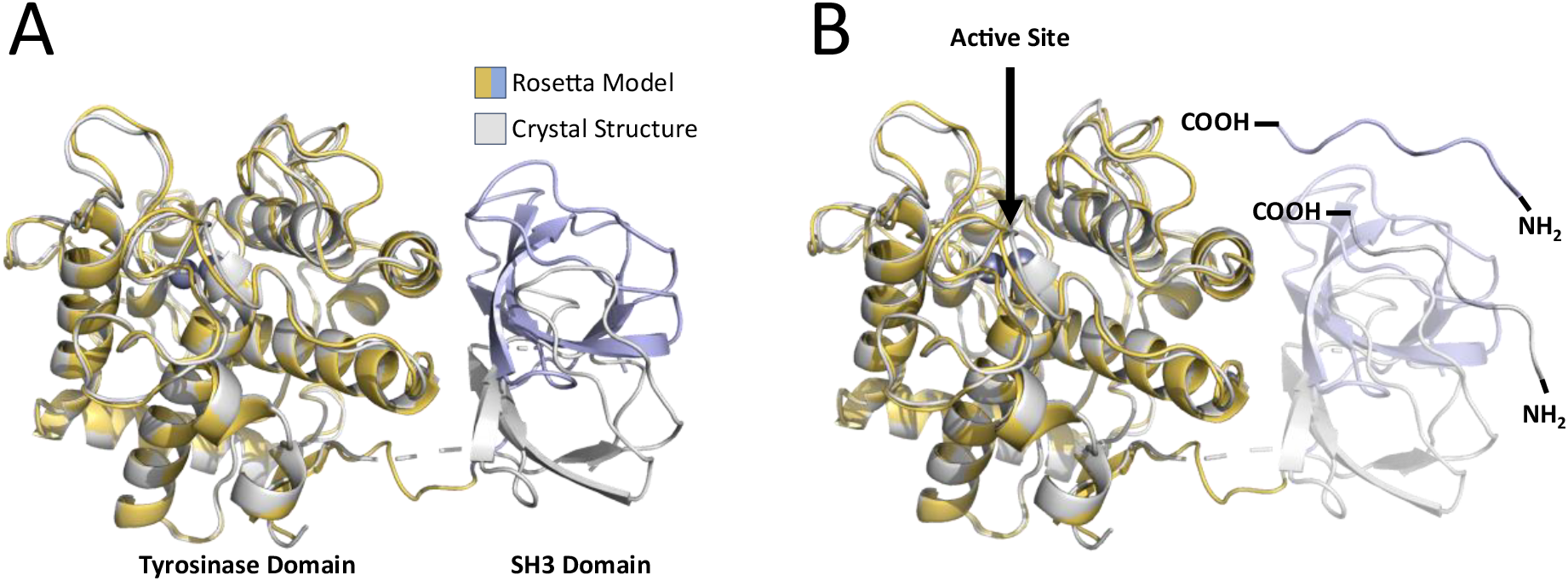
X-ray crystal structure of D42. **A**. X-ray **c**rystal structure of D42 [PDB 9PGZ] (grey) aligned to Rosetta model of D42 (yellow and light blue). **B** Peptide directionality of Rosetta model and crystal structure showing that both structures appropriately positions peptide C-terminus in the direction of D42 active site.

### Antibody Drug conjugation

To demonstrate the utility of SRD-fusion biocatalysts, we chose to employ D42 in the task of creating an antibody-drug conjugate (ADCs), using the formed orthoquinone electrophile as a targetable conjugation moiety. As a first step towards antibody conjugation, we tested D42s ability to install cytotoxic drug molecules onto the directed peptide (peptide 1). For these reactions, we selected two microtubule inhibitor drug molecules that have been used in several FDA approved ADCs: endo-BCN-Val-cit-PAB-monomethyl auristatin E (endo-BCN-PAB-MMAE), and Mertansine (DM1) (Figure 5A). We incubated 50μM of either endo-BCN-PAB-MMAE or DM1 with 10μM of directed peptide (peptide 1) and 250nM enzyme (D42 or bmTyr*) at pH 6.5 and 25°C. D42 rapidly conjugated DM1 to the directed substrate, reaching 76% completion in only 2 minutes. In contrast, bmTyr* took 15 minutes to reach ∼20% completion (Figure 5B). The conjugation of endo-BCN-PAB-MMAE was slower for both enzymes, yielding 40% completion after 2 minutes with D42 and no observed conjugation with bmTyr* at all times tested. We also explored conjugation efficiency for the sterically challenging internal tyrosine peptide (peptide 7) and observed that, as in the case of our monophenolase experiments, bmTyr* struggled to conjugate DM1 to the internal tyrosine peptide, while D42 reached 100% within 15 minutes (Figure S12). Overall, these results show that D42 is significantly more efficient at generating orthoquinone functional handles and facilitating drug conjugation than bmTyr*, and suggest that, in addition to improving enzyme activity for single step reactions, SRDs can dramatically accelerate multi-step enzyme reactions.

**Figure 5.**
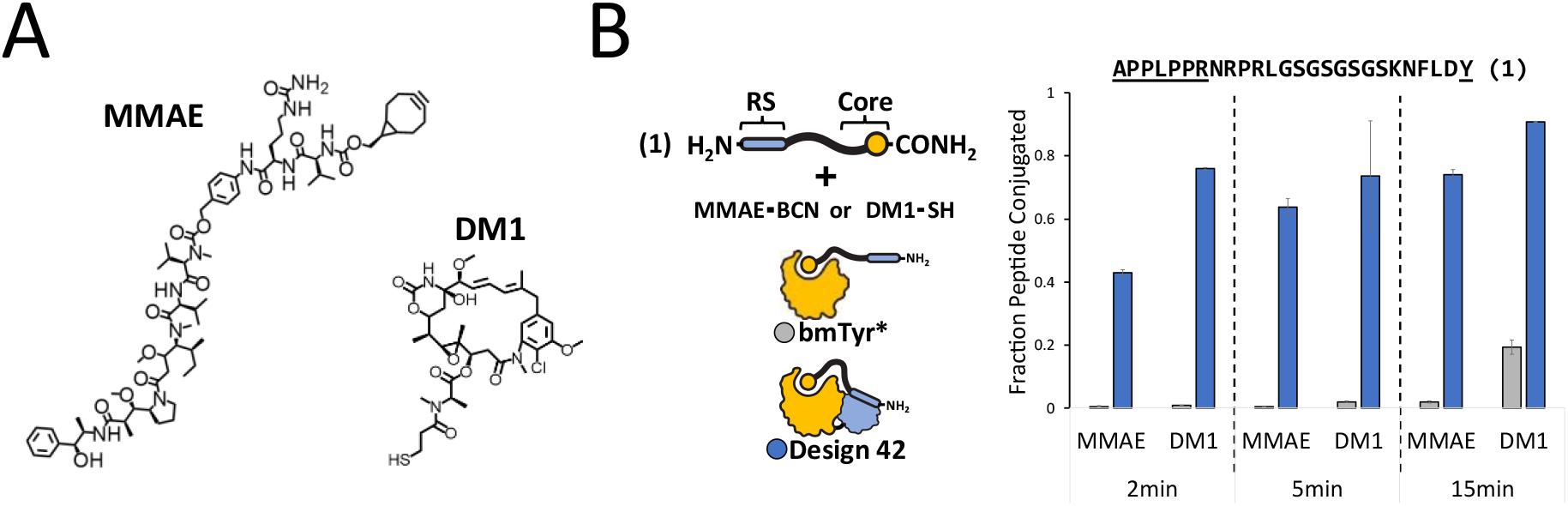
Installing cytotoxic drug compounds onto peptides. **A**. Chemical structures for cytotoxic drug compounds, endo-BCN-Val-cit-PAB-monomethyl auristatin E (MMAE) and Mertansine (DM1), used in this bioconjugation reaction. **B**. Fraction of directed peptide (peptide 1) conjugated to drug compound at various time points. Reactions ran at 10μM peptide, 50μM drug compound and 250nM enzyme. Error bars represent standard deviation across three replicates centered about the mean.

We next evaluated D42’s ability to conjugate DM1 to Rituximab: an anti-CD20 antibody used to treat leukemia and lymphoma^36,37,38,39,40^. To facilitate substrate recruitment with D42, we modified the C-terminus of Ritiximab’s light chain to include a APP12 polyproline motif, a PreScission protease cleavage site for payload release, and a (KGSGSGYAAA) core motif (herein referred to as Rit-APP12-YA3) (Figure 6A). The core motif KGSGSGYAAA was chosen as opposed to KNFLDY to avoid problems with tyrosine sulfation during mammalian expression^41,42^. We incubated 5μM (10μM C-terminal tag equivalents) Rit-APP12-YA3 with 50μM DM1 and 250nM enzyme (D42 or bmTyr*) at pH 6.5 and 25°C. After 10 minutes, reactions were quenched using 4mM tropolone. All reactions were reduced with 10 mM DTT before submitting them for analysis via intact MS. Like the results with the directed peptide, the ADC yield was ∼75% after 10 minutes with D42. We saw only a small amount of ADC generated with bmTyr* (Figure 6B-D). We also explored the potential for the proteolytic release of the DM1 cytotoxic payload. Protease sensitive ADCs are being used in the clinic to promote cytosolic uptake^43,44,45^. As before, we reacted the engineered Rit-APP12-YA3 with bmTyr* or D42 for 10 minutes, quenched with tropolone, and exposed the antibody to PreScission protease for at least 15 minutes at 30°C (Figure 6E-G). We observed complete release of the ADC cytotoxic payload. These results show that D42 selectively modified the engineered tyrosine on the light chain and demonstrates protease-dependent payload release. The protease/cleavage site could be swapped for a different motif recognized by a protease that is significantly upregulated in cancerous cells or abundant within the tumor microenvironment for targeted drug delivery.

**Figure 6.**
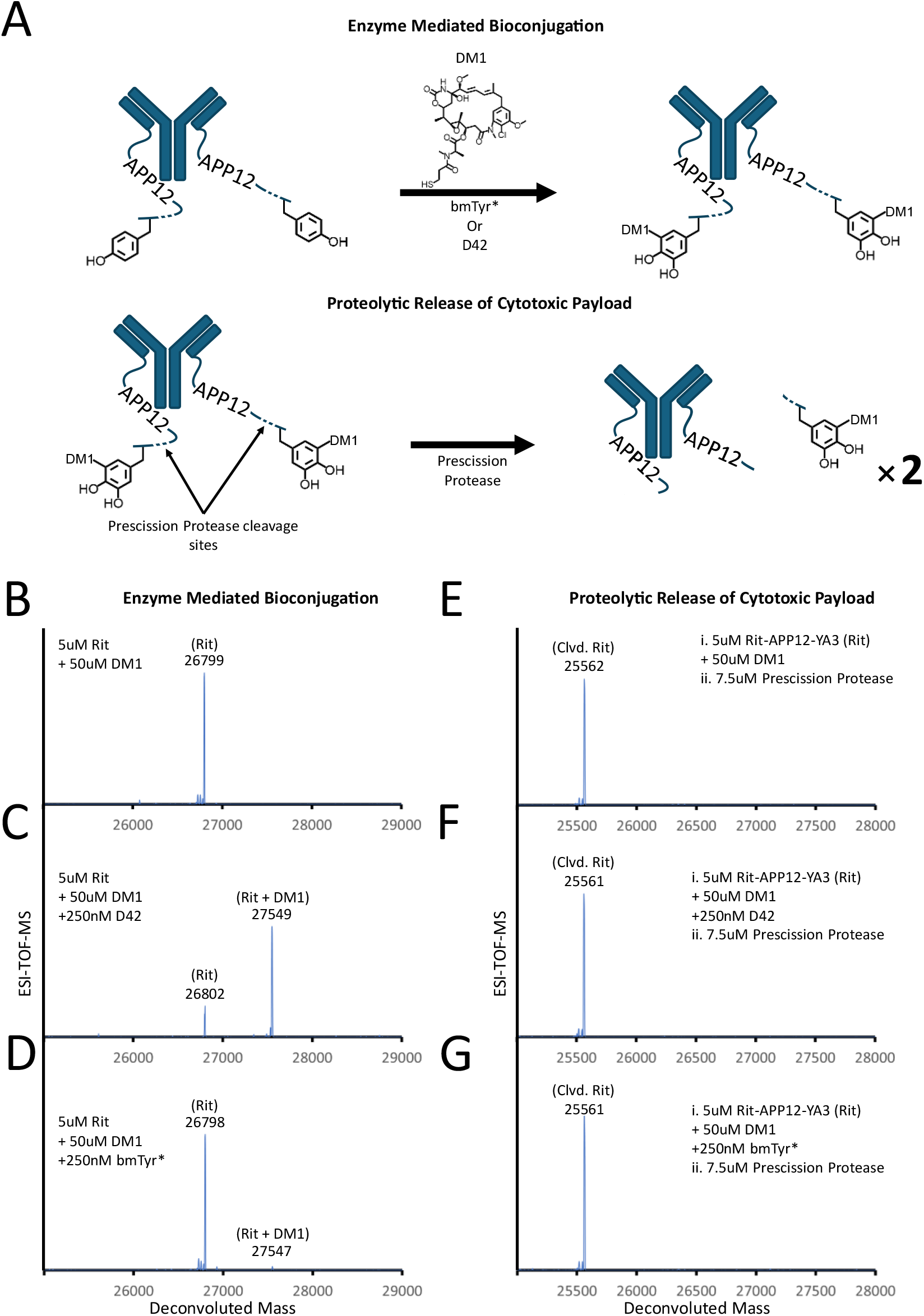
Generating antibody drug conjugates. **A**. Conjugation strategy for installing DM1 onto antibody using bmTyr* or D42 and subsequent proteolytic cleavage of modification tag to validate primary location of modification. **B-D** Conjugation of DM1 to rituximab using **B**. no enzyme, **C**. D42, or **D**. bmTyr*. Reactions were performed for 10min at 25°C before quenching with 4mM Tropolone. **E-G**. After incubating antibody and DM1 with either no enzyme, D42, or bmTyr* for 10min and quenching with 4mM tropolone, each reaction was incubated with Prescission Protease at 30°C for 15 min to cleave off core motif conjugated DM1 payload. Antibody was reduced with 10mM DTT prior to running intact MS.

We performed an additional ADC experiment over longer time intervals to drive the conjugation reaction to completion. We observed D42 produced 100% ADC yield after 15 minutes (Figure S13C), while bmTYR* only produced ADC yields of ∼15% and ∼30% after 15 and 30 minutes respectively (Figure S13D-F). Interestingly, in these longer reactions with D42, we observed an additional peak in the mass spectrometry that corresponds to a second DM1 being conjugated to the target tyrosine. This result is corroborated by an observed mass shift of 738 Da, corresponding to addition of a second DM1 at the original modification site, in contrast to the expected 754 Da mass shift stemming from modification of a second tyrosine residue (16 Da hydroxyl group + 738 Da DM1 molecule. To further validate this result, we used PreScission protease to remove the conjugated peptide from the antibody and performed LC-MS. Masses for both a single and double DM1 incorporation could be observed on the cleaved core motif, confirming that double DM1 conjugation to the same modified tyrosine is indeed the cause (Figure S14). A previous study showed that diphenol rings possess two sites for thiol addition, with the 5’ position being the primary reaction site and a minor site at the 2’ position^10^. Overall, D42 is far more efficient at generating ADC than bmTyr*, and the ability to conjugate multiple DM1 to a single tyrosine could increase toxicity and improve the efficacy of the ADC. Thus, the improved processing enabled by SRD-targeting could allow reliable access to higher drug-antibody ratios from a single tyrosine residue.

## CONCLUSION

Using computational design, we successfully engineered an SRD-enzyme fusion that recruits and orients peptide substrates in a RiPP-like manner. D42 was able to selectively modify the on-target peptide even in a complex reaction environment where another off-target peptidyl-tyrosine was present. Additionally, the presence of a SRD expanded the substrate scope of tyrosinase, as D42 showed improvement in activity on substrates with internal tyrosine residues. When we explored reactions that required the multistep tyrosine oxidation pathway of tyrosinase, D42 displayed a dramatic rate acceleration, allowing for rapid installation of cytotoxic molecules onto our model antibody. Our structural data suggest that computationally designed SRD-enhanced systems do not require high affinity interfaces between the SRD and protein target domains to benefit from an SRD. Only interfaces of sufficient stability to bias the structure of the SRD to ideally position its substrate is needed. This makes engineering such systems readily accessible with the computational design tools available today. An exciting future application for enzymes engineered to include SRDs is the creation of enzyme cascades where each enzyme in the pathway possesses the same SRD, allowing multiple reactions to be performed on a single protein or peptide substrate. Finally, we demonstrated that D42 can be used to install multiple cytotoxic compounds rapidly and selectively onto antibodies engineered to include a polyproline motif and neighboring tyrosine. The ability to genetically encode the antibody-drug linker provides great control over which protease target to use for cytotoxic payload release and the ability to screen many different potential protease targets rapidly and easily during an ADC design campaign.

## SUPPORTING INFORMATION

The Supporting Information is available free of charge at

Additional information about Genes, extended experiments, LCMS traces, and peptide validation (PDF)

## ACKNOWLEDGMENTS

The peptide synthesis was performed in the UNC Peptide Synthesis Core facility (RRID:SCR_017837), work was supported by the National Cancer Institute of the National Institutes of Health under award number P30CA016086. The intact mass spectroscopy was conducted using the UNC Proteomics Core Facility. Work was supported by the National Cancer Institute Center Core Support Grant (award number: 2P30CA016086-45) of the UNC Lineberger Comprehensive Cancer Center. The authors thank Jarrett Pelton for assisting us with running and optimizing our LCMS experiments. The authors also thank Matthew Begley for his donation of HRV 3C plasmids (PreScission protease) and various reagents.

## METHODS

### Enzyme Selection and Initial Dock Generation

Tyrosinase from *Bacillus megaterium* (bmTyr) was selected because if its prior use in bioconjugation reactions, the breadth of enabling bioconjugation chemistries, the availability of high-resolution structures (PDB 4P6S used in this study), and surface-exposed active site. To reduce the charge bias of bmTyr, our enzymes contain the R209H active site mutation and is here referred to as bmTyr*. Fyn-SH3 (PDB 4ZNX) was selected because of its high thermostability^46^, substrate affinity^29^, fast k_on_ and k_off_ rates^28,29^, and use in folding studies^47^,^48^. Initial dock structures were generated by manually positioning the SH3 domain near tyrosinase using Pymol’s drag function (roughly 5Å apart). The SH3 domain was positioned in a manner to mutually complement both tyrosinase and SH3 surface topologies. Dockable and designable residues were selected from large, solvent exposed regions consisting of well-defined secondary structural elements (α-helices on tyrosinase and β-sheet within the β-barrel of SH3) at or near the initial dock structure’s interface. Initial dock structures were pre-relaxed with energy minimization in Rosetta.

### Protein Design

Computationally designed fusion proteins were generated using Rosetta’s multi-stage scripts^26,49,50,51^. Briefly, three separate stages of Rosetta design are implemented with the first stage composed of rigid body docking which brings SH3 into contact distance with tyrosinase and the next two stages consisting of iterative rounds of rotamer and residue optimization to stabilize the newly generated protein-protein interface. The first step is composed of rigid body rotational and translation perturbations of the initial dock PDB pose to bring the two proteins into contact. During this stage, a peptide distance constraint was applied to ensure that the SH3 domain would position its substrate near and in the direction of the tyrosinase domain’s active site. In addition, a termini distance constraint was applied to ensure that only a minimal length linker is needed to covalently link both domains. For each initial dock structure, 10000 docking simulations were performed for each input coordinates and the top overall 10% docked structures based on number of residue contacts were passed on to the second stage. In the second stage, one round of fast design was performed for each docked structure to quickly assess the quality of the docked structures and removal of structures with backbone and sidechain clashes that cannot be rescued by mutating residues or sampling different side chain rotamers. Designs are sorted according to total energy score and the top 25% were passed onto the next stage. During the third stage, two rounds of fast design were implemented to further optimize the amino acid composition and rotamer orientation for the protein-protein interface. The designed structures were again sorted by their total energy score, and only the top 10% are selected for linker design. The entire multi-stage docking, and interface design pipeline was repeated until > 10,000 final designs were generated.

### RosettaRemodel Linker Design

Linkers for each of the 10000 final designs were generated using Rosetta Remodel^52,53,54^. Briefly, 0-10 residues were inserted between tyrosinase and SH3 for all 10000 final interface designs using RosettaRemodel. Linker backbone conformations were then sampled from a non-redundant fragment library derived from structures in the PDB. Immediately after linker backbone generation, a single round of fast design was performed on the residues in the linker. Linker designs with chain breaks (termini that are not covalently linked) and linkers with two or more consecutive residues with positive Rosetta *RAMA* score terms were filtered out. The remaining designs were sorted by total energy score and from the top scoring structures, 57 designs were selected by hand for experimental characterization.

### Bacterial Cell Transformations and Protein Expression

Expression constructs for the final 57 designs were ordered from Twist Biocience using a custom pCDB24_xhol plasmid backbone. These constructs were transformed into heat competent BL21(DE3) pLys cells with 50ng of DNA. Cells were plated onto LB plates containing an ampicillin (100mg/L) selection marker and incubated at 37°C overnight. Colonies were subsequently selected and inoculated into 50mL LB media and grown overnight at 37°C with vigorous shaking at 250 RPM. The following day, 20mL of overgrown LB culture was transferred to 1L of fresh terrific broth (TB) and incubated at 37°C at 230 RPM until an OD_600_ of 1. Protein production was induced using 600μM isopropyl-1-thio-β-D-galactoside (IPTG) and incubated at 16°C and 230rpm overnight. TB cultures were then pelleted at 4°C and 3148g for 30min and stored at −80°C.

### General Protein Purification

Frozen cell pellets were thawed at room temperature. Cells were then resuspended in 90mL of lysis buffer (50mM Tris pH 8, 500mM NaCl, 10mM Imidazole) and sonicated for 8min (5sec on 5 sec at an amplitude of 70%). Cell lysate was then clarified at 4°C and 15000 RPM for 30min. The supernatant was incubated with Ni-NTA at 120 RPM for 1hr. Sample was then loaded onto a 100-200mL column for gravity flow affinity chromatography. The column was then washed with 10 column volumes of high salt buffer (50mM Tris pH 8, 1M NaCl, and 20mM imidazole) and low salt buffer (50mM Tris pH 8, 500mM NaCl, and 20mM imidazole). Protein was eluted in four 3mL fractions with elution buffer (50mM Tris pH 8, 500mM NaCl, and 500mM imidazole). Proteins were further purified and simultaneously buffer exchanged into storage buffer (50mM Tris pH 8, and 500mM NaCl) by size exclusion chromatography on a Superdex S200 prep column. Protein was flash frozen with 10% glycerol at 1-2.5mg/ml and stored at −80°C

### HRV 3C (Prescission Protease) Protein Purification

Frozen cell pellets were thawed at room temperature. Cells were then resuspended in 90mL of lysis buffer (50mM Tris pH 8, 10% glycerol and 5mM DTT) and sonicated for 8min (5sec on 5 sec at an amplitude of 70%). Cell lysate was then clarified at 4°C and 15000 RPM for 30min. The supernatant was incubated with glutathione-agarose at 120 RPM for 1hr. Sample was then loaded onto a 100-200mL column for gravity flow affinity chromatography. The column was washed twice with 10 column volumes of lysis buffer (50mM Tris pH 8, 10% glycerol and 5mM DTT). Protein was eluted in four 3mL fractions with elution buffer (50mM Tris Base pH 8, and 10mM glutathione (reduced)). Proteins were further purified and simultaneously buffer exchanged into storage buffer (25mM Tris pH 7, and 125mM NaCl) by size exclusion chromatography on a Superdex S200 prep column. Protein was flash frozen with 10% glycerol at 1-2.5mg/ml and stored at −80°C

### Crystallization and Structure Determination

Protein samples were purified as described, the poly(His) affinity tag was removed using tobacco etch virus (TEV), and the affinity tag was separated via subtractive chromatography using Ni-NTA. Following size exclusion chromatography on a Superdex S200 column, freshly prepared protein was concentrated to ∼45 mg/mL using an Amicon filter and screened for crystallization using an Art Robbins Gryphon liquid handling system. Initial crystals were identified with NaCl as a precipitant and the final crystallization conditions were 0.5 M NaCl, 0.01 M magnesium chloride hexahydrate, and 0.01 M hexadecyltrimethylammonium bromide. Prior to diffraction analyses, crystals were immersed in crystallization solution supplemented with 25% (v/v) glycerol and vitrified by direct immersion in liquid nitrogen.

Diffraction data were collected at Sector 7 (Macromolecular Crystallography Cornell High Energy Synchrotron Source, MacCHESS) beamline ID 7B2. These data were indexed, scaled, and integrated using the on-site fast_dp auto-processing. Indexing and integration were performed by XDS^55^, followed by Pointless space group determination. Aimless was used to scale the data^56^. Phases were determined using the Phenix suite Phaser-MR program^57^ (top LLG: 3712.8; top TFZ 34.1), utilizing PDB: 4P6S and 4ZNX as search probes. The initial model was built with Phenix’s AutoBuild program, followed by additional rounds of refinement using Phenix Refine. Coot^58^ was used for manual model building, and further refinements were carried out using REFMAC5^59^ as implemented in the CCP4 suite^60^.

### Modification-Tagged Rituximab Expression and Purification

Rit-APP12-YA3 heavy chain (HC) and modification tagged light chain (LC) were cloned into two separate pAH plasmid backbones. Both HC and LC constructs were co-transfected at 100ug DNA (50ug per construct) into 100mL Expi293 cells at ∼3 x 10^6^ cells/mL using the Expi293FExpression System (Thermo Fisher, #14635) with a human serum albumin signal peptide for secretion to the culture medium. EXPI293F cells were maintained at 37°C with 8% CO2 at 250 rpm without antibiotics. Enhancers were added according to the manufacture’s protocol 20 hours after transfection. The cell cultures were harvested 96 hours after transfection and the supernatants were separated from the cells by spinning at 3000 RCF (4°C) and passing through 0.4µm filters. The supernatants were then purified with protein A resin (GOLDBIO cat#: P-400-25). 1mL of protein A resin was used in the purification and were mixed with 100ml supernatants under gentle shaking at 4 °C for 30 minutes to allow protein binding. After incubation, resin was washed with 5ml of water and 10ml of PBS (PH 7,4). Antibodies were eluted using 1.25mL of 100mM glycine buffer (pH 2.5) and immediately mixed with 125uL of 1M Tris buffer (pH 9.0) to balance PH. Fresh eluted antibodies were buffer exchanged into PBS pH 7.4 and flash frozen with 10% glycerol and stored at −80°C in aliquots for future use.

### Peptide Synthesis

Peptides were synthesized at the High-Throughput Peptide Synthesis and Array Facility at the University of North Carolina using a standard Fmoc solid phase peptide synthesis technique. All peptides were purified to >90% as determined by high-performance chromatography and mass validated using matrix assisted laser desorption ionization (MALDI) mass spectrometry. All peptides including purities and masses are available in Supplementary (Figures S14-S20).

### Fluorescence Polarization

Enzymes were prepared using the previously described protein expression and purification method except that proteins were simultaneously purified, and buffer exchanged via size exclusion chromatography into phosphate buffer (50mM pH 8) and stored in 10% glycerol at −80°C. TAMRA-APP12 peptide was added to serially diluted enzyme to a final concentration 200nM peptide in phosphate buffer (50mM pH 8, 0.005% Tween 20) and loaded onto a 384-flat bottom black plate. Polarization was measured using a Molecular Devices SpectraMax M5 plate reader (ex 550nm, em 580nm, 50reads per well). Binding affinities were determined by a best fit to a single site binding quadratic model.

### Rosetta Design Enzyme Screen

Enzyme screens were conducted by first incubating 2x peptide master mix (2 or 20μM peptide, 50mM phosphate buffer pH 8, 25mM ascorbic acid. Enzyme master mix was prepared by taking 10 or 100nM enzyme into 50mM phosphate buffer pH 8 and 2uM CuSO_4_. Master mixes were incubated at 25°C for 5min in a 96 well plate before starting reactions. Reactions were then initiated by mixing peptide and enzyme master mixes at 1:1 and time points were collected by quenching reactions to 10% acetic acid final. Samples were then diluted to ∼100pmols in 100uL. Samples were prepared for analysis by matrix-assisted laser desorption/ionization coupled to time of flight mass spectrometry (MALDI-TOF-MS) using Pierce C18 spin columns. Pierce C18 spin columns were prepared by adding 15uL of elution solution (80% acetonitrile and 0.5% acetic acid) and spinning at 500gs for 30sec. The tips then washed using 15uL of wash solution (4% acetonitrile and 0.5% acetic acid) and spinning at 500gs for 30sec. 70uL of sample (∼70pmol) was then loaded onto spin tip and spun at 400g for 5min. Tips were then washed twice with 15uL of wash solution and 300g: 30 seconds for the first spin and 6min for the second spin. Samples were eluted onto a 96well plate using 3uL of elution solution and spinning for 5min at 500g. Elution solution was allowed to evaporate under ambient conditions. Peptide was then resuspended using a half-saturated MALDI matrix solution (α-cyano-4-hydroxycinnamic acid in 80% acetonitrile and 0.5% acetic acid) and transferred to a 384 well MALDI plate.

### Matrix Assisted Laser Desorption/Ionization (MALDI) Data Acquisition and Analysis

MALDI-TOF-MS was performed on an AB SCIEX TOF/TOF 5800 in positive reflector mode. Wells were analyzed using a random walk and the laser intensity of 3200 to 4000 were used to avoid signal saturation. Each measurement was taken as the average of 2100 shots/well. Fraction modified was calculated as the sum of the integrals for the four most abundant product ion isotopes over the sum of the integrals for the four most abundant substrate and product ion isotopes.

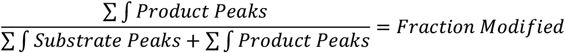

### Enzyme Activity Assays

Enzyme activity assays were conducted by first incubating 2x peptide master mix (20μM peptide, 100mM phosphate buffer pH 8, 2μM CuSO_4_ and 25mM ascorbic acid) and 2x enzyme stock (100nM in diH_2_O) at 25°C for 5min in a 96 well plate. Reactions were then initiated by mixing peptide and enzyme master mixes at 1:1 and time points were collected by taking reaction aliquots into a 10x tropolone quench solution (4mM tropolone final). Reaction samples were prepared for Liquid Chromatography– Mass Spectrometry (LCMS) by adding acetonitrile at 1:1 and spinning down at 16000rpm for 10min to remove any aggregate. The supernatant was then transferred to 100uL vials and stored at −20°C until they were analyzed by LCMS.

### Enzyme Mediated Conjugation of DM1 to Peptide

Cytotoxin-peptide bioconjugation reactions were conducted by first incubating 2x peptide master mix (20μM peptide, 50mM phosphate buffer pH 6.5, and 100μM of either MMAE or DM1) and 2x enzyme stock (500nM enzyme, 50mM phosphate buffer pH 6.5, and 2μM CuSO_4_) at 25°C for 5min in a 96 well plate. Reactions were then initiated by mixing peptide and enzyme master mixes at 1:1 and time points were collected by taking reaction aliquots into a 10x tropolone quench solution (4mM tropolone final). Reaction samples were prepared for Liquid Chromatography–Mass Spectrometry (LCMS) by adding acetonitrile at 1:1 and spinning down at 16000rpm for 10min to remove any aggregate. The supernatant was then transferred to 100uL vials and stored at −20°C until they were analyzed by LCMS.

### Liquid Chromatography–Mass Spectrometry (LCMS) Data Acquisition and Analysis

LCMS was performed on a Kinetex 2.6 µm C 8 column and Agilent 6520 Accurate-Mass Q-TOF ESI in high resolution mode. Reverse phase LC was performed at 0.5mL/min using a two-buffer gradient system: Buffer A (0.1% formic acid in water) and Buffer B (0.1% formic acid in ACN) with the following protocol:

| Solvent B (%) | Time (min) |
| --- | --- |
| 2 | 0-2 |
| 2-45 | 2-4 |
| 45-46.5 | 4.5-6.5 |
| 46.5-95 | 6.5-7.5 |
| 95 | 7.5-8.5 |
| 95-2 | 8.5-9.5 |

Fraction modified was calculated as the ratio of the product to the sum of the substrate and product EICs (10 or 100ppm threshold) (Figure S2-S8):

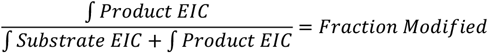

For the peptide-cytotoxic drug conjugation reactions, fraction modified was determined by extracting the EICs (10 ppm threshold) for the substrate (S) ion, dopa-product (DP) ion, orthoquinone-product (OP) ion, and cytotoxin conjugate-product (CCP) ion then calculating the ratio of the conjugate-product ion integral to the sum of the substrate and product ion integrals (Figure S10):

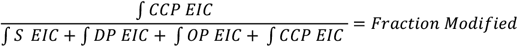

### Enzyme Mediated Conjugation of DM1 to Rituximab and Proteolytic Cytotoxic Payload Release

Rituximab antibody conjugation reactions were conducted by incubating 2x antibody master mix (10μM Rituximab, 100uM DM1, 50mM phosphate buffer pH 6.5) and 2x enzyme master mix (500nM enzyme, 50mM phosphate acetate pH 6.5 and 2μM CuSO_4_) in a 96well plate at 25°C for 5min. Reactions were initiated by adding antibody master mix to enzyme master mix at 1:1 and incubated at 25°C before quenching with 4mM tropolone (final). For the release of cytotoxic payload, 2x HRV 3C protease master mix (12.5uM HRV 3C and 50mM phosphate buffer pH 8) was then exposed to the quenched ADC reactions for at least 15min followed by antibody reduction with 10mM DTT for 30min. Reactions were then buffer exchanged into 50mM ammonium acetate solution at pH 6.5 or pH 8 using 10kDa MWCO Amicon spin tubes and stored at −80°C until ready to submit for intact mass spec.

### Extended Conjugation Reactions with DM1 and Rituximab

Rituximab antibody conjugation reactions were conducted by incubating 2x antibody master mix (0μM Rituximab, 50mM phosphate buffer pH 6.5) and 2x enzyme master mix (500nM enzyme, 100uM DM1, 50mM phosphate acetate pH 6.5 and 2μM CuSO_4_) in a 96well plate at 25°C for 5min. Reactions were initiated by adding antibody master mix to enzyme master mix at 1:1 and ran for 15 or 30 minutes at 25°C before quenching with 4mM tropolone (final). Reactions were subsequently reduced with 10mM DTT for 30min and buffer exchanged into 50mM ammonium acetate solution at pH 6.5 or pH 8 using 10kDa MWCO Amicon spin tubes. Reaction samples were stored at - 80°C until submitted for intact mass spectrometry.

### Intact Mass *Spectrometry*

Intact mass spec was performed at the Metabolomics and Proteomics (MAP) Core at the University of North Carolina. Briefly, samples were injected onto HR Chip (908Decives Inc.) using a ZipChip system interfaced to a Thermo QExactive HF Biopharma mass spectrometer. MS as acquired using Tune (v. 2.9) with a scan range of 1000-4000m/z, in-source CID 50 eV, mass resolution 15,000, 5 microscans AGC target 1e6, and 20ms maximum injection time. Intact mass spectra were deconvoluted using either the PMI Intact Mass software (Protein Metrics Inc) or Universal Deconvolution of Mass Spectra (UniDec).

