## supplemental_materials for "Enhancing enzymatic bioconjugation efficiency via installation of a substrate recruitment domain"

**Table S1.** Full list of peptide constructs used in this work. All peptides were ordered with C-terminal amides from the UNC High Throughput Peptide Synthesis and Array Facility (HTPSA).

| Name | Peptide # | Sequence[1] |
|---|---|---|
| *Directed Substrate* | 1 | **APPLPPR**NRPRLGSGSGSGSKNFLD**Y** |
| *Undirected Substrate* | 2 | AKAARSAEAKEAGRGSGSGSKNFLD**Y** |
| *Misdirected Substrate* | 3 | **Y**DLFNKGSGSGSGSGSGSGSGS**APPLPPR**NRPRL |
| *Reversed Substrate* | 4 | **Y**DLFNKGSGSGSGSGSVSLAR**RPLPPLP** |
| *Short Substrate* | 5 | **APPLPPR**NRPRLKNFLD**Y** |
| *Easy Substrate* | 6 | **APPLPPR**NRPRLGSGSGSGSGSGSG**Y** |
| *Difficult Substrate* | 7 | EW**APPLPPRNR**PRRSGGSGGSGGSGGSKET**Y**SK |
| *Negative Substrate* | 8 | **APPLPPR**NRPRLGSGSGSGSKEEEE**Y** |
| *TAMRA Substrate* | 9 | [TAMRA]-**APPLPPR**NRPRL |

[1]Recognition sequence and tyrosine residue are shown in bold and underlined

**Table S2.** Table of full-length protein sequences used in all experiments.

| **Protein Names** | **Design Sequences (6xHis+SUMO tag[1], Enzymatic Domain [Tyrosinase], Linker, Substrate Scaffold Domain [SH3])**<br>**Protease Sequences (GST, Linker, HRV 3C)**<br>**Antibody Sequences (Parent Antibody, Modification Tag)** |
|---|---|
| *bmTyr** | MGHHHHHHHHHHGSLQDSEVNQEAKPEVKPEVKPETHINLKVSDGSSEIFFKIKKTPLRRLMEAFAKRQGKEMDSLRFLYDGIRIQADQAPEDLDMEDNDIIEAHREQIGGGSYRVRKNVLHLTDTEKRDFVRTVLILKEKGIYDRYIAWHGAAGKFHTPPGSDRNAAHMSSAFLPWHREYLLRFERDLQSINPEVTLPYWEWETDAQMQDPSQSQIWSADFMGGNGNPIKDFIVDTGPFAAGRWTTIDEQGNPSGGLKRNFGATKEAPTLPTRDDVLNALKITQYDTPPWDMTSQNSFRNQLEGFINGPQLHNRVHHWVGGQMGVVPTAPNDPVFFLHHANVDRIWAVWQIIHRNQNYQPMKNGPFGQNFRDPMYPWNTTPEDVMNHRKLGYVYDIEL |
| *Naïve Fusion* | MGHHHHHHHHHHGSLQDSEVNQEAKPEVKPEVKPETHINLKVSDGSSEIFFKIKKTPLRRLMEAFAKRQGKEMDSLRFLYDGIRIQADQAPEDLDMEDNDIIEAHREQIGGKYRVRKNVLHLTDTEKRDFVRTVLILKEKGIYDRYIAWHGAAGKFHTPPGSDRNAAHMSSAFLPWHREYLLRFERDLQSINPEVTLPYWEWETDAQMQDPSQSQIWSADFMGGNGNPIKDFIVDTGPFAAGRWTTIDEQGNPSGGLKRNFGATKEAPTLPTRDDVLNALKITQYDTPPWDMTSQNSFRNQLEGFINGPQLHNRVHHWVGGQMGVVPTAPNDPVFFLHHANVDRIWAVWQIIHRNQNYQPMKNGPFGQNFRDPMYPWNTTPEDVMNHRKLGYVYDIELGSGSGSVTLFVALYDYEARTEDDLSFHKGEKFQILNSSEGDWWEARSLTTGETGYIPSNYVAPV |
| *Design 7* | MGHHHHHHHHHHGSLQDSEVNQEAKPEVKPEVKPETHINLKVSDGSSEIFFKIKKTPLRRLMEAFAKRQGKEMDSLRFLYDGIRIQADQAPEDLDMEDNDIIEAHREQIGGYRVRKNVLHLTDTEKRDFVRTVLILKEKGIYDRYIAWHGAAGKFHTPPGSDRNAAHMSSAFLPWHREYLLRFERDLQSINPEVTLPYWAWEVDAQMQDPSQSQIWSADFMGGNGNPIKDFIVDTGPFAAGRWTTIDEQGNPSGGLKRNFGATKEAPTLPTLDDVMEALKITQYDTPPWDMTSQNSFRNQLEGFINGPQLHNRVHHWVGGQMGVVPTAPNDPVFFLHHANVDRIWAVWQIIHRNQNYQPMKNGPFGQNFRDPMYPWNTTPEDVMNHRKLGYVYDKEGLIFVALYDYEARTEKDLSFHKGEKFQILNSDTGDWWLAVSLTTGELGYIPSNYVAPV |
| *Design 8* | MGHHHHHHHHHHGSLQDSEVNQEAKPEVKPEVKPETHINLKVSDGSSEIFFKIKKTPLRRLMEAFAKRQGKEMDSLRFLYDGIRIQADQAPEDLDMEDNDIIEAHREQIGGYRVRKNVLHLTDTEKRDFVRTVLILKEKGIYDRYIAWHGAAGKFHTPPGSDRNAAHMSSAFLPWHREYLLRFERDLQSINPEVTLPYWAWEYDAQMQDPSQSQIWSADFMGGNGNPIKDFIVDTGPFAAGRWTTIDEQGNPSGGLKRNFGATKEAPTLPTAADVARALKITQYDTPPWDMTSQNSFRNQLEGFINGPQLHNRVHHWVGGQMGVVPTAPNDPVFFLHHANVDRIWAVWQIIHRNQNYQPMKNGPFGQNFRDPMYPWNTTPEDVMNHRKLGYVYDTFQPEGAFVALYDYEARTEDDLSFHKGEKFAIILTAAGDWWLALSLTTGEIGYIPSNYVAPV |
| *Design 12* | MGHHHHHHHHHHGSLQDSEVNQEAKPEVKPEVKPETHINLKVSDGSSEIFFKIKKTPLRRLMEAFAKRQGKEMDSLRFLYDGIRIQADQAPEDLDMEDNDIIEAHREQIGGYRVRKNVLHLTDTEKRDFVRTVLILKEKGIYDRYIAWHGAAGKFHTPPGSDRNAAHMSSAFLPWHREYLLRFERDLQSINPEVTLPYWAWETDAQMQDPSQSQIWSADFMGGNGNPIKDFIVDTGPFAAGRWTTIDEQGNPSGGLKRNFGATKEAPTLPTLDDVLNALKITQYDTPPWDMTSQNSFRNQLEGFINGPQLHNRVHHWVGGQMGVVPTAPNDPVFFLHHANVDRIWAVWQIIHRNQNYQPMKNGPFGQNFRDPMYPWNTTPEDVMNHRKLGYVYDQEMALFVALYDYEARTEKDLSFHKGEKFAILNTTLGDWWIAYSLTTGEVGYIPSNYVAPV |
| *Design 40* | MGHHHHHHHHHHGSLQDSEVNQEAKPEVKPEVKPETHINLKVSDGSSEIFFKIKKTPLRRLMEAFAKRQGKEMDSLRFLYDGIRIQADQAPEDLDMEDNDIIEAHREQIGGYRVRKNVLHLTDKEKRDFVRTVLILKEKGIYDRYIAWHGAAGKFHTPPGSDRNAAHMSSAFLPWHREYLLRFERDLQSINPEVTLPYWPWETDAQMQDPSQSQIWSADFMGGNGNPIKDFIVDTGPFAAGRWTTIDEQGNPSGGLKRNFGATKEAPTLPTRADVWKALAITQYDTPPWDMTSQNSFRNQLEGFINGPQLHNRVH**H**WVGGQMGVVPTAPNDPVFFLHHANVDRIWAVWQIIHRNQNYQPMKNGPFGQNFRDPMYPWNTTPEDVMNHRKLGYVYDIEDKREIGPIFVALYDYEARTEDDLSFHKGEKFQILDSSEGDWWMARSLTTGEIGYIPSNYVAPV |
| *Design 42* | MGHHHHHHHHHHGSLQDSEVNQEAKPEVKPEVKPETHINLKVSDGSSEIFFKIKKTPLRRLMEAFAKRQGKEMDSLRFLYDGIRIQADQAPEDLDMEDNDIIEAHREQIGGYRVRKNVLHLTDTEKRDFVRTVLILKEKGIYDRYIAWHGAAGKFHTPPGSDRNAAHMSSAFLPWHREYLLRFERDLQSINPEVTLPYWDWTTDAQMQDPSQSQIWSADFMGGNGNPIKDFIVDTGPFAAGRWTTIDEQGNPSGGLKRNFGATKEAPTLPTRADVVKALAITQYDTPPWDMTSQNSFRNQLEGFINGPQLHNRVH**H**WVGGQMGVVPTAPNDPVFFLHHANVDRIWAVWQIIHRNQNYQPMKNGPFGQNFRDPMYPWNTTPEDVMNHRKLGYVYDIEDRNDINGPIFVALYDYEARTEDDLSFHKGEKFQILDSSEGDWWAARSLTTGEIGYIPSNYVAPV |
| *Design 48* | MGHHHHHHHHHHGSLQDSEVNQEAKPEVKPEVKPETHINLKVSDGSSEIFFKIKKTPLRRLMEAFAKRQGKEMDSLRFLYDGIRIQADQAPEDLDMEDNDIIEAHREQIGGYRVRKNVLHLTDTEKRDFVRTVLILKEKGIYDRYIAWHGAAGKFHTPPGSDRNAAHMSSAFLPWHREYLLRFERDLQSINPEVTLPYWAWEYDAQMQDPSQSQIWSADFMGGNGNPIKDFIVDTGPFAAGRWTTIDEQGNPSGGLKRNFGATKEAPTLPTAADVARALKITQYDTPPWDMTSQNSFRNQLEGFINGPQLHNRVHHWVGGQMGVVPTAPNDPVFFLHHANVDRIWAVWQIIHRNQNYQPMKNGPFGQNFRDPMYPWNTTPEDVMNHRKLGYVYDMEGEEKESAFVALYDYEARTEDDLSFHKGEKFAIILTAEGDWWLALSLTTGEIGYIPSNYVAPV |

| | |
|---|---|
| *GST-HRV 3C (Prescission Protease)* | MSPILGYWKIKGLVQPTRLLLEYLEEKYEEHLYERDEGDKWRNKKFELGLEFPNLPYYIDGDVKLTQSMAIIRYI<br>ADKHNMLGGCPKERAEISMLEGAVLDIRYGVSRIAYSKDFETLKVDFLSKLPEMLKMFEDRLCHKTYLNGDHV<br>THPDFMLYDALDVVLYMDPMCLDAFPKLVCFKKRIEAIPQIDKYLKSSKYIAWPLQGWQATFGGGDHPPKSDL<br>VPRGSGPNTEFALSLLRKNIMTITTSKGEFTGLGIHDRVCVIPTHAQPGDDVLVNGQKIRVKDKYKLVDPENINL<br>ELTVLTLDRNEKFRDIRGFISEDLEGVDATLVVHSNNFTNTILEVGPVTMAGLINLSSTPTNRMIRYDYATKTGQ<br>CGGVLCATGKIFGIHVGGNGRQGFSAQLKKQYFVEKQ |
| *Rituximab HC Tag 2* | QVQLQQPGAELVKPGASVKMSCKASGYTFTSYNMHWVKQTPGRGLEWIGAIYPGNGDTSYNQKFKGKATL<br>TADKSSSTAYMQLSSLTSEDSAVYYCARSTYYGGDWYFNVWGAGTTVTVSAASTKGPSVFPLAPSSKSTSG<br>GTAALGCLVKDYFPEPVTVSWNSGALTSGVHTFPAVLQSSGLYSLSSVVTVPSSSLGTQTYICNVNHKPSNT<br>KVDKKVEPKSCTHTCPPCPAPELLGGPSVFLFPPKPKDTLMISRTPEVTCVVVDVSHEDPEVKFNWYVDGVE<br>VHNAKTKPREEQYQSTYRVVSVLTVLHQDWLNGKEYKCKVSNKALPAPIEKTISKAKGQPREPQVYTLPPSR<br>EEMTKNQVSLTCLVKGFYPSDIAVEWESNGQPENNYKTTPPVLDSDGSFFLYSKLTVDKSRWQQGNVFSCS<br>VMHEALHNHYTQKSLSLSPGK |
| *Rituximab LC Tag 2* | QIVLSQSPAILSASPGEKVTMTCRASSSVSYIHWFQQKPGSSPKPWIYATSNLASGVPVRFSGSGSGTSYSLT<br>ISRVEAEDAATYYCQQWTSNPPTFGGGTKLEIKRTVAAPSVFIFPPSDEQLKSGTASVVCLLNNFYPREAKVQ<br>WKVDNALQSGNSQESVTEQDSKDSTYSLSSTLTLSKADYEKHKVYACEVTHQGLSSPVTKSFNRGECGSGS<br>GSAPPLPPRNRPRLLEVLFQGPSGSGKGSGSYAAA |

[1]6xHis+SUMO selection and expression tags are included as part of the pCDB24_XhoI plasmids ordered from Twist Bioscience

**Table S3.** Table of genes ordered from Twist Bioscience using pCDB24_XhoI plasmid backbone.

| Design Names | Gene Inserts Ordered from Twist Bioscience |
|---|---|
| *bmTyr** | GGGTCATATCGGGTCAGAAAAAACGTGCTGCATCTTACGGACACTGAAAAACGCGACTTCGTCCGCAC<br>GGTGTTGATTTTGAAAGAAAAAGGGATTTATGATCGCTACATTGCGTGGCATGGTGCAGCGGGGAAAT<br>TTCACACCCCACCTGGCTCCGACCGGAATGCGGCGCACATGTCATCGGCATTCCTGCCGTGGCATCG<br>CGAATACCTGTTACGTTTCGAGCGGGACTTGCAATCCATCAACCCTGAGGTAACCCTTCCATATTGGG<br>AATGGGAGACGGATGCACAGATGCAAGACCCCTCACAATCGCAGATATGGAGCGCAGACTTTATGGG<br>GGGTAACGGCAACCCGATCAAAGATTTCATTGTGGATACGGGGCCTTTCGCCGCCGGTCGTTGGACG<br>ACCATAGATGAGCAAGGAAATCCTTCAGGGGGATTAAAGAGAAACTTTGGGGCAACTAAAGAAGCACC<br>AACTTTACCGACCCGCGACGATGTATTAAACGCACTTAAGATCACTCAGTATGACACCCCCCCATGGG<br>ACATGACATCGCAGAATAGCTTCCGTAACCAACTGGAGGGTTTCATCAATGGGCCGCAACTGCACAAT<br>CGCGTTCACCATTGGGTTGGCGGGCAGATGGGGGTCGTACCCACAGCGCCCAACGACCCTGTGTTCT<br>TTCTTCATCACGCAAACGTGGACCGGATCTGGGCTGTTTGGCAGATTATCCATAGAAACCAGAATTATC<br>AACCGATGAAGAATGGCCCGTTTGGCCAAAATTTCCGCGACCCCATGTATCCATGGAATACCACGCCC<br>GAGGACGTGATGAATCACCGGAAGTTAGGTTATGTATACGATATTGAATTATAATAG |
| *Naïve Fusion* | AAATACCGCGTACGGAAGAACGTTCTGCATTTGACAGACACGGAGAAGCGTGACTTTGTCCGCACGGT<br>GCTGATTTTGAAGGAGAAAGGCATCTATGATAGATATATCGCATGGCATGGAGCGGCTGGGAAATTTC<br>ATACTCCTCCGGGCTCCGACCGTAACGCGGCCCACATGAGTTCGGCGTTTCTTCCCTGGCACCGTGA<br>GTACTTACTTCGTTTCGAGAGAGATTTGCAAAGTATAAATCCTGAGGTTACTCTGCCCTATTGGGAGTG<br>GGAGACCGATGCTCAAATGCAGGACCCATCTCAATCGCAAATCTGGAGCGCCGACTTTATGGGGGGG<br>AACGGAAATCCGATCAAAGACTTCATCGTAGACACTGGGCCCTTTGCTGCCGGGAGATGGACGACTAT<br>CGATGAACAAGGGAACCCATCCGGGGGTTTGAAACGCAACTTTGGCGCTACGAAAGAGGCGCCCACC<br>CTGCCCACTCGGGACGATGTGTTGAACGCTCTGAAAATAACCCAGTATGACACACCCCCGTGGACAT<br>GACTTCGCAGAACTCTTTTCGTAATCAGCTTGAAGGGTTCATCAATGGGCCTCAGTTACATAATCGTGT<br>GCATCATTGGGTGGGCGGACAAATGGGAGTCGTACCCACGGCCCCTAATGACCCTGTCTTTTTTCTTC<br>ATCATGCTAATGTGGACCGGATTTGGGCGGTGTGGCAAATCATCCACCGGAACCAGAACTACCAGCCC<br>ATGAAAAATGGGCCGTTCGGGCAGAATTTTAGAGATCCTATGTATCCGTGGAATACAACACCTGAGGA<br>TGTCATGAACCATAGAAAATTGGGCTACGTCTATGATATAGAACTTGGCAGTGGCAGCGGCTCAGTTA<br>CCTTGTTCGTCGCCCTGTACGATTATGAAGCTCGTACGGAAGATGATCTGTCATTTCACAAGGGGGAA<br>AAATTTCAGATCCTTAACTCGTCTGAAGGGGATTGGTGGGAAGCCCGGAGTCTTACTACAGGTGAGAC<br>TGGTTACATTCCTTCTAACTATGTAGCCCCGGTG |

| | |
|---|---|
| *Design 7* | TATCGGGTACGTAAAAATGTTTTGCATTTGACCGATACAGAAAAAAGAGATTTTGTGCGTACGGTTTTAA<br>TACTTAAAGAAAAGGGGATCTATGACCGCTATATTGCTTGGCACGGAGCTGCTGGCAAGTTTCACACC<br>CCACCAGGTAGTGACCGGAATGCAGCACACATGTCGAGTGCTTTTTTACCCTGGCACCGTGAATACCT<br>GCTGAGATTCGAGCGGGATCTTCAATCTATTAATCCTGAAGTGACATTACCTTACTGGGCCTGGGAGG<br>TAGATGCGCAAATGCAGGACCCTTCCCAGTCCCAAATTTGGTCAGCAGACTTCATGGGTGGGAACGGC<br>AACCCCATTAAAGATTTTATTGTAGACACTGGCCCGTTTGCTGCTGGTCGTTGGACGACAATTGACGAG<br>CAGGGAAATCCTAGCGGAGGGTTAAAGCGTAATTTCGGGGCCACGAAAGAGGCGCCGACCCTGCCCA<br>CTTTAGATGACGTGATGGAAGCATTAAAAATAACCCAATACGATACCCCGCCTTGGGATATGACGTCTC<br>AAAACTCCTTTCGGAACCAACTGGAGGGGTTCATTAACGGCCCGCAGCTTCATAACCGGGTACATCAC<br>TGGGTTGGGGGGCAGATGGGCGTAGTTCCCACCGCACCGAACGATCCAGTGTTTTTCTTACATCATGC<br>TAACGTCGACCGTATTTGGGCTGTTTGGCAAATTATTCATAGAAACCAAAATTATCAACCTATGAAGAAT<br>GGTCCATTCGGACAGAATTTTCGGGACCCTATGTATCCTTGGAACACCACGCCTGAGGACGTTATGAA<br>TCACAGAAAATTGGGCTATGTGTACGACAAGGAAGGGTTAATCTTTGTGGCTCTGTATGATTACGAGGC<br>GCGGACCGAGAAAGACTTGAGTTTCCACAAAGGCGAGAAGTTTCAGATCCTTAATAGTGACACAGGGG<br>ACTGGTGGCTTGCTGTCTCCTTGACTACTGGCGAACTTGGATACATACCAAGCAACTATGTGGCGCCG<br>GTG |
| *Design 8* | TATCGTGTACGGAAAAACGTATTGCATTTAACAGACACAGAGAAGAGAGACTTTGTAAGAACTGTCTT<br>AATCCTGAAAGAGAAGGGCATCTATGACAGATATATAGCTTGGCATGGGGCCGCAGGCAAGTTTCAC<br>ACTCCACCCGGATCTGACCGGAATGCGGCCCACATGAGTAGTGCGTTCTTGCCATGGCATCGGGAG<br>TATTTGCTTCGGTTTGAACGCGACTTACAGTCGATAAATCCAGAAGTGACCTTGCCTTATTGGGCGTG<br>GGAGTACGACGCCCAAATGCAAGACCCGAGCCAGTCACAGATATGGTCAGCCGACTTCATGGGAGG<br>TAACGGTAACCCAATCAAAGATTTCATTGTGGACACTGGACCGTTTGCTGCGGGACGCTGGACTACG<br>ATCGATGAGCAAGGAAATCCGTCAGGCGGACTTAAGAGAAACTTTGGCGCGACAAAGGAAGCTCCA<br>ACTCTTCCGACCGCAGCCGATGTGGCCCGTGCCTTGAAGATCACTCAATACGACACTCCCCCGTGG<br>GATATGACAAGTCAAAATTCGTTTCGGAATCAACTTGAGGGCTTTATCAATGGGCCGCAATTACATAA<br>TAGAGTTCACCACTGGGTAGGTGGACAGATGGGAGTGGTGCCCACTGCTCCGAATGATCCAGTTTT<br>CTTCTTGCATCACGCAAATGTAGACCGTATCTGGGCCGTGTGGCAAATCATTCATCGCAACCAAAATT<br>ACCAACCCATGAAGAACGGGCCTTTCGGACAGAATTTTCGTGATCCGATGTACCCCTGGAATACCAC<br>TCCGGAAGACGTTATGAATCATCGTAAACTGGGGTACGTGTACGACACTTTCCAACCAGAGGGTGCT<br>TTTGTGGCGCTGTATGACTACGAGGCCCGGACGGAAGACGATCTTTCGTTTCATAAGGGAGAGAAAT<br>TTGCAATTATCCTGACCGCAGCGGGGGATTGGTGGCTGGCCTTATCACTGACTACAGGGGAGATCG<br>GGTACATACCTTCGAACTATGTGGCCCCTGTG |
| *Design 12* | TATAGAGTTCGTAAAAATGTGTTGCACTTAACAGATACGGAAAAGCGGGACTTTGTTCGTACAGTATT<br>AATACTGAAAGAGAAGGGGATCTATGATAGATATATAGCATGGCATGGAGCCGCCGGCAAATTTCAC<br>ACTCCCCCCGGCTCAGACCGGAACGCCGCGCACATGTCATCCGCATTTTTACCGTGGCATAGAGAA<br>TACCTTTTACGTTTTGAACGGGACTTGCAAAGCATAAACCCGGAAGTGACACTTCCTTACTGGGCGT<br>GGGAGACCGACGCCCAGATGCAGGACCCGTCGCAGTCTCAGATTTGGTCCGCCGATTTCATGGGTG<br>GCAACGGTAATCCTATAAAGGACTTCATTGTAGACACGGGGCCGTTCGCTGCGGGCCGCTGGACGA<br>CGATCGACGAGCAGGGTAACCCGTCCGGCGGGCTTAAACGCAATTTTGGAGCCACTAAGGAAGCTC<br>CTACATTACCGACACTGGATGATGTGTTAAATGCCCTTAAGATTACTCAGTACGACACACCTCCTTGG<br>GATATGACTAGCCAAAACTCTTTCCGTAACCAGTTGGAAGGCTTTATAAACGGGCCTCAGTTACACAA<br>CAGAGTCCATCATTGGGTTGGTGGGCAGATGGGCGTTGTCCCAACCGCGCCTAATGACCCCGTATT<br>TTTCCTTCATCATGCTAATGTCGACCGCATCTGGGCTGTCTGGCAGATAATTCACCGTAACCAAAACT<br>ATCAGCCAATGAAGAATGGTCCCTTCGGACAAAACTTCCGGGACCCCATGTACCCTTGGAACACTAC<br>ACCAGAAGACGTGATGAACCATAGAAAACTTGGTTACGTCTATGACCAGGAAATGGCCCTGTTTGTA<br>GCACTTTACGATTACGAGGCACGCACCGAAAAAGACTTGTCGTTTCACAAGGGTGAAAAGTTCGCTA<br>TATTGAATACAACCTTAGGGGACTGGTGGATCGCTTATTCGCTTACAACAGGAGAGGTCGGCTACAT<br>CCCCTCAAACTATGTCGCCCCGGTG |

| | |
|---|---|
| *Design 40* | TATAGAGTCCGGAAGAACGTCTTGCACCTGACCGATAAAGAAAAGAGAGATTTTGTTAGAACGGTGT<br>TAATTTTGAAAGAAAAAGGAATATATGATCGCTATATCGCGTGGCACGGGGCCGCTGGCAAATTCCA<br>CACACCTCCGGGATCTGATCGTAACGCTGCGCACATGTCTTCCGCTTTCCTTCCTTGGCACAGAGAG<br>TATTTGTTAAGATTCGAGCGGGATTTACAATCCATCAATCCCGAGGTGACACTTCCGTATTGGCCATG<br>GGAAACGGACGCTCAAATGCAGGACCCCTCCCAATCTCAGATCTGGTCAGCGGACTTCATGGGAGG<br>GAACGGCAACCCCATTAAGGATTTTATCGTTGACACGGGTCCGTTTGCGGCAGGACGCTGGACGAC<br>GATCGACGAGCAAGGTAATCCATCAGGAGGCCTGAAACGTAATTTCGGAGCAACCAAAGAAGCCCC<br>AACTTTACCTACACGCGCAGACGTGTGGAAAGCACTGGCGATCACACAGTACGACACGCCGCCATG<br>GGACATGACGAGTCAAAATTCATTTAGAAATCAACTGGAAGGTTTCATAAACGGACCACAGCTTCACA<br>ACCGTGTCCACCATTGGGTGGGCGGACAGATGGGCGTCGTTCCTACTGCCCCTAACGATCCAGTAT<br>TCTTCTTACATCATGCAAATGTGGATCGCATCTGGGCAGTATGGCAAATAATTCACCGCAACCAGAAT<br>TACCAACCTATGAAAAATGGTCCTTTTGGCCAGAATTTTCGCGATCCCATGTATCCGTGGAATACCAC<br>TCCGGAAGACGTGATGAATCACCGTAAATTAGGCTACGTGTACGACATAGAAGATAAACGTGAGATT<br>GGCCCAATATTTGTGGCGCTGTATGATTACGAGGCACGCACGGAAGACGACTTGTCCTTTCACAAAG<br>GGGAGAAATTCCAGATCTTGGACTCATCGGAGGGTGACTGGTGGATGGCTAGAAGCCTTACGACAG<br>GTGAAATCGGGTACATACCAAGCAATTACGTTGCACCCGTG |
| *Design 42* | TATCGTGTTCGTAAGAATGTGTTACACTTGACTGACACGGAGAAACGGGATTTTGTCCGCACAGTGC<br>TTATCCTGAAGGAAAAGGGGATCTACGACCGCTATATCGCCTGGCATGGCGCAGCAGGTAAGTTCC<br>ACACCCCACCGGGATCGGACCGCAATGCAGCACATATGAGTAGCGCCTTCTTGCCATGGCATAGAG<br>AATACTTATTGAGATTTGAACGCGATCTGCAATCGATAAACCCGGAAGTAACGTTGCCGTACTGGGA<br>CTGGACGACCGATGCACAGATGCAAGATCCTTCGCAATCGCAAATATGGTCAGCCGACTTCATGGG<br>GGGAAACGGCAATCCCATAAAGGACTTTATTGTTGACACCGGTCCCTTTGCTGCAGGTCGGTGGACA<br>ACCATTGATGAACAGGGGAATCCAAGTGGCGGGTTAAAACGGAACTTCGGCGCCACCAAGGAAGCG<br>CCAACCTTACCAACCCGCGCGGACGTAGTGAAGGCATTAGCAATAACACAATACGATACACCACCAT<br>GGGACATGACCTCTCAAAACAGTTTCCGCAATCAGCTTGAAGGTTTTATAAATGGTCCCCAACTTCAC<br>AATCGTGTCCATCATTGGGTGGGAGGTCAGATGGGTGTCGTACCCACGGCGCCGAATGACCCTGTT<br>TTCTTCTTGCATCATGCTAACGTTGACAGAATCTGGGCTGTTTGGCAAATCATACATCGGAACCAGAA<br>CTACCAACCAATGAAAAATGGGCCATTCGGTCAAAATTTTCGTGACCCTATGTACCCCTGGAACACTA<br>CACCGGAGGATGTAATGAATCACAGAAAACTTGGGTACGTCTACGATATAGAGGACCGCAACGATAT<br>AAACGGCCCAATCTTCGTCGCGCTGTACGACTATGAAGCGAGAACGGAGGATGATCTGTCTTTCCAC<br>AAAGGCGAAAAGTTCCAGATTCTTGATTCGTCTGAAGGGGACTGGTGGGCGGCACGCAGCCTTACG<br>ACTGGCGAAATTGGGTACATACCGAGTAACTATGTTGCACCTGTG |
| *Design 48* | TATAGAGTGCGTAAGAATGTGCTTCATTTGACGGACACTGAAAAGAGAGATTTTGTCCGTACTGTTTT<br>AATACTTAAAGAGAAAGGGATCTATGATCGGTATATAGCCTGGCATGGTGCGGCGGGAAAGTTCCAC<br>ACACCGCCCGGTTCAGACCGTAACGCCGCCCATATGAGTTCAGCCTTTTTGCCATGGCATCGGGAA<br>TACTTACTGAGATTCGAAAGAGACCTGCAGTCCATTAATCCGGAGGTTACTTTGCCTTATTGGGCTTG<br>GGAGTATGATGCGCAGATGCAAGATCCGTCCCAGAGTCAGATTTGGTCCGCTGACTTCATGGGTGG<br>GAATGGTAATCCAATAAAAGACTTTATCGTAGACACTGGTCCTTTCGCGGCGGGGCGTTGGACAACA<br>ATTGACGAACAGGGTAATCCTTCTGGCGGACTTAAACGTAACTTTGGAGCTACGAAAGAAGCCCCGA<br>CTTTGCCGACCGCAGCCGATGTAGCAAGAGCCTTGAAGATCACACAATATGACACGCCGCCATGGG<br>ATATGACATCCCAGAATTCGTTTCGGAATCAGCTGGAAGGCTTCATAAACGGGCCGCAGTTACACAA<br>CCGTGTTCACCATTGGGTAGGAGGCCAGATGGGCGTAGTTCCCACAGCTCCCAATGACCCGGTATT<br>CTTCCTTCACCACGCAAACGTAGATCGTATATGGGCAGTTTGGCAAATTATCCATCGTAATCAAAATT<br>ACCAACCAATGAAGAACGGTCCTTTCGGACAAAATTTTCGTGATCCAATGTACCCTTGGAATACAACT<br>CCGGAGGACGTTATGAATCATAGAAAGTTGGGCTACGTTTATGACATGGAGGGGGAGGAAAAGGAA<br>AGTGCCTTCGTCGCATTGTATGACTACGAAGCACGTACTGAGGATGACCTGTCCTTTCACAAGGGAG<br>AGAAATTTGCGATAATACTTACCGCCGAAGGGGACTGGTGGCTTGCCTTAAGTCTTACAACGGGTGA<br>GATTGGATATATTCCGTCCAATTACGTTGCCCCCGTG |

| | |
|---|---|
| *GST-HRV 3C (Prescission Protease)* | ATGTCCCCTATACTAGGTTATTGGAAAATTAAGGGCCTTGTGCAACCCACTCGACTTCTTTTGGAATA<br>TCTTGAAGAAAAATATGAAGAGCATTTGTATGAGCGCGATGAAGGTGATAAATGGCGAAACAAAAAG<br>TTTGAATTGGGTTTGGAGTTTCCCAATCTTCCTTATTATATTGATGGTGATGTTAAATTAACACAGTCT<br>ATGGCCATCATACGTTATATAGCTGACAAGCACAACATGTTGGGTGGTTGTCCAAAAGAGCGTGCAG<br>AGATTTCAATGCTTGAAGGAGCGGTTTTGGATATTAGATACGGTGTTTCGAGAATTGCATATAGTAAA<br>GACTTTGAAACTCTCAAAGTTGATTTTCTTAGCAAGCTACCTGAAATGCTGAAAATGTTCGAAGATCG<br>TTTATGTCATAAAACATATTTAAATGGTGATCATGTAACCCATCCTGACTTCATGTTGTATGACGCTCT<br>TGATGTTGTTTTATACATGGACCCAATGTGCCTGGATGCGTTCCCAAAATTAGTTTGTTTTAAAAAACG<br>TATTGAAGCTATCCCACAAATTGATAAGTACTTGAAATCCAGCAAGTATATAGCATGGCCTTTGCAGG<br>GCTGGCAAGCCACGTTTGGTGGTGGCGACCATCCTCCAAAATCGGATCTGGTTCCGCGTGGATCCG<br>GTCCGAATACCGAATTTGCACTGAGCCTGCTGCGTAAAAACATTATGACCATTACCACCTCCAAAGG<br>CGAATTTACCGGTCTGGGTATTCATGATCGTGTTTGTGTTATTCCGACCCATGCACAGCCTGGTGAT<br>GATGTTCTGGTTAATGGTCAGAAAATTCGCGTGAAAGATAAATACAAACTGGTGGACCCGGAAAACA<br>TTAATCTGGAACTGACCGTTCTGACCCTGGATCGTAATGAAAAATTTCGTGATATCCGTGGCTTTATC<br>AGCGAAGATCTGGAAGGTGTTGATGCAACCCTGGTTGTTCATAGCAATAACTTTACCAACACCATTCT<br>GGAAGTTGGTCCGGTTACAATGGCAGGTCTGATTAATCTGAGCAGCACCCCGACCAATCGTATGATT<br>CGTTATGATTATGCAACCAAAACCGGTCAGTGTGGTGGTGTTCTGTGTGCAACCGGTAAAATCTTTG<br>GCATTCATGTTGGTGGCAATGGTCGTCAGGGTTTTAGCGCACAGCTGAAAAAACAGTATTTCGTCGA<br>AAAACAGTAA |
| *Rituximab HC* | ATGAAGTGGGTAACCTTTATTTCCTTGCTGTTCCTCTTTAGCTCGGCTTATTCCCAAGTGCAGCTGCA<br>ACAGCCCGGCGCCGAGCTGGTGAAGCCTGGAGCCAGCGTGAAAATGAGCTGCAAGGCTAGCGGTT<br>ATACCTTCACCTCCTACAACATGCACTGGGTGAAGCAGACCCCTGGAAGAGGCCTTGAGTGGATCG<br>GCGCTATCTACCCTGGAAATGGCGATACAAGCTACAATCAAAAGTTCAAGGGCAAAGCTACACTGAC<br>AGCCGACAAGAGCAGCTCCACAGCCTACATGCAGCTGAGCTCTCTGACCAGCGAGGACAGCGCCG<br>TGTACTACTGCGCCAGATCTACTTACTACGGCGGCGACTGGTATTTCAACGTGTGGGGAGCCGGCA<br>CCACCGTGACAGTGTCTGCTGCTTCTACCAAGGGCCCAAGCGTATTCCCACTGGCCCCTTCCAGTAA<br>GTCCACCAGCGGAGGCACCGCCGCCCTGGGCTGTCTGGTCAAGGACTACTTCCCTGAGCCTGTTAC<br>CGTGTCTTGGAACAGCGGCGCCCTGACCTCCGGCGTGCATACATTTCCTGCCGTGCTGCAGAGCTC<br>AGGCCTGTACAGCCTCTCTAGCGTGGTCACCGTGCCTAGCTCTAGCCTGGGCACACAGACATACAT<br>CTGCAACGTGAACCACAAACCTAGCAACACCAAGGTGGATAAGAAGGTGGAACCTAAGAGTTGTACC<br>CACACCTGCCCGCCGTGCCCTGCTCCTGAGCTGCTGGGCGGCCCCTCCGTGTTTCTGTTCCCTCCT<br>AAGCCCAAAGACACCCTGATGATCAGCAGAACCCCCGAGGTCACATGTGTGGTGGTGGACGTGAGC<br>CACGAAGATCCTGAGGTGAAATTCAACTGGTACGTGGACGGCGTCGAGGTGCACAATGCCAAGACC<br>AAACCCCGGGAAGAACAGTACCAGAGCACATATAGAGTGGTCAGCGTGCTGACCGTGCTGCACCAG<br>GACTGGCTGAACGGCAAGGAATACAAGTGCAAAGTGTCTAACAAGGCCCTGCCCGCCCCAATTGAG<br>AAGACCATCAGCAAGGCCAAGGGACAGCCTAGAGAGCCTCAGGTGTACACACTGCCTCCTAGCAGA<br>GAAGAGATGACCAAGAACCAGGTGTCCCTGACCTGCCTGGTGAAGGGGTTCTACCCCAGCGACATC<br>GCCGTGGAATGGGAGAGCAACGGCCAGCCCGAAAACAACTACAAGACCACACCCCCCGTGCTGGA<br>TTCTGATGGCAGCTTCTTCCTGTACAGCAAGCTGACCGTTGACAAGAGCCGGTGGCAGCAGGGCAA<br>CGTGTTCAGCTGCAGCGTGATGCACGAGGCCCTGCACAACCACTACACCCAGAAATCTCTGAGCCT<br>CAGCCCCGGAAAG |
| *Rituximab LC* | ATGAAGTGGGTAACCTTTATTTCCTTGCTGTTCCTCTTTAGCTCGGCTTATTCCCAAATCGTACTCTCT<br>CAATCCCCGGCTATTCTTAGCGCCAGTCCTGGCGAAAAGGTGACCATGACATGCAGGGCCTCATCTT<br>CTGTTAGCTATATCCATTGGTTCCAGCAAAAACCAGGAAGCTCCCCCAAGCCGTGGATTTATGCGAC<br>GAGTAACTTGGCTTCTGGAGTACCTGTGCGCTTCTCAGGCTCCGGTAGTGGCACCTCATACTCCCTG<br>ACAATAAGTAGAGTCGAGGCAGAGGACGCGGCAACGTACTACTGTCAACAATGGACATCTAACCCG<br>CCTACATTCGGTGGAGGCACGAAGCTGGAAATTAAAAGAACGGTTGCTGCGCCGAGTGTATTCATCT<br>TCCCTCCCTCAGACGAACAATTGAAAAGTGGTACAGCTTCCGTAGTCTGTTTGCTCAATAATTTTTAC<br>CCTAGAGAAGCGAAGGTGCAGTGGAAGGTCGACAACGCGCTTCAAAGTGGAAATTCCCAAGAGAGT<br>GTTACAGAGCAAGACAGCAAGGATTCAACATACAGTTTGTCTTCTACTTTGACTCTCTCTAAGGCCGA<br>CTATGAGAAACACAAAGTATACGCTTGTGAGGTTACTCACCAAGGATTGTCCAGCCCGGTAACTAAG<br>AGCTTCAATAGGGGTGAGTGTGGCAGCGGCTCTGGCTCCGCCCCTCCACTGCCTCCACGCAACAGA<br>CCTCGGCTACTGGAAGTGCTGTTTCAGGGCCCTTCCGGAAGCGGCAAGGGATCCGGCAGCTACGC<br>TGCCGCTTGATGA |

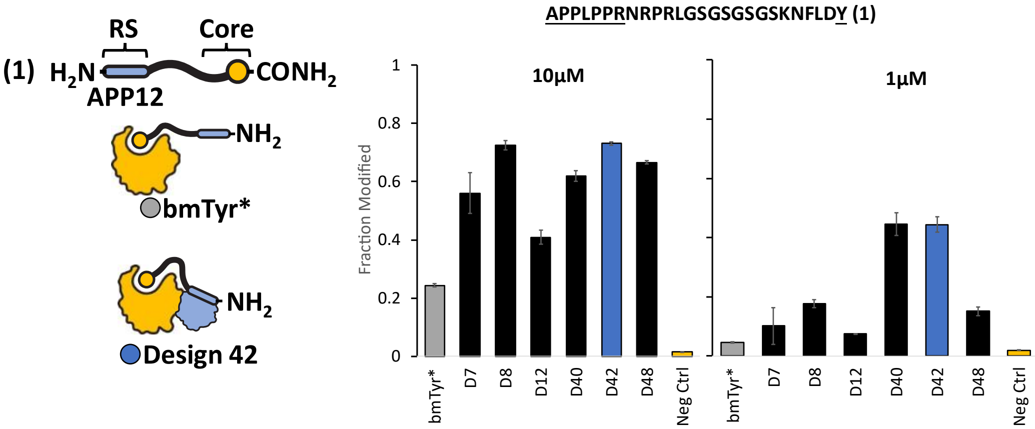

**Figure S1.** Enzyme design screen against directed peptide substrate. Incubation (**A**) 50nM enzyme with 10µM peptide 1 (directed substrate) or (**B**) 5nM enzyme with 1µM directed peptide substrate for 2min. Fraction modified is calculated as area under curve for hydroxylated peptide over sum of substrate and hydroxylated peptide as measured using MALDI. Bars show mean of triplicate wells and error bars indicate S.D. The top performing enzyme (D42) is shown with blue bar while bmTyr* activity and NF is shown with grey and orange bars respectively. Yellow bar show amount of hydroxylated peptide present at end of reaction in absence of enzyme.

A

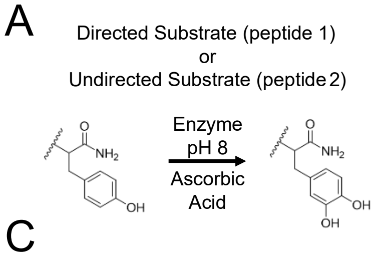

B

| Directed Substrate (Peptide 1) | | | Undirected Substrate (Peptide 2) | | |
|---|---|---|---|---|---|
| APPLPPRNRPRLGSGSGSGSKNFLD**Y** | | | AKAARSAEAKEAGRGSGSGSKNFLD**Y** | | |
| **Charge** | **Substrate** | **Product** | **Charge** | **Substrate** | **Product** |
| 0 | 2739.4398 | 2755.4348 | 0 | 2627.3245 | 2641.3402 |
| $[M+4]^{+4}$ | 685.8672 | 689.8660 | $[M+4]^{+4}$ | 657.8384 | 661.8371 |

C

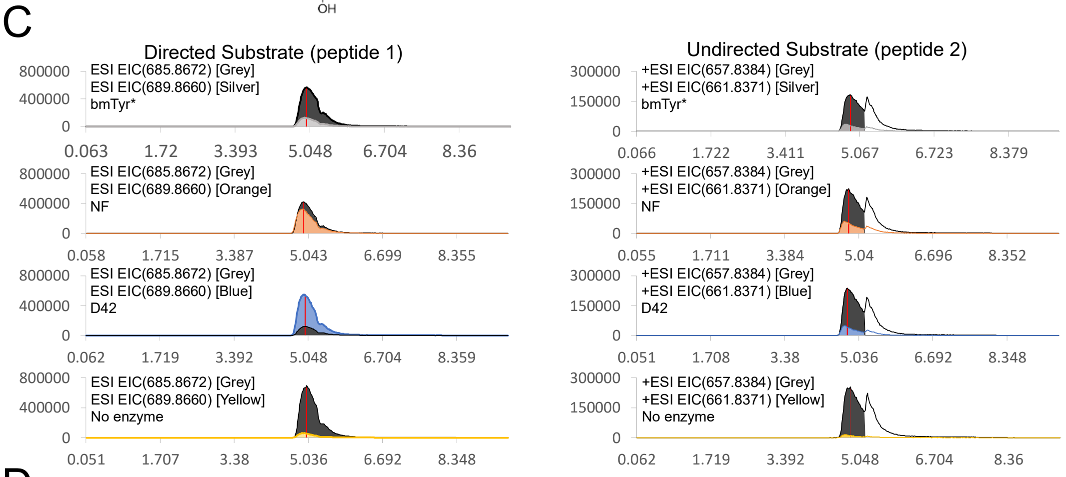

D

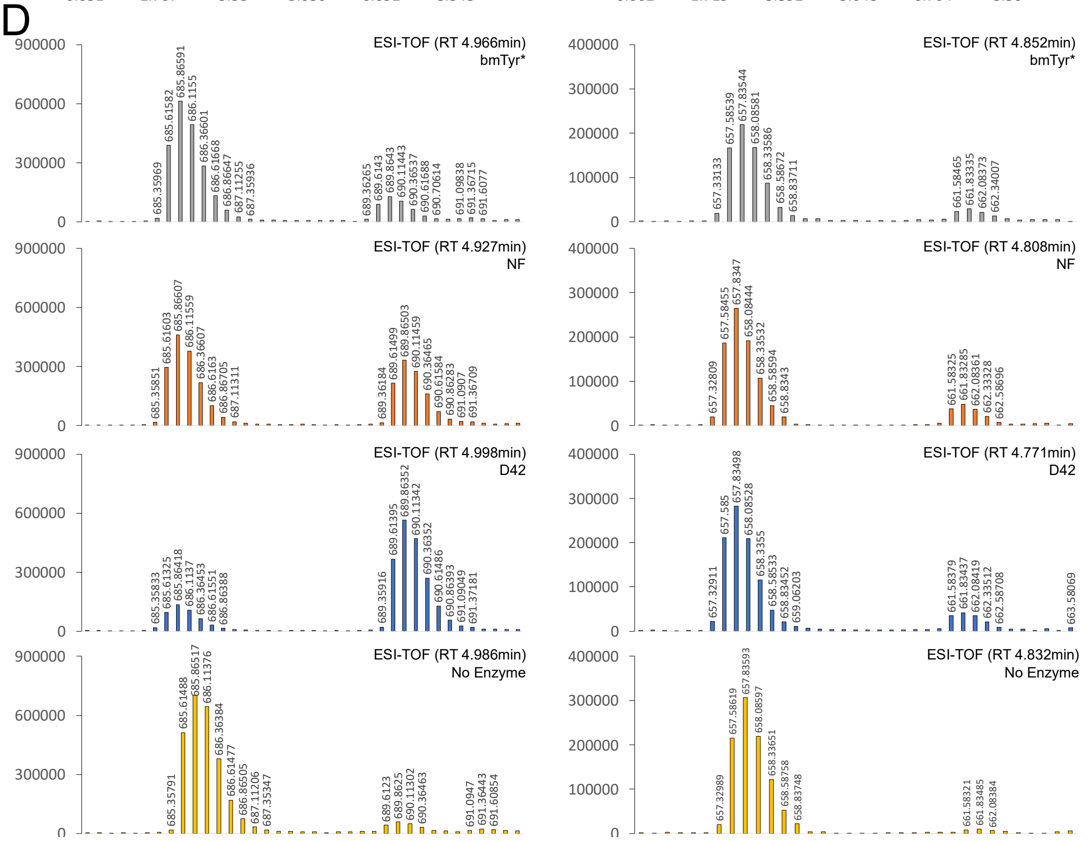

**Figure S2.** Representative EICs for directed vs undirected substrate experiments after 2min reactions. **A.** Structure of L-dopa product formed after incubation with enzyme in presence of ascorbic acid **B.** Expected substrate and product masses for the directed (peptide 1) and undirected (peptide 2) peptides. **C.** EICs for reactions with directed substrate and undirected substrate. Substrate EICs (grey) are overlayed with EIC's for product formed by bmTyr* (silver), NF (orange), D42 (blue) or the no-enzyme control (yellow). All EICs were determined using a standard error cutoff of 10ppm. **D.** Mass spectra for each reaction at the retention time indicated by the red bars in panel C respectively.

A

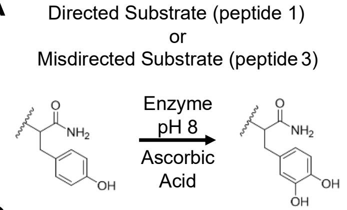

B

| Directed Substrate (Peptide 1) | | |
|---|---|---|
| APPLPPRNRPRLGSGSGSGSKNFLD**Y** | | |
| **Charge** | **Substrate** | **Product** |
| 0 | 2739.4398 | 2755.4348 |
| [M+4]+4 | 685.8672 | 689.8660 |

| Misdirected Substrate (Peptide 3) | | |
|---|---|---|
| **Y**DLFNKGSGSGSGSGSGSGSGSAPPLPPRNRPRL | | |
| **Charge** | **Substrate** | **Product** |
| 0 | 3331.64872 | 3331.64872 |
| [M+4]+4 | 829.92073 | 833.91946 |

C

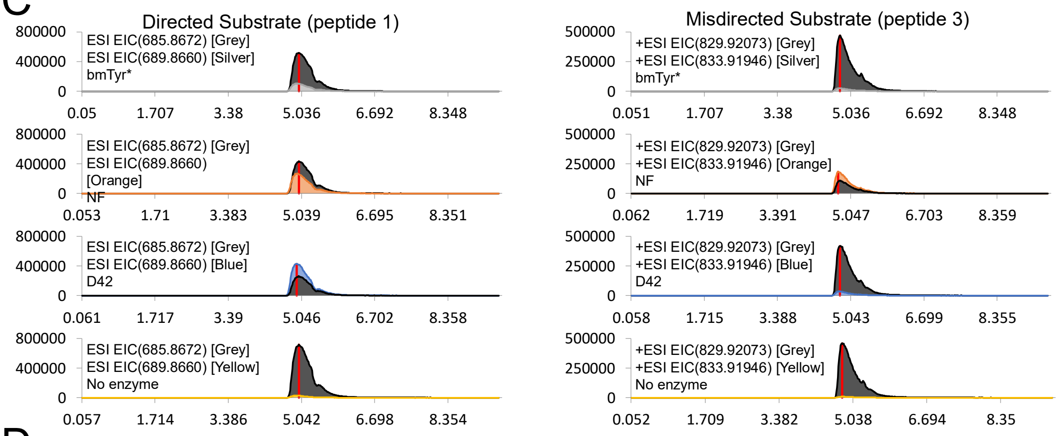

D

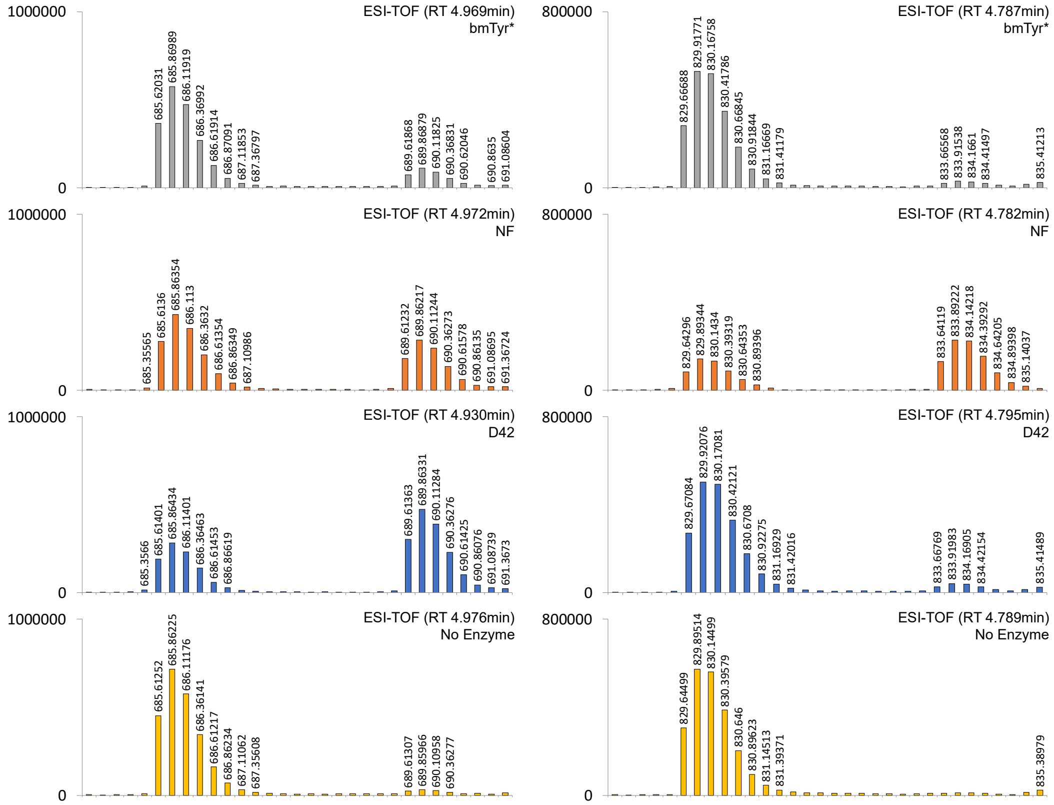

**Figure S3.** Representative EICs for directed vs misdirected substrate experiments after 2min reactions. **A.** Structure of L-dopa product formed after incubation with enzyme in presence of ascorbic acid **B.** Expected substrate and product masses for the directed (peptide 1) and misdirected (peptide 3) peptides. **C** EICs for reactions with directed substrate and undirected substrate. Substrate EICs (grey) are overlayed with EIC's for product formed by bmTyr* (silver), NF (orange), D42 (blue) or the no-enzyme control (yellow). All EICs were determined using a standard error cutoff of 10ppm. **D.** Mass spectra for each reaction at the retention time indicated by the red bars in panel C respectively.

A

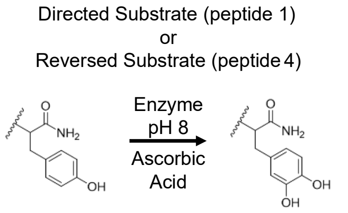

B

| Directed Substrate (Peptide 1) | | |
|---|---|---|
| APPLPPRNRPRLGSGSGSGSKNFLD**Y** | | |
| **Charge** | **Substrate** | **Product** |
| 0 | 2739.4398 | 2755.4348 |
| $[M+3]^{+3}$ | 914.15389 | 919.48552 |

| Reversed Substrate (Peptide 4) | | |
|---|---|---|
| YDLFNKGSGSGSGSGSVSLARRPLPPLP | | |
| **Charge** | **Substrate** | **Product** |
| 0 | 2815.48103 | 2831.47594 |
| $[M+3]^{+3}$ | 939.50095 | 944.83259 |

C

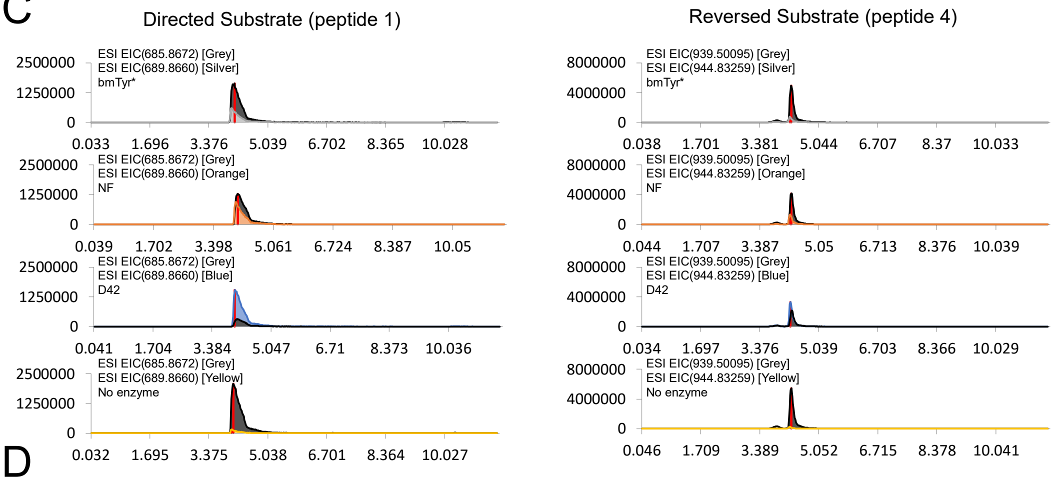

D

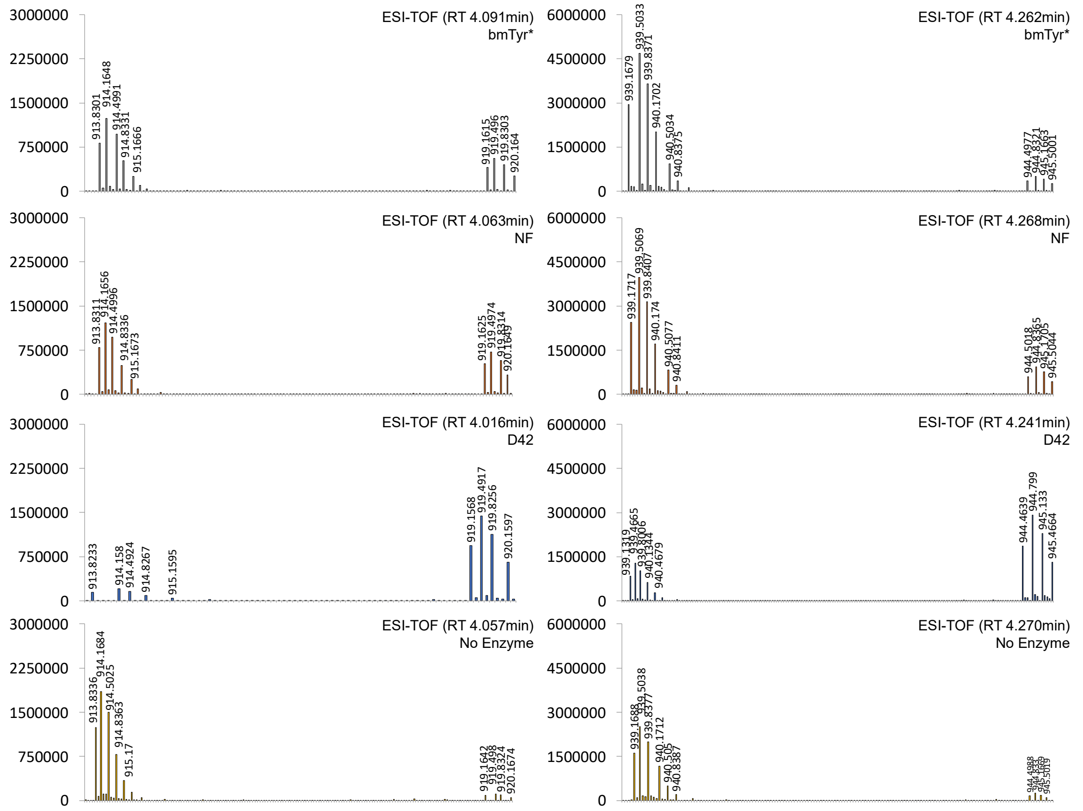

**Figure S4.** Representative EICs for directed vs reversed substrate experiments after 2min reactions. **A.** Structure of L-dopa product formed after incubation with enzyme in presence of ascorbic acid **B.** Expected substrate and product masses for the directed (peptide 1) and reversed (peptide 4) peptides. **C** EICs for reactions with directed substrate and undirected substrate. Substrate EICs (grey) are overlayed with EIC's for product formed by bmTyr* (silver), NF (orange), D42 (blue) or the no-enzyme control (yellow). All EICs were determined using a standard error cutoff of 100ppm. **D.** Mass spectra for each reaction at the retention time indicated by the red bars in panel C respectively.

A

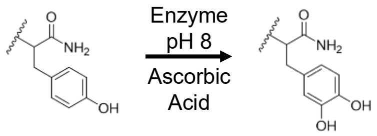

B

| Directed Substrate (Peptide 1) APPLPPRNRPRLGSGSGSGSKNFLD**Y** | | |
|---|---|---|
| **Charge** | **Substrate** | **Product** |
| 0 | 2739.4398 | 2755.4348 |
| $[M+4]^{+4}$ | 685.8672 | 689.8660 |

| Shortened Substrate (Peptide 5) APPLPPRNRPRLKNFLD**Y** | | |
|---|---|---|
| **Charge** | **Substrate** | **Product** |
| 0 | 2163.22587 | 2179.22078 |
| $[M+3]^{+3}$ | 722.0826 | 727.4142 |

C

D

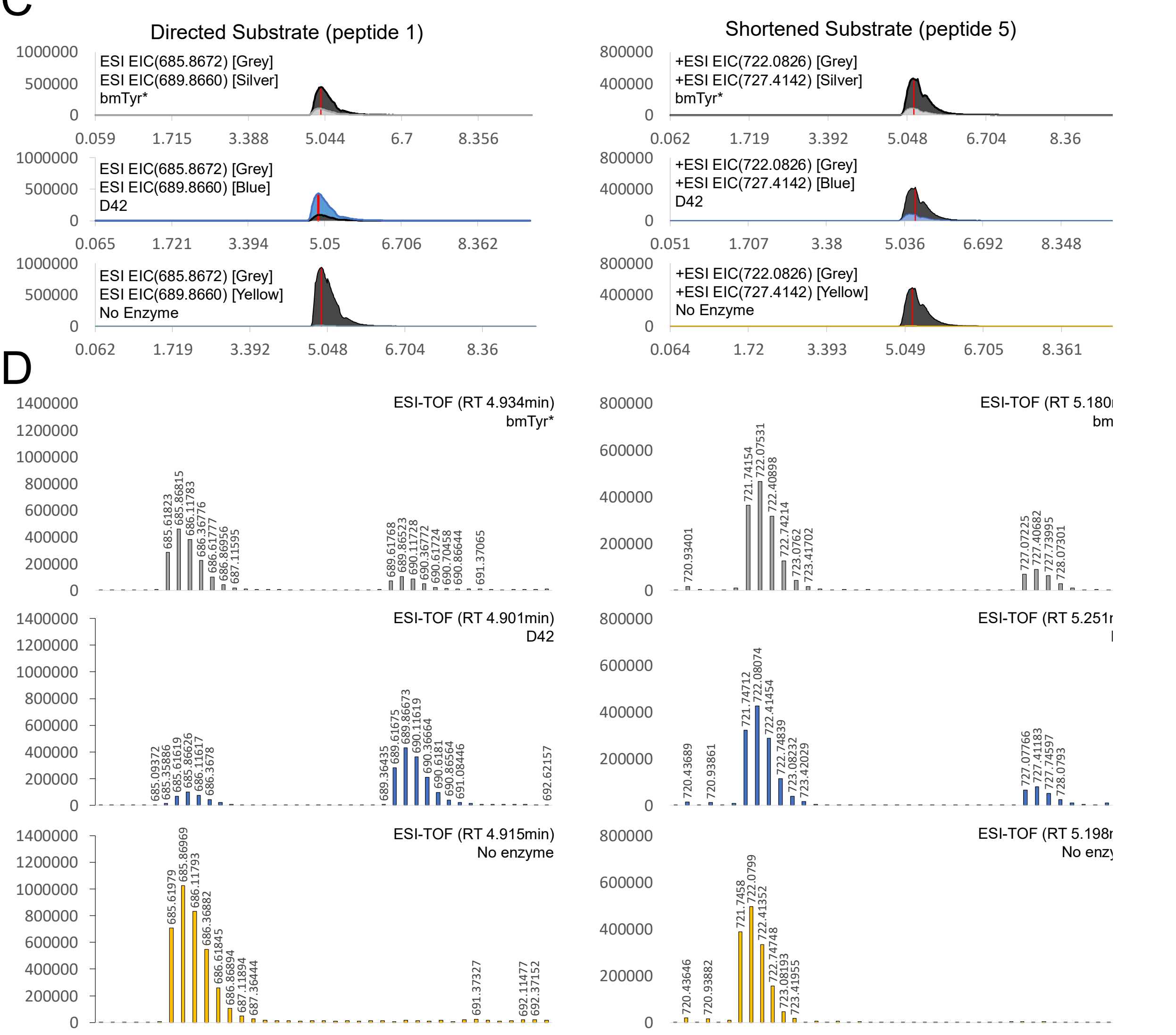

**Figure S5.** Representative EICs for directed vs shortened substrate experiments after 2min reactions. **A.** Structure of L-dopa product formed after incubation with enzyme in presence of ascorbic acid **B.** Expected substrate and product masses for the directed (peptide 1) and shortened (peptide 5) peptides. **C** EICs for reactions with directed substrate and undirected substrate. Substrate EICs (grey) are overlayed with EIC's for product formed by bmTyr* (silver), NF (orange), D42 (blue) or the no-enzyme control (yellow). All EICs were determined using a standard error cutoff of 10ppm. **D.** Mass spectra for each reaction at the retention time indicated by the red bars in panel C respectively.

**A**

Easy Substrate (peptide 6)
Medium Substrate (peptide 1)
Or
Hard Substrate (peptide 7)

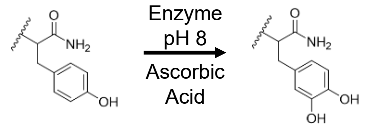

**B**

**Medium Substrate (Peptide 1)**

APPLPPRNRPRLGSGSGSGSKNFLD**Y**

| Charge | Substrate | Product |
|---|---|---|
| 0 | 2739.4398 | 2755.4348 |
| $[M+4]^{+4}$ | 685.8672 | 689.8660 |

**Easy Substrate (Peptide 6)**

APPLPPRNRPRLGSGSGSGSGSGSG**Y**

| Charge | Substrate | Product |
|---|---|---|
| 0 | 2467.25097 | 2483.24589 |
| $[M+3]^{+3}$ | 823.42427 | 828.75590 |

**Hard Substrate (Peptide 7)**

EWAPPLPPRNRPRRSGGSGGSGGSGGSKET**Y**SK

| Charge | Substrate | Product |
|---|---|---|
| 0 | 3368.69158 | 3384.68650 |
| $[M+5]^{+5}$ | 674.7456 | 677.9446 |

**C**

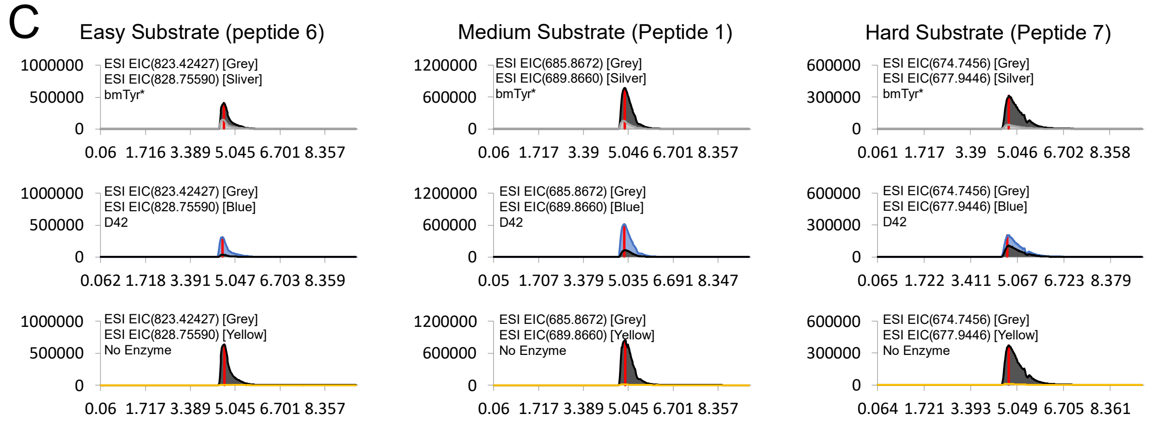

**D**

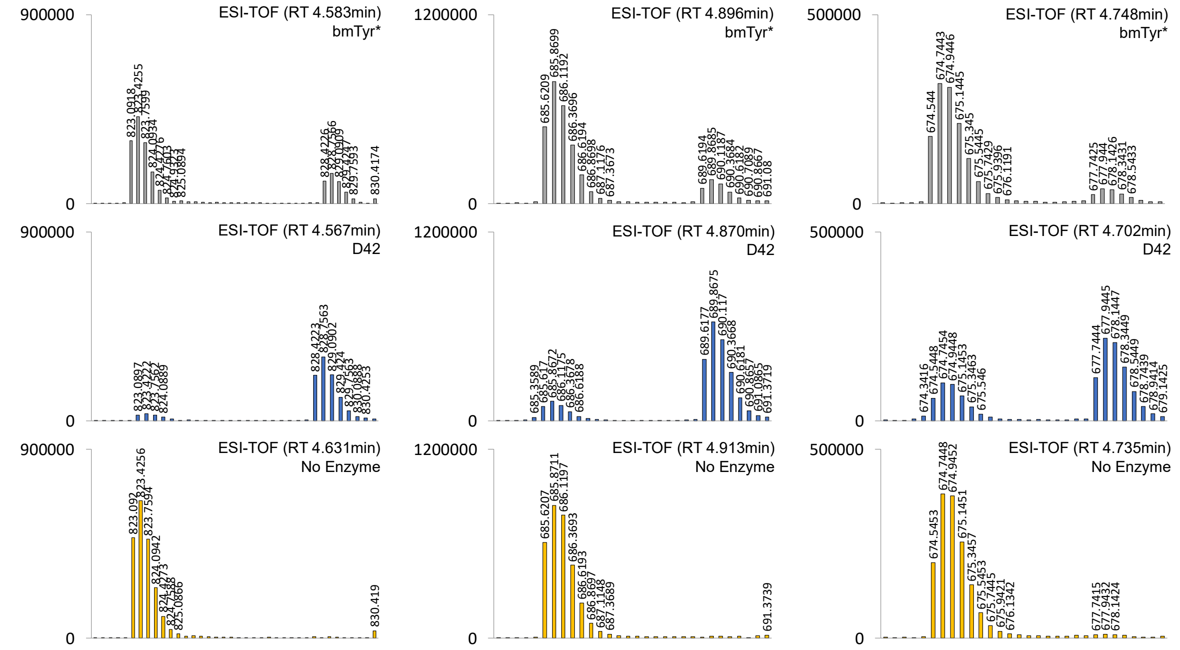

**Figure S6.** Representative EICs for easy, directed or hard substrate experiments after 2min reactions. **A.** Structure of L-dopa product formed after incubation with enzyme in presence of ascorbic acid **B.** Expected substrate and product masses for the easy (peptide 6), directed (peptide 1) and hard (peptide 7) peptides. **C** EICs for reactions with easy, directed, and hard substrates. Substrate EICs (grey) are overlayed with EIC's for product formed by bmTyr* (silver), NF (orange), D42 (blue) or the no-enzyme control (yellow). All EICs were determined using a standard error cutoff of 10ppm. **D.** Mass spectra for each reaction at the retention time indicated by the red bars in panel C respectively.

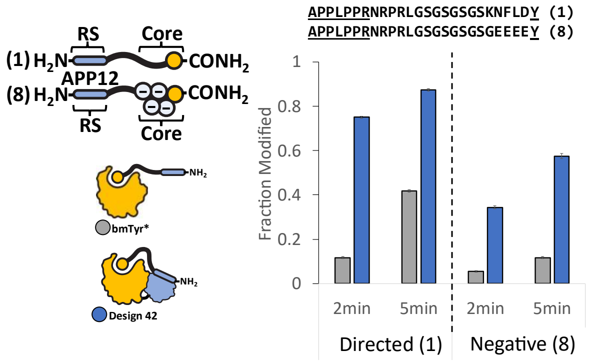

**Figure S7** Activity Against Negatively Charged Substrate. 50nM of bmTyr* [grey] or D42 [blue] was reacted against 10uM of directed (peptide 1) or negatively charged (peptide 8) peptides in 50mM phosphate buffer pH 8, 12.5mM ascorbic acid and 1uM CuSO4 for indicated times. Fraction modified is calculated as area under curve for hydroxylated peptide over sum of substrate and hydroxylated peptide as measured using MALDI. Bars show mean of triplicate wells and error bars indicate S.D

A

Directed Substrate (peptide 1)
And
Undirected Substrate (peptide 2)

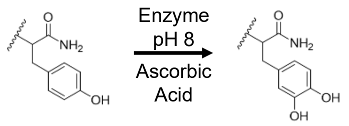

B

| Directed Substrate (Peptide 1) | | | Undirected Substrate (Peptide 2) | | |
|---|---|---|---|---|---|
| APPLPPRNRPRLGSGSGSGSKNFLD**Y** | | | AKAARSAEAKEAGRGSGSGSKNFLD**Y** | | |
| **Charge** | **Substrate** | **Product** | **Charge** | **Substrate** | **Product** |
| 0 | 2739.4398 | 2755.4348 | 0 | 2627.3245 | 2641.3402 |
| $[M+4]^{+4}$ | 685.8672 | 689.8660 | $[M+4]^{+4}$ | 657.8384 | 661.8371 |

C

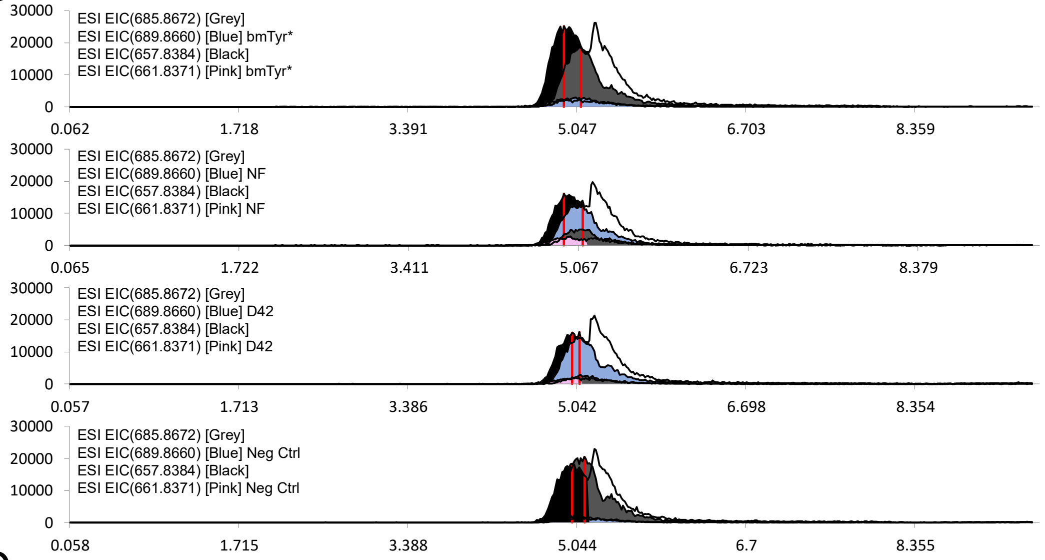

D

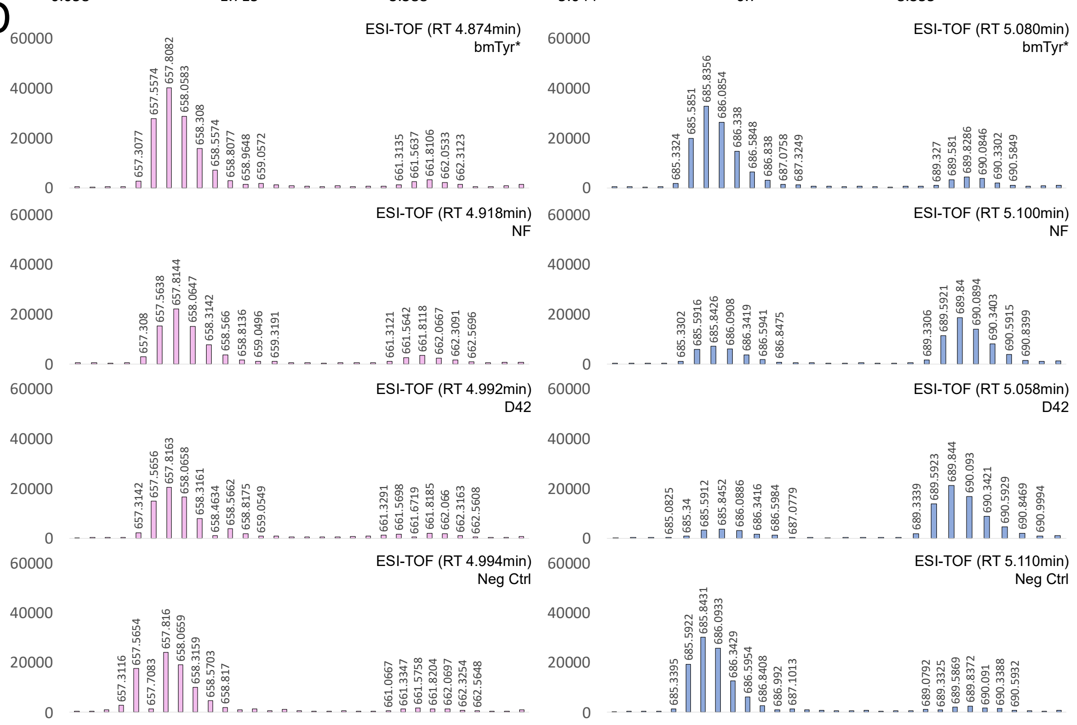

**Figure S8** Representative EICs for coincubation of directed and undirected substrate at 1uM each (2uM total). **A**. Structures and **B**. expected masses for the most abundant ions of directed (peptide 1) and undirected (peptide 2) peptides. **C.** EICs for directed substrate (dark grey) and product (light blue), and undirected substrate (black) and product (light pink). Fraction modified was calculated as the area under the curve for the product EIC divided by the sum of the areas under the curve for both product and substrate EICs. All EICs were determined using a standard error cutoff of 10ppm. **D.** Mass spectra for each reaction at the retention time indicated by the red bars in panel C

A

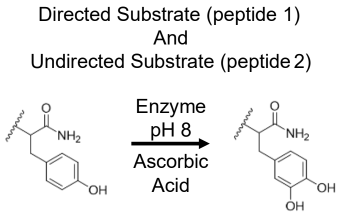

B

| Directed Substrate (Peptide 1) | | |
|---|---|---|
| APPLPPRNRPRLGSGSGSGSGSKNFLD**Y** | | |
| **Charge** | **Substrate** | **Product** |
| 0 | 2739.4398 | 2755.4348 |
| $[M+4]^{+4}$ | 685.8672 | 689.8660 |

| Undirected Substrate (Peptide 2) | | |
|---|---|---|
| AKAARSAEAKEAGRGSGSGSKNFLD**Y** | | |
| **Charge** | **Substrate** | **Product** |
| 0 | 2627.3245 | 2641.3402 |
| $[M+4]^{+4}$ | 657.8384 | 661.8371 |

C

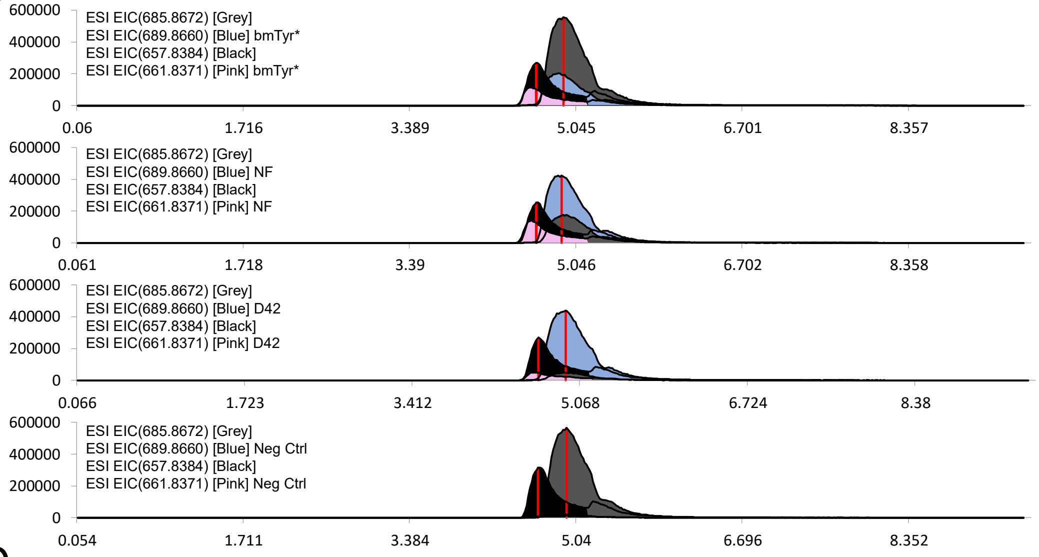

D

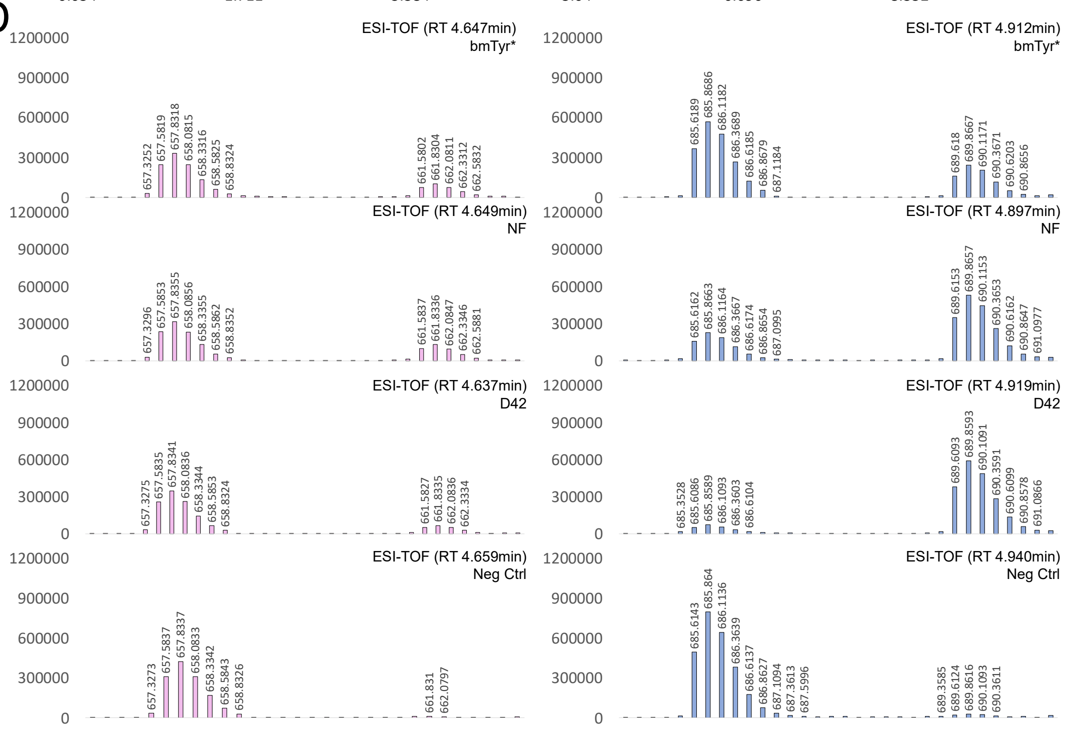

**Figure S9.** Representative EICs for coincubation of directed and undirected substrate at 10uM each (20uM total). **A**. Structures and **B**. expected masses for the most abundant ions of directed (peptide 1) and undirected (peptide 2) peptides. **C.** EICs for directed substrate (dark grey) and product (light blue), and undirected substrate (black) and product (light pink). Fraction modified was calculated as the area under the curve for the product EIC divided by the sum of the areas under the curve for both product and substrate EICs. All EICs were determined using a standard error cutoff of 10ppm. **D.** Mass spectra for each reaction at the retention time indicated by the red bars in panel C

**Table S4**. Crystallography Data Collection and Refinement Statistics

| | Tyrosinase-SRD |
|---|---|
| **Data collection** | |
| Wavelength (Å) | 0.9686 |
| Space group | $P4_332$ |
| Unit Cell (Å/degrees) | 150.3, 150.3, 150.3 |
| Resolution range (Å)[1] | 37.6 - 2.6 (2.6 -2.61) |
| Total reflections | 506,390 |
| Unique reflections | 18,413 |
| Multiplicity | 27.5 (26.1) |
| Completeness (%) | 100.0 (100.0) |
| Mean I/sigma (I) | 23.9 (2.9) |
| R-merge (%)[2] | 12.8 (140.8) |
| R-pim (%) | 2.5 (27.9) |
| CC ½ | 1.000 (0.699) |
| **Refinement** | |
| Resolution (Å) | 25.0-2.6 |
| Number of reflections | 18,412 |
| R-work | 20.1 |
| R-free[3] | 24.6 |
| Number of atoms | |
| Macromolecules | 2827 |
| Solvent | 93 |
| Average B-factor | |
| Macromolecules | 61.9 |
| Solvent | 59.9 |
| RMS (bond lengths) | 0.007 |
| RMS (bond angles) | 1.994 |
| Molprobity clash score | 6.19 |
| Favored (%) | 93.02 |
| Allowed (%) | 5.52 |
| Outliers (%) | 1.45 |

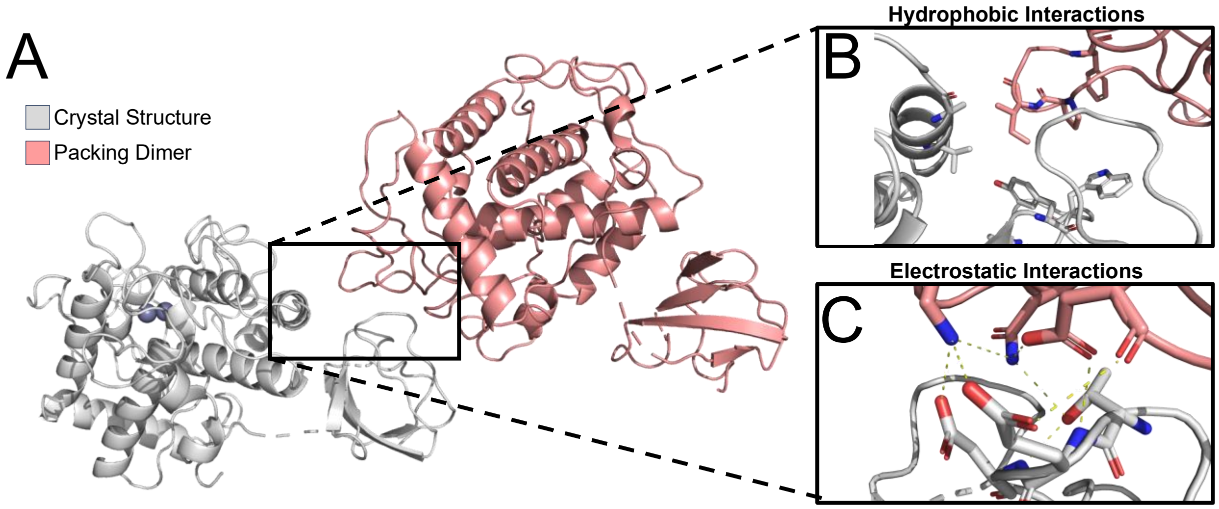

**Figure S10.** Crystal packing interaction between SH3 domain and tyrosinase domain of a D42 dimer. **A.** 2.6Å X-ray **c**rystal structure showing structure of D42 dimer. **B** Hydrophobic interactions between SH3 and tyrosinase domains at the D42 dimer interface. **C.** Charged and polar residues at D42 dimer interface within 3Å of each other

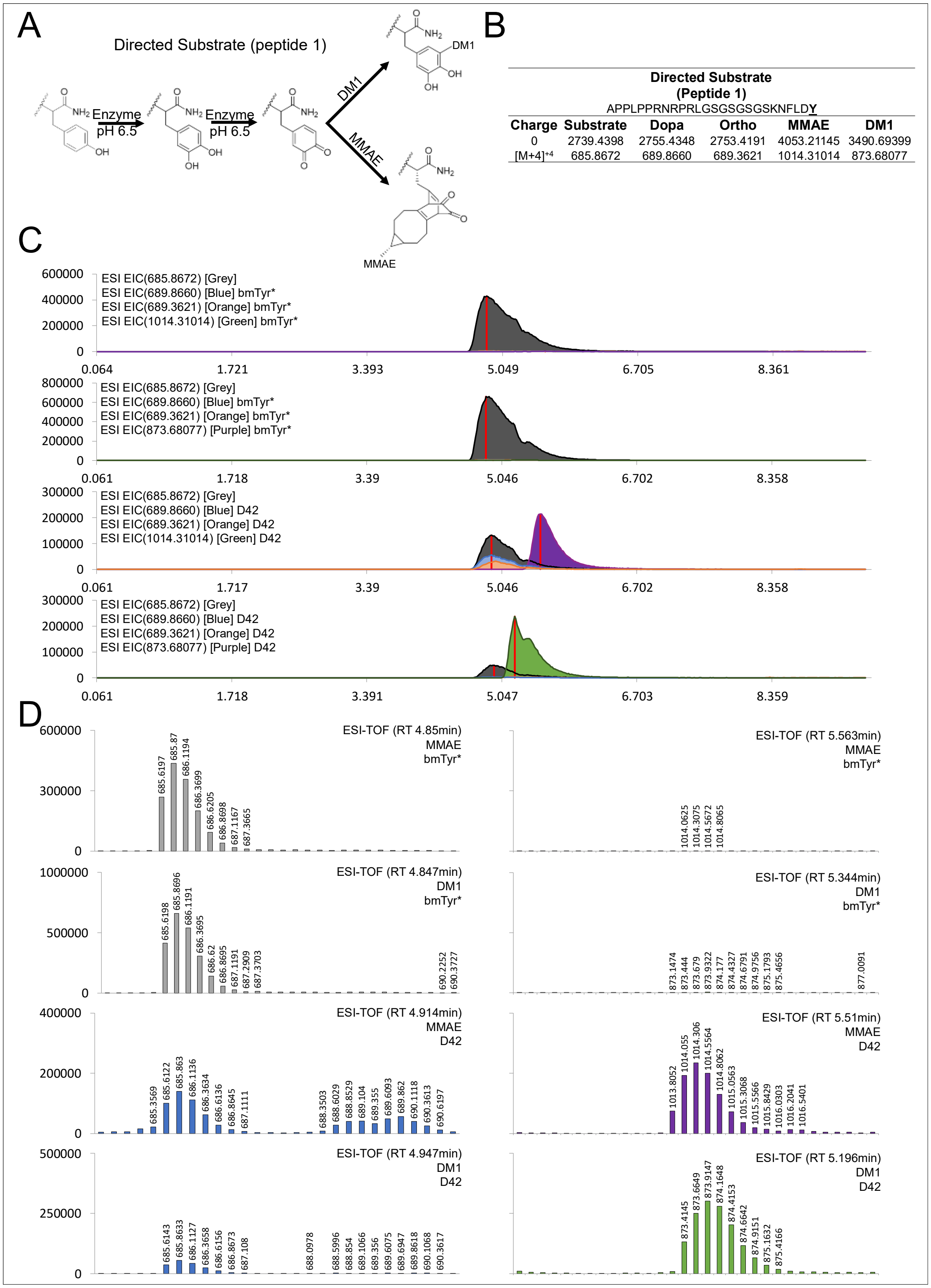

| Charge | Substrate | Dopa | Ortho | MMAE | DM1 |
|---|---|---|---|---|---|
| | Directed Substrate (Peptide 1) APPLPPRNRPRLGSGSGSGSKNFLDY | | | | |
| 0 | 2739.4398 | 2755.4348 | 2753.4191 | 4053.21145 | 3490.69399 |
| [M+4]$^{+4}$ | 685.8672 | 689.8660 | 689.3621 | 1014.31014 | 873.68077 |

**Figure S11** Representative EICs for conjugation reactions with directed substrate experiments after 2-minute reactions. **A.** Structures and **B.** expected masses for MMAE or DM1 conjugation to directed substrate peptides. **C.** EICs for substrate (black) conjugated to either MMAE (purple) or DM1 (green). EICs for dopa product (orange) and unreacted orthoquinone (orange) are also shown. All EICs were determined using a standard error cutoff of 10ppm. **D.** Mass spectra for each reaction at the retention time indicated by the red bars in panel C

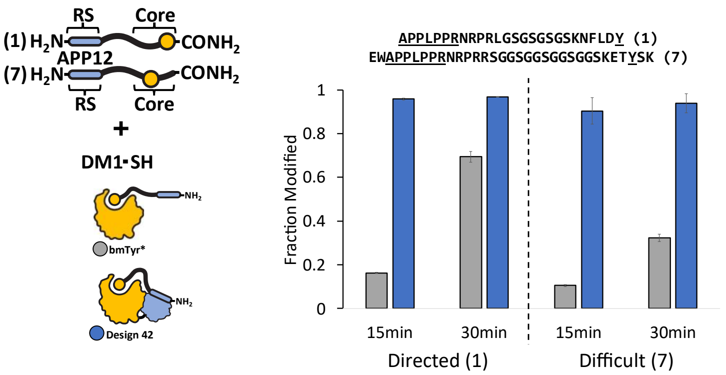

**Figure S12.** Extended DM1 conjugation reaction times with WT vs R209H mut enzyme systems. Medium (peptide 1) and difficult (peptide 7) peptide substrates were incubated together with 50uM DM1 and either 250nM enzyme (bmTyr* [grey] and D42 [blue]) for indicated times. All reactions were ran at 50mM phosphate buffer at 6.5 and 25ºC and enzyme reactions quenched with 4mM Tropolone before running on LCMS. Error bars represent standard deviation across three replicates centered about the mean.

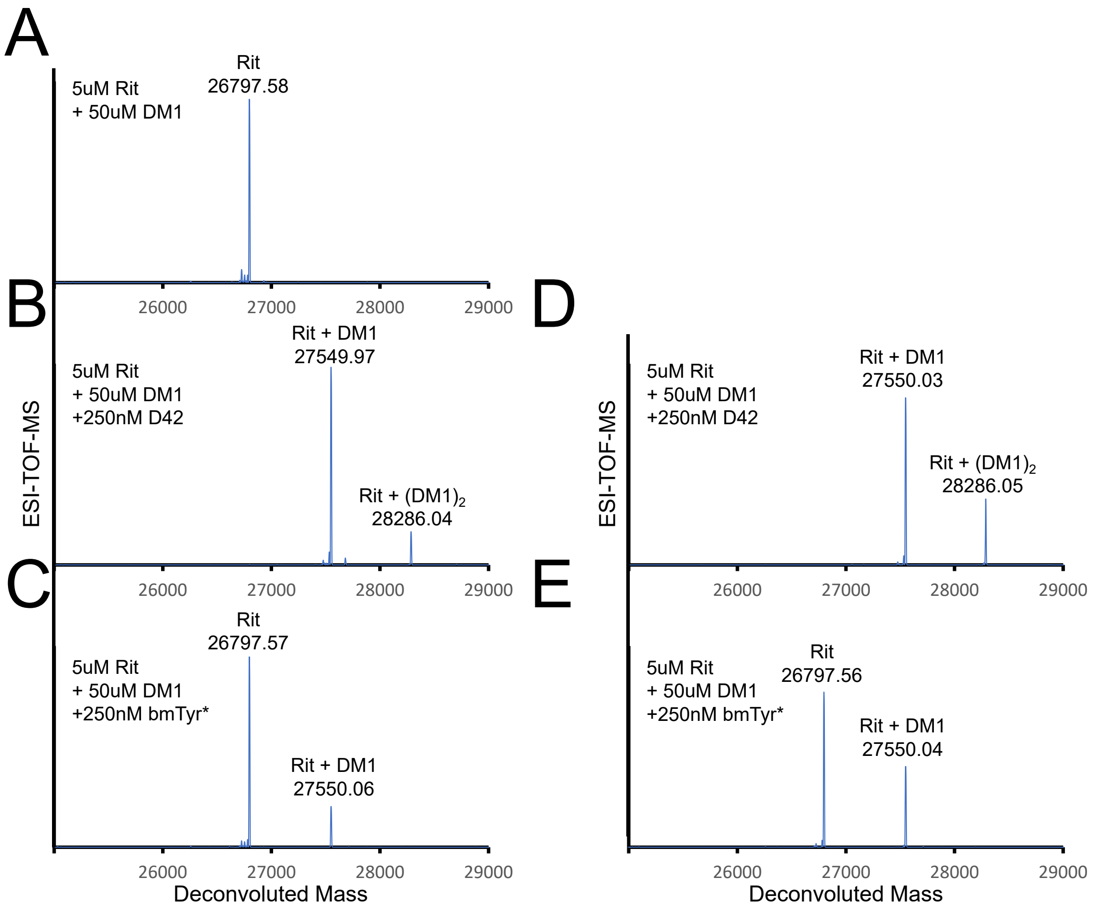

**Figure S13.** Extended antibody drug conjugation reaction times. **A.** Conjugation strategy for installing DM1 onto antibody using **B.** no enzyme, **C. and E.** D42, or **D. and F.** bmTyr*. Reactions were performed for 15min **C** and **D.** and 30min **E.** and **F.** Reactions were performed at 25ºC and quenched with 4mM Tropolone. All reactions were ran at 50mM phosphate buffer at pH 6.5 and 25ºC and reduced with 10mM DTT prior to submitting for intact MS.

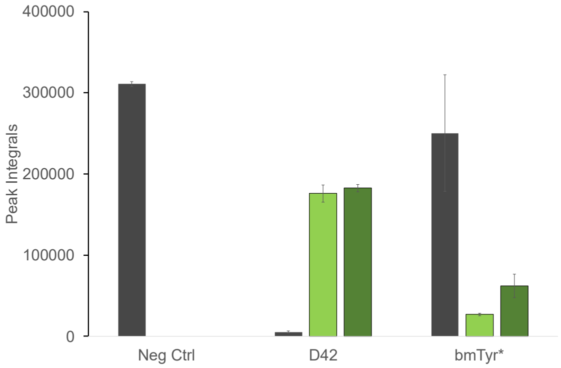

**Figure S14.** Installation of DM1 onto Rituximab modification tag. 5uM Rit was incubated with either no enzyme (Neg Ctrl) or 250nM enzyme (D42 or bmTyr*) and 1uM $CuSO_4$ for 15min. Modification tag was cleaved by incubating quenched reactions with 7.5uM Prescission protease for 15min. Bars show peak integrals for unmodified cleavage product (grey), DM1 (light green) and doubly incorporated DM1 [$(DM1)_2$] (dark green) as determined by LCMS. All reactions were ran at 50mM phosphate buffer at 6.5 and 25°C and enzyme reactions quenched with 4mM Tropolone. Error bars represent standard deviation across two replicates centered about the mean.

**Directed Substrate [P4527] (Peptide 1),** APPLPPRNRPRLGSGSGSGSKNFLDY-$NH_2$, MW 2740.09

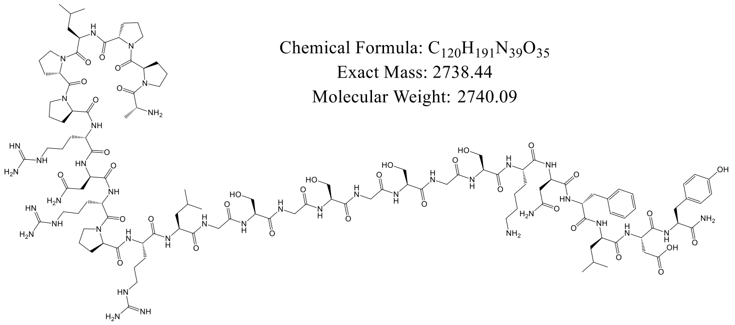

**MALDI**

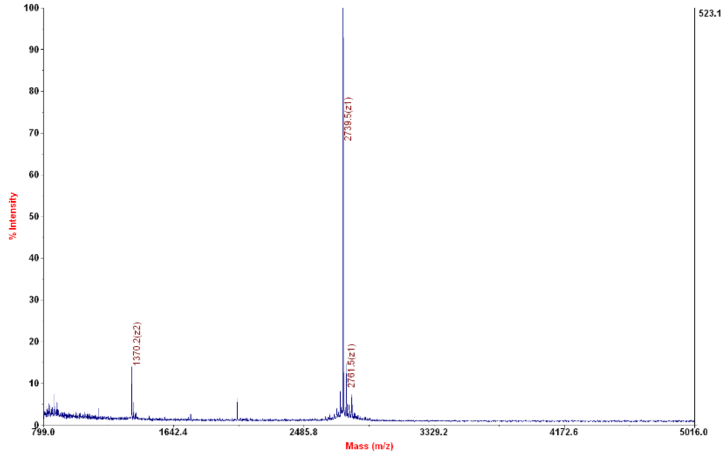

**HPLC**

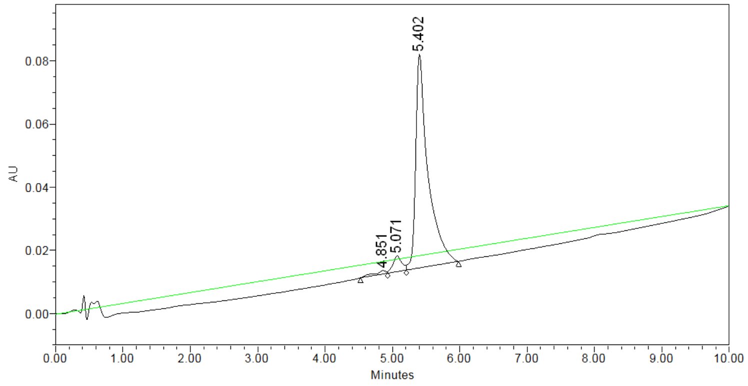

| | RT | % Area | Area | Height |
|---|---|---|---|---|
| 1 | 4.851 | 1.46 | 13390 | 1091 |
| 2 | 5.071 | 4.92 | 45260 | 4939 |
| 3 | 5.402 | 93.62 | 860644 | 67649 |

**Figure S15** MALDI and HPLC of directed substrate APPLPPRNRPRLGSGSGSGSKNFLDY-$NH_2$

**Undirected Substrate [P4634 I] (Peptide 2)**, AKAARSAEAKEAGRGSGSGSKNFLDY-$NH_2$, MW 2627.86

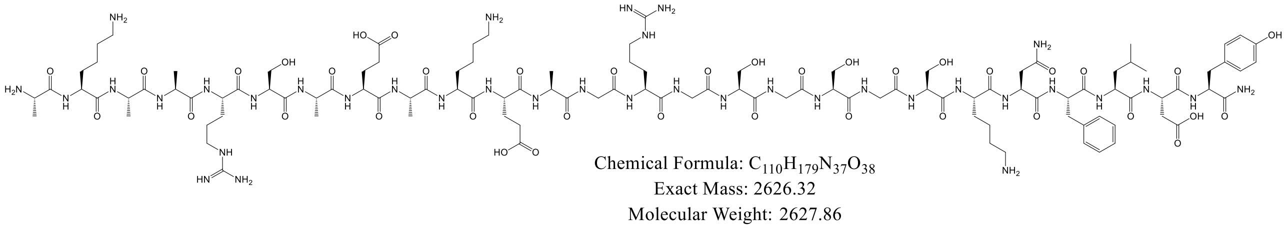

**MALDI**

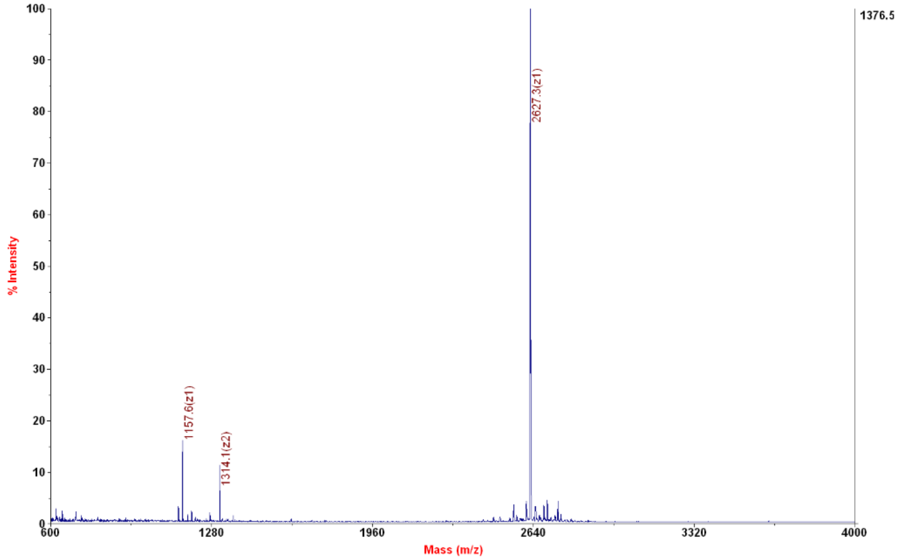

**HPLC**

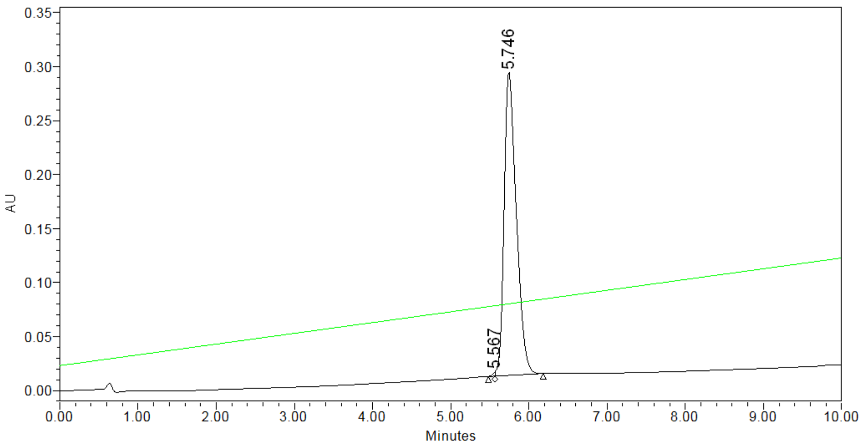

| | RT | % Area | Area | Height |
|---|---|---|---|---|
| 1 | 5.567 | 0.24 | 7170 | 3996 |
| 2 | 5.746 | 99.76 | 3039282 | 280666 |

**Figure S16** MALDI and HPLC of undirected substrate AKAARSAEAKEAGRGSGSGSKNFLDY-$NH_2$

**Misdirected Substrate [P4541 I] (Peptide 3)**, YDLFNKGSGSGSGSGSGSGSGSAPPLPPRNRPRL-$NH_2$, MW 3316.61

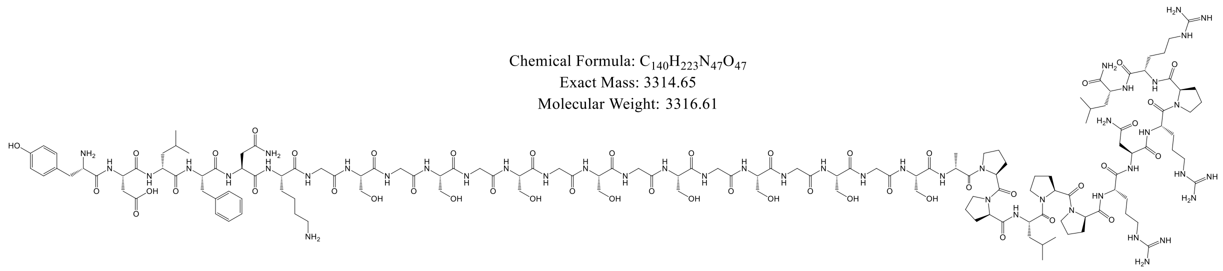

**MALDI**

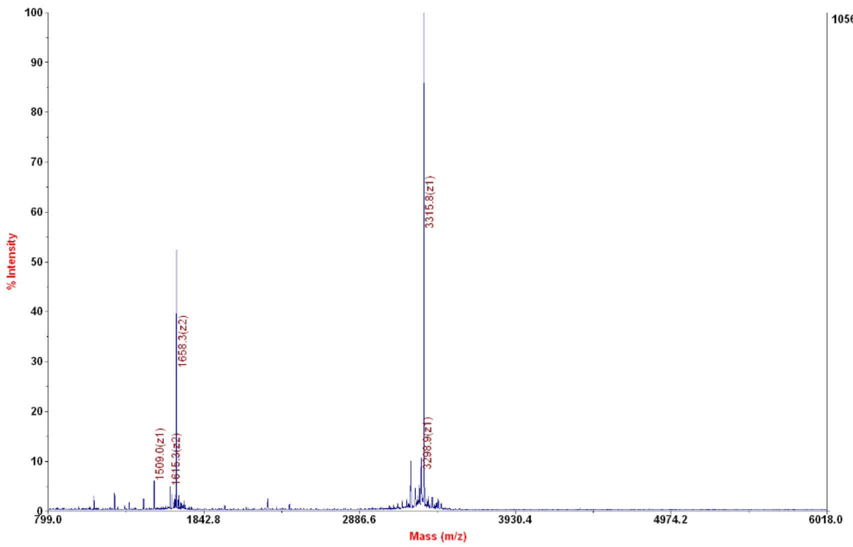

**HPLC**

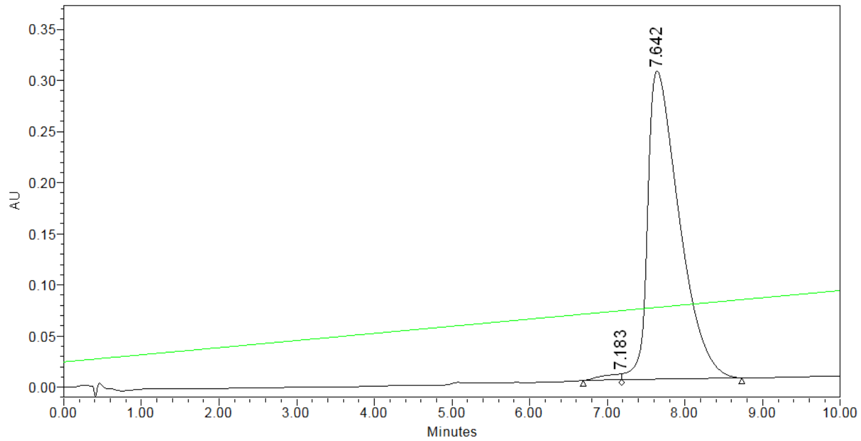

| | RT | % Area | Area | Height |
|---|---|---|---|---|
| 1 | 7.183 | 1.15 | 98896 | 5835 |
| 2 | 7.642 | 98.85 | 8527926 | 301329 |

**Figure S17** MALDI and HPLC of misdirected substrate YDLFNKGSGSGSGSGSGSGS-APPLPPRNRPRL-$NH_2$

**Reversed Substrate [P4531] (Peptide 4),** YDLFNKGSGSGSGSGSVSLARRPLPPLP-$NH_2$, MW 2816.18

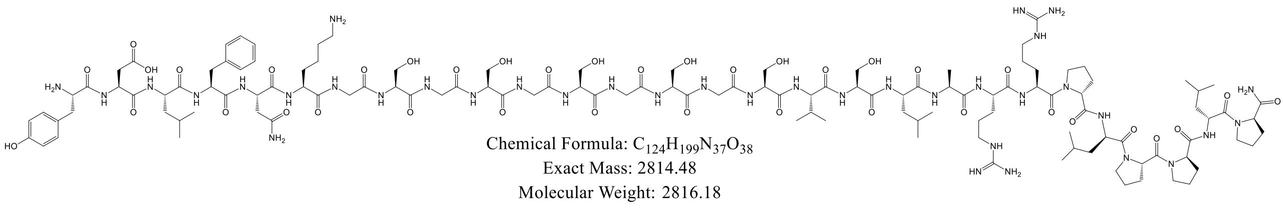

**MALDI**

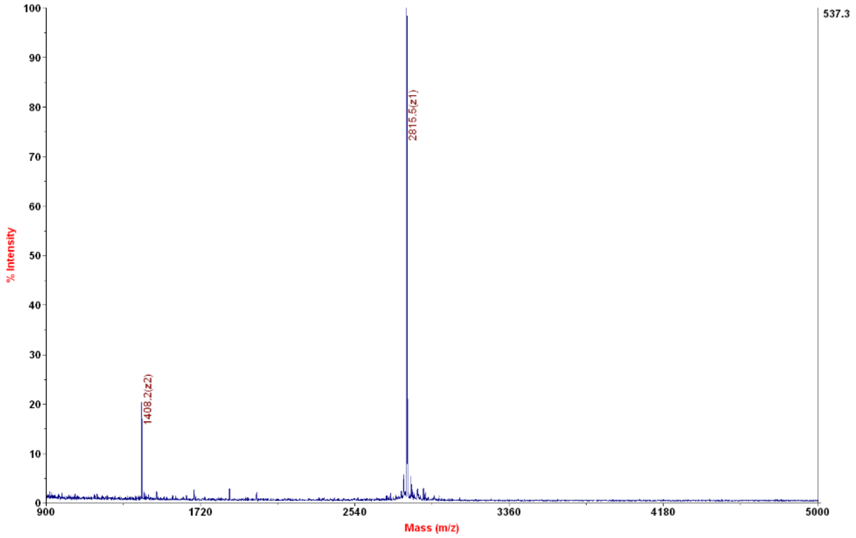

**HPLC**

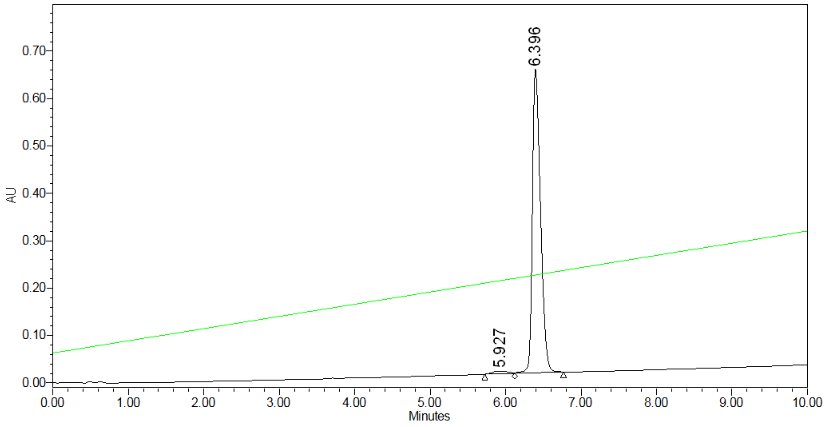

| | RT | % Area | Area | Height |
|---|---|---|---|---|
| 1 | 5.927 | 1.63 | 78930 | 5594 |
| 2 | 6.396 | 98.37 | 4776121 | 643393 |

**Figure S18** MALDI and HPLC of reversed substrate YDLFNKGSGSGSGSGSVSLARRPLPPLP-$NH_2$

**Shortened Substrate [P4528] (Peptide 5),** APPLPPRNRPRLKNFLDY-$NH_2$, MW 2163.0

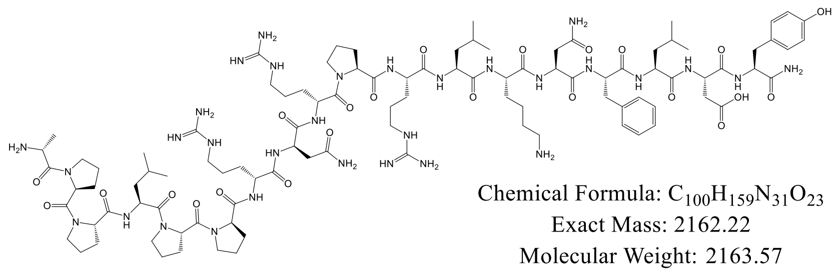

**MALDI**

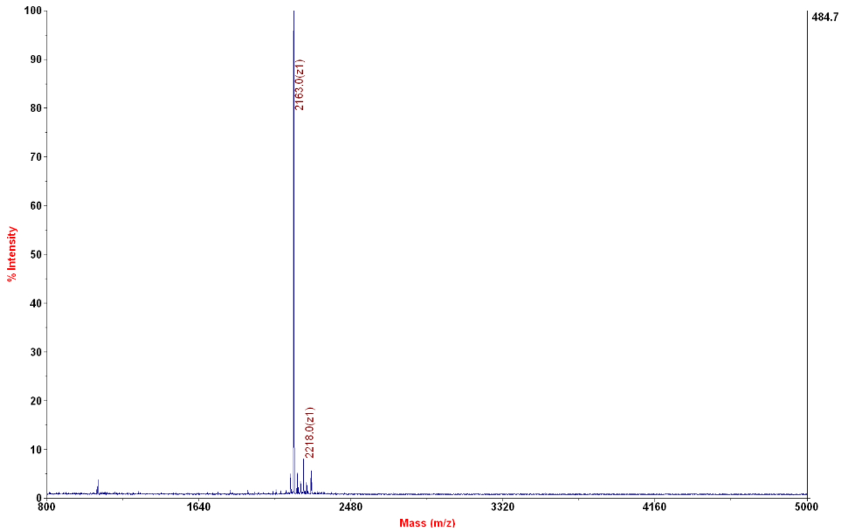

**HPLC**

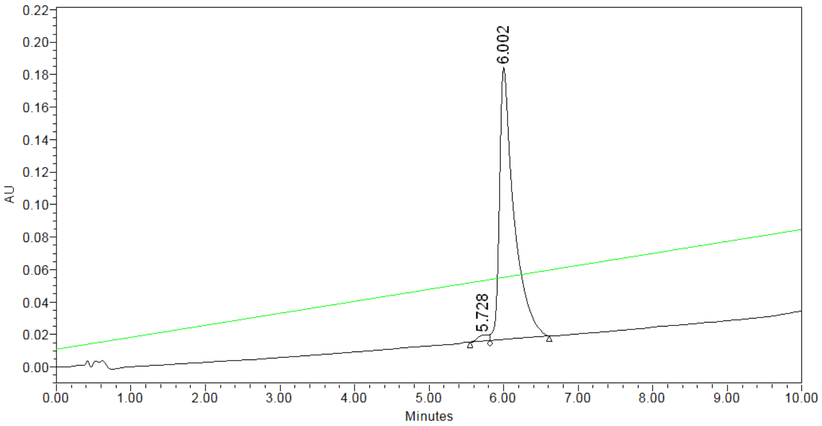

| | RT | % Area | Area | Height |
|---|---|---|---|---|
| 1 | 5.728 | 1.66 | 38114 | 3676 |
| 2 | 6.002 | 98.34 | 2252019 | 167534 |

**Figure S19** MALDI and HPLC of shortened substrate APPLPPRNRPRLKNFLDY-$NH_2$

**Easy Substrate [P4540 I] (Peptide 6)**, APPLPPRNRPRLGSGSGSGSGSGSGY-$NH_2$, MW 2467.69

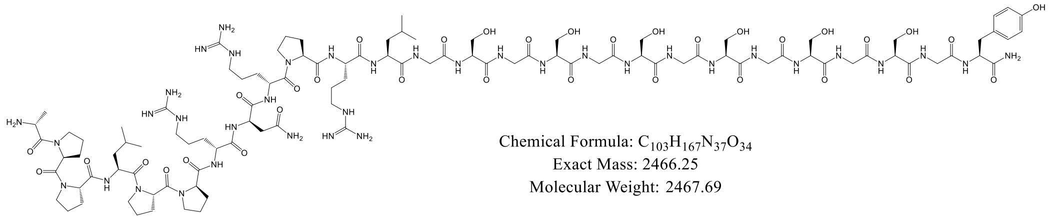

**MALDI**

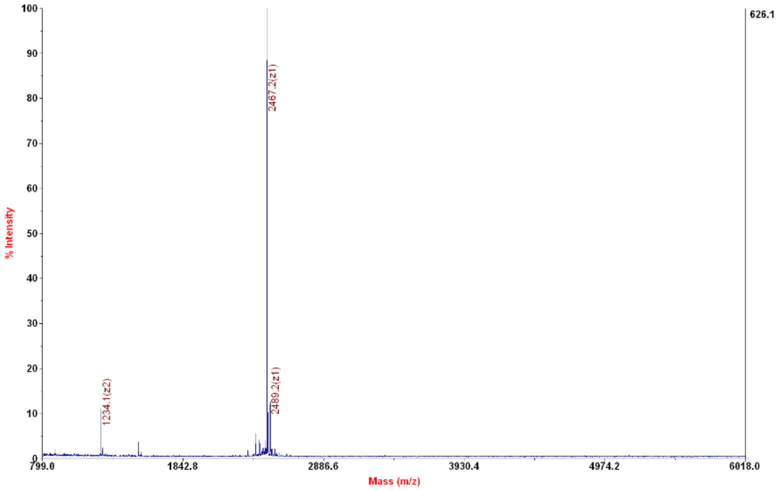

**HPLC**

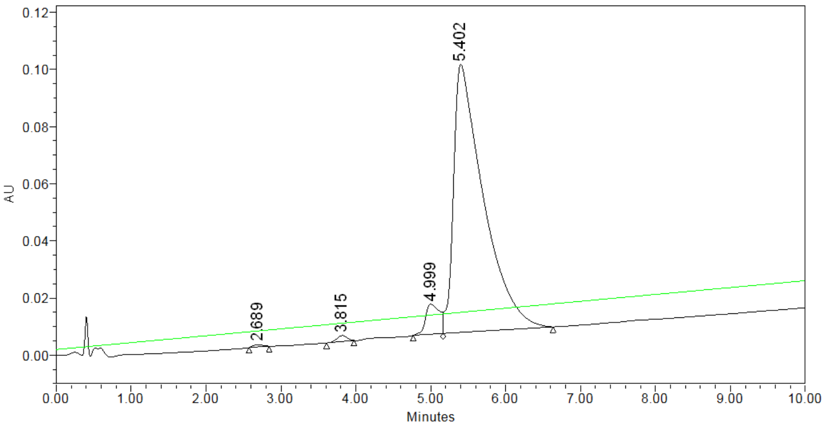

| | RT | % Area | Area | Height |
|---|---|---|---|---|
| 1 | 2.689 | 0.24 | 6584 | 709 |
| 2 | 3.815 | 0.71 | 19494 | 2066 |
| 3 | 4.999 | 5.03 | 138903 | 10552 |
| 4 | 5.402 | 94.03 | 2598526 | 93791 |

**Figure S20** MALDI and HPLC of easy substrate APPLPPRNRPRLGSGSGSGSGSGSGY-$NH_2$

**Difficult Substrate [P4482 I] (Peptide 7),** EWAPPLPPRNRPRRSGGSGGSGGSGGSKETYSK-$NH_2$, MW 3369.67

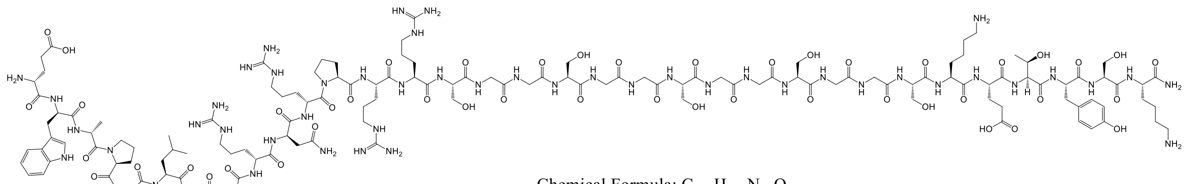

**MALDI**

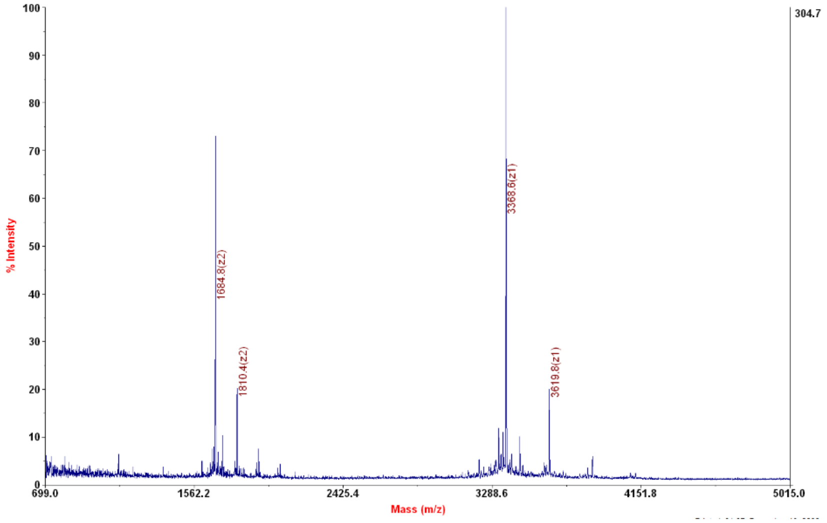

**HPLC**

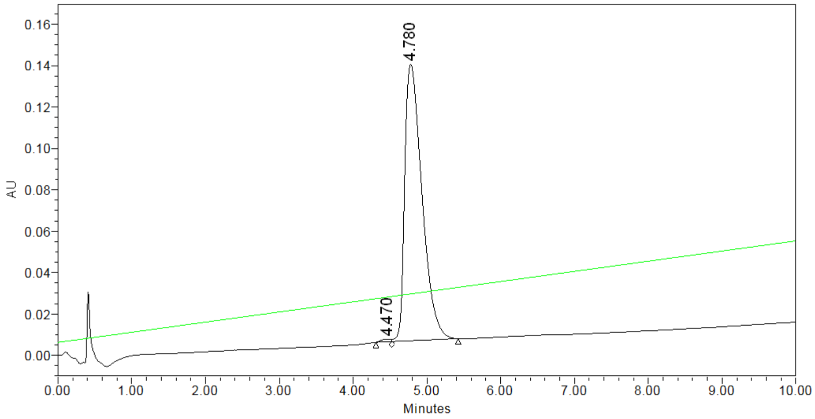

| | RT | % Area | Area | Height |
|---|---|---|---|---|
| 1 | 4.470 | 0.52 | 11357 | 1194 |
| 2 | 4.780 | 99.48 | 2171148 | 133693 |

**Figure S21** MALDI and HPLC of difficult substrate EWAPPLPPRNRPRRSGGSGGSGGSGGSKETYSK-$NH_2$

**Negative Substrate [P4772 I] (Peptide 8),** APPLPPRNRPRLGSGSGSGSGEEEEY-$NH_2$, MW 2695.91

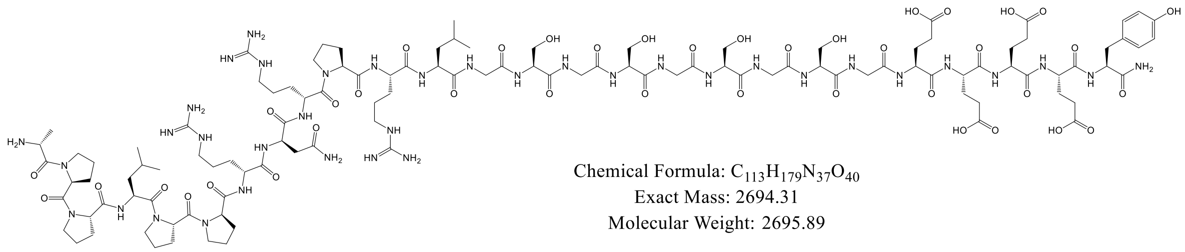

**MALDI**

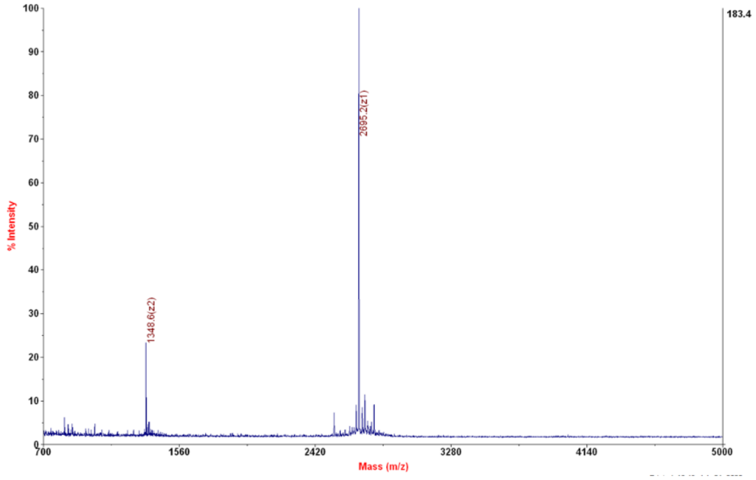

**HPLC**

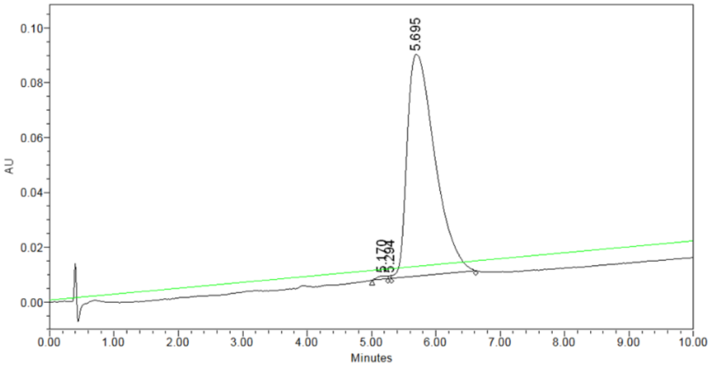

| | RT | % Area | Area | Height |
|---|---|---|---|---|
| 1 | 5.170 | 0.50 | 12156 | 1220 |
| 2 | 5.294 | 0.12 | 2836 | 1018 |
| 3 | 5.695 | 99.39 | 2424267 | 81013 |

**Figure S22** MALDI and HPLC of negative substrate APPLPPRNRPRLGSGSGSGSGEEEEY -$NH_2$

**Design Protocol**

```
<JobDefinitionFile>
  <Job>
    <Input>
      <PDB listfile="4znx_inputs_list.txt" />
    </Input>
    <Output>
      <PDB path="output" />
    </Output>
  </Job>

  <Common>

    <SCOREFXNS>
        <ScoreFunction name="low_res_sfxn" weights="interchain_cen">
            <Reweight scoretype="atom_pair_constraint" weight="1"/>
            <Reweight scoretype="interchain_vdw" weight="100"/>
        </ScoreFunction>

        <ScoreFunction name="sfxn_high_res" weights="ref2015_cst"/>
        <ScoreFunction name="sfxn_basic_design" weights="ref2015">
            <Reweight scoretype="res_type_constraint" weight="1"/>
        </ScoreFunction>
        <ScoreFunction name="sfxn_basic" weights="ref2015"/>
    </SCOREFXNS>

    <RESIDUE_SELECTORS>
        <Chain chains="A" name="enzyme"/>
        <Chain chains="B" name="pep_binder"/>
        <Chain chains="C" name="peptide"/>

        Index name="pep_binder_face" resnums="38B"/>
        Index name="enzyme_center" resnums="227A"/>

        <Index name="A_designable"
resnums="13A,16A,17A,20A,27A,28A,30A,32A,33A,34A,36A,37A,39A,40A,43A,67A,71A,74A,75A,77A,78A,89A,91A,92A,161A,162
A,163A,165A,166A,169A,189A,202A,229A,232A"/> ***
        <Index name="A_dockable"
resnums="3A,5A,6A,9A,11A,14A,87A,89A,95A,91A,92A,168A,229A,232A,236A,239A,240A,242A,283A" />
                <Index name="B_dockable" resnums="15B,26B,28B,29B,30B,31B,37B,39B,44B,45B,46B"/>
        Index name="B_addl_designable" resnums="" />
        Index name="peptide" resnums="1C,2C,3C,4C,5C,6C,7C,8C,9C,10C,11C,12C"/>

        <Index name="pep_binder_term" resnums="57B"/> This and next to select termini of domains, for distance constraint
        <Index name="enzyme_term" resnums="1A"/>
        <Index name="active_site_pointer" resnums="196A"/> To make sure peptide stays on top of the domain pointing at the active
site, distance constraint
        <Index name="pep_c_term" resnums="10C"/>
        <Index name="pep_binding_resi" resnums="32B,48B"/>
                Index name="res_91A" resnums="91A"/>
        Index name="low_sheet_res" resnums="14B"/>
        Index name="low_helix_res" resnums="110A"/>

                <Or name="A_or_B_dockable" selectors="A_dockable,A_designable,B_dockable"/>
        <Or name="A_or_B_designable" selectors="A_designable,B_dockable"/>
        <InterfaceByVector name="interface"> Selects residues that are in contact at interface
            <Chain chains="A"/> This is somehow sensitive to the order of A and B
            <Chain chains="B"/>
        </InterfaceByVector>

        <Not name="not_designable" selector="A_or_B_designable"/>
        <Not name="not_interface" selector="interface"/>
```

```
            <And name="dockable_interface_residues" selectors="interface,A_or_B_dockable"/>
        <And name="designable_interface_residues" selectors="interface,A_or_B_designable"/> Intersection: residues at the interface AND listed as designable
        <And name="undesignable_unpackable_residues" selectors="not_interface,pep_binding_resi"/>
            <InterfaceByVector name="B_touching_peptide" grp1_selector="peptide" grp2_selector="enzyme"/> Contacting residues between peptide and COMT
    </RESIDUE_SELECTORS>

    <TASKOPERATIONS>
        <OperateOnResidueSubset name="no_design_to" selector="not_designable" >
            <RestrictToRepackingRLT/> Turn off design (allows repacking)
        </OperateOnResidueSubset>

        <OperateOnResidueSubset name="no_pack_design_to" selector="undesignable_unpackable_residues" >
            <PreventRepackingRLT/> Turn off design and repacking (still minimizes)
        </OperateOnResidueSubset>

        <OperateOnResidueSubset name="designable_to" selector="A_or_B_designable">
            <RestrictAbsentCanonicalAASRLT aas="ADEFGHIKLMNPQRSTVWY"/> Don't include CYS during designs
        </OperateOnResidueSubset>

        <InitializeFromCommandline name="ifcl_to" />
        <ExtraRotamersGeneric name="extra_chi" ex1="1" ex2="1" />
    </TASKOPERATIONS>

    <FILTERS>
        DOCKING
            <ResidueCount name="f_dockable_interface_residues" max_residue_count="9999" min_residue_count="0" residue_selector="dockable_interface_residues"/>
        <ResidueCount name="f_designable_interface_residues" max_residue_count="9999" min_residue_count="0" residue_selector="designable_interface_residues"/> SET THIS NUMBER
        <ResidueCount name="f_B_touching_peptide" max_residue_count="9999" min_residue_count="0" residue_selector="B_touching_peptide"/>

        <ScoreType name="distance_constraint_filter" score_type="atom_pair_constraint" scorefxn="low_res_sfxn" threshold="9999"/>

        <ResidueDistance name="termini_distance" res1_pdb_num="1A" res2_pdb_num="57B" distance="9999"/>***
        <ResidueDistance name="pep_active_site_distance" res1_pdb_num="10C" res2_pdb_num="196A" distance="9999"/>***

        DESIGN
        <ScoreType name="f_sfxn_basic" scorefxn="sfxn_basic" threshold="9999"/>
        <ScoreType name="f_sfxn_basic_design" scorefxn="sfxn_basic_design" threshold="9999"/>
        <ReadPoseExtraScoreFilter name="read_dG_over_dSASA" term_name="dG_separated/dSASAx100" threshold="9999"/>
        <ReadPoseExtraScoreFilter name="read_dG_separated" term_name="dG_separated" threshold="9999"/>

        <CalculatorFilter equation="((dG*250) + (Etot))/2" name="f_combined_scores" threshold="99999">
            <Var filter="f_sfxn_basic" name="Etot"/>
            <Var filter="read_dG_over_dSASA" name="dG"/>
        </CalculatorFilter>

        <Sasa name="f_sasa" threshold="0"/> 1300

        <BuriedUnsatHbonds name="new_buns_bb" residue_selector="interface" report_bb_heavy_atom_unsats="true" scorefxn="sfxn_basic" cutoff="99" residue_surface_cutoff="20.0" ignore_surface_res="true" print_out_info_to_pdb="true" use_ddG_style="true" />
        <BuriedUnsatHbonds name="new_buns_sc" residue_selector="interface" report_sc_heavy_atom_unsats="true" scorefxn="sfxn_basic" cutoff="99" residue_surface_cutoff="20.0" ignore_surface_res="true" print_out_info_to_pdb="true" use_ddG_style="true" />

        <CalculatorFilter equation="(buns_bb + buns_sc)" name="f_combined_buns" threshold="9">
            <Var filter="new_buns_bb" name="buns_bb"/>
            <Var filter="new_buns_sc" name="buns_sc"/>
        </CalculatorFilter>
    </FILTERS>

    <MOVERS>
```

```
DOCKING
<DockSetupMover name="dock_setup" partners="A_BC"/>
<DockingInitialPerturbation dock_pert="1" name="dock_set_global" randomize1="0" rot="15" trans="25" />  DAVID
randomize2=1 GLOBAL
<DockingProtocol docking_score_low="low_res_sfxn" low_res_protocol_only="true" name="dock_low_res" partners="A_BC"/>

<DockingInitialPerturbation dock_pert="1" name="dock_set_local" randomize2="1" rot="7" trans="7" />    DAVID randomize2=0
LOCAL

<AddConstraints name="dock_constraint">
  <DistanceConstraintGenerator function="FLAT_HARMONIC 0 1 30" name="terminus_distance_constraint"
residue_selector1="pep_binder_term" residue_selector2="enzyme_term"/> 15
  DistanceConstraintGenerator function="FLAT_HARMONIC 0 1 9" name="terminus_distance_constraint_res91A"
residue_selector1="pep_binder_term" residue_selector2="res_91A"/> 15
  <DistanceConstraintGenerator function="FLAT_HARMONIC 0 1 10" name="peptide_distance_constraint"
residue_selector1="active_site_pointer" residue_selector2="pep_c_term"/> 35
  DistanceConstraintGenerator function="FLAT_HARMONIC 0 1 5" name="lower_sheet_res_cst"
residue_selector1="low_sheet_res" residue_selector2="low_helix_res"/> 15
</AddConstraints>

<ClearConstraintsMover name="clear_constraints"/>

<SwitchResidueTypeSetMover name="cent_to_full" set="fa_standard"/>

AddConstraintsToCurrentConformationMover name="bb_cst" use_distance_cst="0" coord_dev="1.0" bound_width="1"
cst_weight="1.0" residue_selector="receptor" CA_only="1" /> backbone coordinate constraints with 1 angstrom flat-bottom

<SaveAndRetrieveSidechains allsc="1" multi_use="0" name="store_side_chains" two_step="1"/>

FilterReportAsPoseExtraScoresMover name="save_f_contact_A_B" report_as="f_contact_A_B_copy"
filter_name="f_contact_A_B"/>
    <FilterReportAsPoseExtraScoresMover name="save_f_dockable_interface_residues"
report_as="f_designable_interface_residues_copy" filter_name="f_dockable_interface_residues" />
    <FilterReportAsPoseExtraScoresMover name="save_f_designable_interface_residues"
report_as="f_designable_interface_residues_copy" filter_name="f_designable_interface_residues" />
<FilterReportAsPoseExtraScoresMover name="save_f_distance_constraint_filter" report_as="distance_constraint_filter_copy"
filter_name="distance_constraint_filter" />
<FilterReportAsPoseExtraScoresMover name="save_termini_distance" report_as="termini_distance_copy"
filter_name="termini_distance" />
<FilterReportAsPoseExtraScoresMover name="save_pep_active_site_distance" report_as="pep_active_site_distance_copy"
filter_name="pep_active_site_distance" />
<FilterReportAsPoseExtraScoresMover name="save_f_B_touching_peptide" report_as="B_touching_peptide_copy"
filter_name="f_B_touching_peptide" />
<FilterReportAsPoseExtraScoresMover name="save_f_sfxn_basic_design" report_as="f_sfxn_basic_design"
filter_name="f_sfxn_basic_design" />
<FilterReportAsPoseExtraScoresMover name="save_f_sfxn_basic" report_as="f_sfxn_basic" filter_name="f_sfxn_basic" />

DESIGN
<FavorSequenceProfile name="favor_native" scaling="prob" weight="3"  use_starting="1" chain="0" matrix="BLOSUM62"
scorefxns="sfxn_basic_design" />

<FastDesign name="fast_design_1" repeats="1" scorefxn="sfxn_basic_design" relaxscript="KillA2019"
task_operations="no_design_to,no_pack_design_to,designable_to,ifcl_to,extra_chi"/>
<FastDesign name="fast_design_2" repeats="3" scorefxn="sfxn_basic_design" relaxscript="KillA2019"
task_operations="no_design_to,no_pack_design_to,designable_to,ifcl_to,extra_chi"/>

<FilterReportAsPoseExtraScoresMover name="save_f_combined_buns" report_as="f_combined_buns"
filter_name="f_combined_buns"/>
<FilterReportAsPoseExtraScoresMover name="save_f_combined_scores" report_as="f_combined_scores"
filter_name="f_combined_scores"/>

AddConstraintsToCurrentConformationMover name="bb_cst" use_distance_cst="0" coord_dev="1.0" bound_width="1"
cst_weight="1.0" residue_selector="receptor" CA_only="1" /> backbone coordinate constraints with 1 angstrom flat-bottom

<InterfaceAnalyzerMover name="IfaceAnalyzer" scorefxn="sfxn_basic" packstat="1" interface_sc="false" pack_input="false"
pack_separated="1" interface="A_B" tracer="false" />
    FavorSequenceProfile name="favor_native" weight="1" use_native="" matrix="BLOSUM62"/>
    AddCompositionConstraintMover name="percent_composition" filename="70percent_int_composition.comp"/>
```

```
</MOVERS>

<PROTOCOLS>
        protocol is broken up into stages 1-4
  Stage num_runs_per_input_struct="100000" total_num_results_to_keep="10000"> DAVID 100000 10000 GLOBAL
    Add mover="store_side_chains"/>

    Add mover="dock_constraint"/>
    Add mover="dock_setup"/>
    Add mover="dock_set_global"/>
    Add mover="dock_low_res"/>
    Add filter="f_b-site"/> Remove models if lacking a designable residue within 15 of 'receptor-face'

    Add filter="f_dockable_interface_residues"/> Count the number of dockable residues at the interface
    Add mover="save_f_designable_interface_residues"/>
    Add filter="distance_constraint_filter"/>
    Add mover="save_f_distance_constraint_filter"/>

    Add mover="clear_constraints"/>
    Add mover="cent_to_full"/>
    Add mover="store_side_chains"/>

    Sort negative_score_is_good="false" filter="f_dockable_interface_residues"/> Maybe sort by how well it satisfies the
constraints and then sort by number of contacts below
  /Stage>
  <Stage num_runs_per_input_struct="20000" total_num_results_to_keep="10000"> DAVID 10 10000 LOCAL
    <Add mover="store_side_chains"/>

    <Add mover="dock_constraint"/>
    <Add mover="dock_setup"/>
    <Add mover="dock_set_local"/>
    <Add mover="dock_low_res"/>

    <Add filter="f_dockable_interface_residues"/> Count the number of designable residues at the interface
    <Add mover="save_f_dockable_interface_residues"/>
    <Add filter="distance_constraint_filter"/>
    <Add mover="save_f_distance_constraint_filter"/>
    <Add filter="termini_distance"/>
    <Add mover="save_termini_distance"/>
    <Add filter="pep_active_site_distance"/>
    <Add mover="save_pep_active_site_distance"/>
    <Add filter="f_B_touching_peptide"/>
    <Add mover="save_f_B_touching_peptide"/>

    <Add mover="clear_constraints"/>
    <Add mover="cent_to_full"/>
    <Add mover="store_side_chains"/>

    <Sort negative_score_is_good="false" filter="f_dockable_interface_residues"/> Consider sorting by number of contacts here
  </Stage>
  <Stage num_runs_per_input_struct="1" total_num_results_to_keep="2500" > DAVID 1 2500
    <Add mover_name="favor_native" />

    <Add mover_name="fast_design_1"/>

    <Add filter="new_buns_bb"/>
    <Add filter="new_buns_sc"/>
    <Add filter="f_combined_buns"/>

    <Add filter="f_sasa"/>
    <Add mover_name="IfaceAnalyzer"/>

    <Add filter="f_combined_scores"/>
    Add filter="read_dG_cutoff"/> DONT USE THIS
    <Add filter="read_dG_over_dSASA"/>
    <Add filter="read_dG_separated"/>
    <Sort filter="f_combined_scores"/>
  </Stage>
  <Stage num_runs_per_input_struct="1" total_num_results_to_keep="200" > 1 200
```

```
            <Add mover_name="fast_design_2"/>

            <Add filter="f_sfxn_basic_design"/>
            <Add mover_name="save_f_sfxn_basic_design"/> COMPARE DESIGN_NAT_SCORE WITH

            <Add filter="f_sfxn_basic"/>
            <Add mover_name="save_f_sfxn_basic"/> COMPARE DESIGN_NAT_SCORE WITH

            <Add filter="new_buns_bb"/>
            <Add filter="new_buns_sc"/>
            <Add filter="f_combined_buns"/>

            <Add filter="f_sasa"/>
            <Add mover_name="IfaceAnalyzer"/>

            <Add filter="f_combined_scores"/>
            Add filter="read_dG_cutoff"/> DONT USE THIS
            <Add filter="read_dG_over_dSASA"/>
            <Add filter="read_dG_separated"/>

            <Add mover="save_f_combined_scores"/>
            <Add mover="save_f_combined_buns"/>

            <Add filter="f_designable_interface_residues"/>
            <Add mover="save_f_designable_interface_residues"/>
            Add filter="distance_constraint_filter"/> DONT USE THIS
            Add mover="save_f_distance_constraint_filter"/> DONT USE THIS
            <Add filter="termini_distance"/>
            <Add mover="save_termini_distance"/>
            <Add filter="pep_active_site_distance"/>
            <Add mover="save_pep_active_site_distance"/>
            <Add filter="f_B_touching_peptide"/>
            <Add mover="save_f_B_touching_peptide"/>

            NEED ANOTHER SCORE MOVER

            <Sort filter="f_combined_scores"/>
        </Stage>
    </PROTOCOLS>

  </Common>

</JobDefinitionFile>
```

**Linker Design Protocol**

```
<ROSETTASCRIPTS>
  <SCOREFXNS>
    <ScoreFunction name="sfxn_high_res" weights="ref2015_cst"/> Use for fast design
  </SCOREFXNS>

  <RESIDUE_SELECTORS>
    <Index name="linker" resnums="%%RESIDUES%%"/>

    <Not name="not_linker" selector="linker"/>

    <Neighborhood name="neighbor_residues" selector="linker" distance="5"/>

    <Or name="linker_or_neighbors" selectors="linker,neighbor_residues"/>
    <Not name="not_linker_or_neighbor" selector="linker_or_neighbors"/>
  </RESIDUE_SELECTORS>

  <MOVE_MAP_FACTORIES>
    <MoveMapFactory name="movemap" bb="0" chi="0">
      <Backbone residue_selector="not_linker_or_neighbor"/>
      <Chi residue_selector="not_linker_or_neighbor"/>
    </MoveMapFactory>
  </MOVE_MAP_FACTORIES>

  <TASKOPERATIONS>
    <OperateOnResidueSubset name="not_design_TO" selector="not_linker" > Not sure to use this one or next one **
      <PreventRepackingRLT/> Turn off design (allows repacking)
    </OperateOnResidueSubset>

    ReadResfile name="natrot_resfile" filename="RESFILELOCATION" selector="not_neighbors" />

    <OperateOnResidueSubset name="designable_TO" selector="linker"> Can I specify 2??->nope **
      <RestrictAbsentCanonicalAASRLT aas="ADEFGHIKLMNPQRSTVWY"/> Don't include CYS during designs
    </OperateOnResidueSubset>

    <InitializeFromCommandline name="ifcl_TO" />

    <ExtraRotamersGeneric name="extra_chi" ex1="1" ex2="1" />
  </TASKOPERATIONS>

  <FILTERS> I don't think I need a filter section. Maybe rmsd of backbone and closure term
    <ChainBreak name="chain_break_filter" chain_num="1" tolerance=".13" threshold="1"/>

    <ResidueCount name="res_count" />

    <EnergyPerResidue name="max_rama_filter" scorefxn="sfxn_high_res" score_type="rama_prepro"
resnums="%%RESIDUES%%" energy_cutoff=".3"/>
  </FILTERS>

  <MOVERS>
    <FastDesign name="fast_design" repeats="4" scorefxn="sfxn_high_res" relaxscript="KillA2019" movemap_factory="movemap"
task_operations="not_design_TO,designable_TO,ifcl_TO,extra_chi"/>natrot_resfile,

    <RemodelMover name="remod" blueprint="%%BLUEPRINT%%"/>

    <FilterReportAsPoseExtraScoresMover name="report_res_count" report_as="report_res_count" filter_name="res_count"/>
  </MOVERS>

  <PROTOCOLS>
    <Add mover="remod"/>
    <Add filter="chain_break_filter"/>
    <Add mover="fast_design"/>
    <Add filter="max_rama_filter"/>
```

```
    <Add mover="report_res_count" />
  </PROTOCOLS>

  <OUTPUT />
</ROSETTASCRIPTS>
```
